# Vessel Organoids Reveal FOXF1 Variant-Specific Regulation of Mesoderm and Capillary Development

**DOI:** 10.64898/2026.07.12.737936

**Authors:** Nicole M. Pek, Konrad Thorner, Minzhe Guo, Hayden Dennison, Tharushi Rajaguru, George Stan, Keishi Kishimoto, Robbert J. Rottier, Darrell N. Kotton, Aaron M. Zorn, Mingxia Gu

**Affiliations:** Center for Stem Cell and Organoid Medicine (CuSTOM), Cincinnati Children’s Hospital Medical Center (CCHMC), Cincinnati, OH,45229, USA; Division of Pulmonary Biology, CCHMC, Cincinnati, OH, 45229, USA; Division of Developmental Biology, CCHMC, Cincinnati, OH, 45229, USA; College of Medicine, University of Cincinnati, Cincinnati, OH, 45229, USA; Department of Chemistry, University of Cincinnati, Cincinnati, OH, 45221, USA; Department of Cell and Developmental Biology, Erasmus MC, Rotterdam, the Netherlands; Department of Pediatric Surgery, Erasmus MC-Sophia Children’s Hospital, Rotterdam, the Netherlands; Center for Regenerative Medicine, Boston University and Boston Medical Center, Boston, MA, 02118, USA; The Pulmonary Center and Department of Medicine, Boston University School of Medicine, Boston, MA, 02118, USA; Department of Anesthesiology and Perioperative Medicine, David Geffen School of Medicine, University of California, Los Angeles (UCLA), Los Angeles, CA, 90095, USA; Eli and Edythe Broad Center for Regenerative Medicine and Stem Cell Biology, UCLA, Los Angeles, CA, 90095, USA

**Author notes:** Correspondence (N.P.). Correspondence (M.Gu) and (A.Z.). Laboratory for Lung Development and Regeneration, RIKEN Center for Biosystems Dynamics Research (BDR), Kobe, 650-0047, Japan. Senior author (Aaron M. Zorn, Mingxia Gu).

**Keywords:** FOXF1, allelic variants, hiPSCs, vessel organoids, mesoderm, capillaries, endothelial cells, pericytes, endothelial progenitor cells, mural progenitor cells, lipid nanoparticles, mRNA therapy

## Abstract

How allelic variants in lineage-regulating transcription factors drive diverging human developmental outcomes remains poorly understood. This is partly due to the lack of human model systems. Here, we used vessel organoids from human induced pluripotent stem cells (hiPSCs) to resolve variant-specific functions of Forkhead Box F1 (FOXF1), a critical regulator of mesoderm and vascular development. Using three patient-derived hiPSC lines harboring unique *FOXF1* variants, we show that heterozygous variants cause capillary maldevelopment of varying severity. Single-nucleus multiomic analysis revealed variant-specific mechanisms – a severe variant impairs differentiation of nascent mesoderm to lateral plate mesoderm and disrupts vascular progenitor specification, while moderate variants permit mesoderm differentiation but rewire vascular progenitor states and function. Restoration of wild-type *FOXF1* via lipid nanoparticle-mediated mRNA delivery rescued capillary formation in a variant- and developmental-stage-dependent manner. Together, these findings demonstrate that different variants disrupt stage-specific FOXF1 functions in human mesoderm-to-vascular development, underscoring the importance of variant-specific therapeutic strategies.

HIGHLIGHTS

- Human vessel organoids reveal variant-specific roles of *FOXF1* in mesoderm patterning and capillary development.
- Severe *FOXF1* variant c.253T>A (p.F85I) impairs nascent mesoderm-to-lateral plate mesoderm differentiation and disrupts vascular progenitor specification.
- ‘Moderate’ *FOXF1* variants differentially rewire endothelial and mural progenitor cell states and function.
- Lipid nanoparticle-mediated *FOXF1* mRNA delivery rescues capillary formation in a variant- and developmental-stage-dependent manner.

## INTRODUCTION

How distinct allelic variants of lineage-regulating transcription factors (TF) give rise to diverging human developmental outcomes and disease remains a long-standing question in developmental biology. TFs are often studied through complete loss-of-function (LOF) animal models. However, heterozygous missense variants of genes encoding TFs, which may differentially perturb protein function, are often found in human patients. Associating these variants to their effects on cell fate decisions and developmental trajectories in humans remains difficult since access to early developmental stages pose technical and ethical challenges.

Forkhead Box F1 (FOXF1) is a clinically important TF that is well-suited to address this question. FOXF1 is a critical regulator of mesoderm and vascular development^1^ belonging to the Forkhead family of TFs, with an evolutionarily conserved winged helix DNA-binding domain. FOXF1 plays broad roles in development, tissue homeostasis, and disease^2^. Early LOF studies in mice established the importance of FOXF1 in early mesoderm and vascular development - complete loss of Foxf1 in mice is embryonic lethal due to severe defects in the extra-embryonic and lateral plate mesoderm^3^, while heterozygous null mice exhibited early vascular abnormalities, including the absence of vasculature in the yolk sac and allantois^3^. *Foxf1^−/+^* mice also exhibited severe pulmonary defects, including loss of lung vasculature, defects in alveolarization, and fusion of lung lobules^4,5^. While informative, these LOF models yield relatively uniform phenotypes that reflect gene-level haploinsufficiency and do not accurately recapitulate variability across human patients, thereby limiting insights into how specific *FOXF1* variants differentially affect mesoderm differentiation and vascular lineage specification.

More than 50 allelic variants of *FOXF1* have been identified in patients with Alveolar Capillary Dysplasia with Misalignment of Pulmonary Veins (ACDMPV), a rare and lethal congenital lung disorder^6,7^. These variants include missense variants and large genomic copy number variant (CNV) deletions at 16q24.1, which contains the *FOXF1* locus^7,8^. ACDMPV is characterized by profound loss of alveolar capillaries^9^. More than 80% of ACDMPV patients also present extra-pulmonary features such as esophageal atresia, intestinal malrotation, and even urogenital abnormalities^9^. No effective therapies presently exist for the treatment of ACDMPV^9^. Notably, ACDMPV patients exhibit variability in disease severity and clinical presentation, suggesting that distinct *FOXF1* variants differentially perturb developmental programs^10–13^.

To date, only one pathogenic *FOXF1* missense variant, c.155C<T (p.S52F), has been modeled and characterized in mice. Remarkably, Foxf1^WT/S52F^ mice recapitulated certain clinical features of ACDMPV, including lobular underdevelopment and loss of distal pulmonary vasculature^14,15^. S52F FOXF1 was found to be transcriptionally inactive, resulting in FOXF1 haploinsufficiency. However, the c.155C<T *FOXF1* variant certainly does not represent the full spectrum of *FOXF1* allelic variants. Therefore, the molecular and functional consequences of other variants remain largely unknown. A recent study showed that different ACDMPV-causing *FOXF1* missense variants have highly-varied effects on FOXF1 function^16^. How these effects subsequently impact mesoderm and vascular cell fate specification in humans remains unclear. This knowledge gap stems, in part, from the lack of patient-specific human models that enable temporal study of how *FOXF1* variants affect key stages of mesoderm-to-vascular development. Addressing this is essential for connecting genotype to developmental consequences and guiding precise, variant-informed therapeutic strategies.

Here, we utilized human vessel organoids (VOs) from human induced pluripotent stem cells (hiPSCs) derived from three different ACDMPV patients carrying distinct *FOXF1* allelic variants: one with a large genomic CNV variant at 16q24.1, spanning the *FOXF1* locus (16q24.1del)^17,18^, and two with unique heterozygous *FOXF1* missense variants c.166C>G (p.L56V) and c.253T>A (p.F85I)^16,19^. Both residues 56 and 85 lie within the DNA-binding domain (DBD) of FOXF1^16^. We demonstrated that different *FOXF1* allelic variants result in capillary maldevelopment across a spectrum of severity, with the F85I FOXF1 variant producing the most profound developmental defect. By employing single-nucleus multiomic analysis, we uncovered variant-specific mechanisms underlying these phenotypes. ‘Severe’ F85I FOXF1 variant perturbs early mesodermal differentiation, disrupts FOXF1-MECOM regulatory axis in endothelial progenitors, and reduces the population of collagen-producing mural progenitors. Conversely, the other two ‘moderate’ variants, L56V FOXF1 and 16q24.1del, permit proper mesoderm differentiation, but rewire endothelial and mural progenitor transcriptional programs that stifled proper capillary formation. Lastly, we showed that lipid nanoparticle-mediated delivery of wild-type *FOXF1* mRNA (LNP-*FOXF1*) rescues capillary defects in a variant- and developmental-stage-dependent manner. Together, these findings define the unique consequences of *FOXF1* variants on human vascular development at a single-cell resolution and highlight the importance of considering variant specificity during therapeutic interventions.

## RESULTS

### Patient-specific FOXF1 allelic variants cause capillary maldevelopment of varying severity

To elucidate the patient-specific effects of *FOXF1* variants, we differentiated three hiPSC lines generated from ACDMPV patients (ACD-iPSCs: ACD1, ACD2, and ACD3), each harboring a unique *FOXF1* variant (**Fig. 1A, S1A**, and **Table S1**), to VOs (ACD-VOs) using an established protocol^20,21^ (**Fig. S1B**). ACD1 (clone FOXF1.1) patient has a heterozygous 1.7Mb deletion mutation of the chr16:85,063,102-86,721,852 region, which spans the *FOXF1* locus^17,18^. ACD2 and ACD3 patients carry heterozygous missense variants c.166C>G and c.253T>A respectively^19^ (**Fig. 1A** and **S1A**). Isogenic hiPSC controls for ACD2 and ACD3 (named iACD2 and iACD3, respectively) were generated by correcting the point mutations via CRISPR-Cas9 genome editing (**Fig. S1A** and **S1C**). Lastly, an unrelated healthy, normal donor hiPSC line was used as a control for ACD1 (**Fig. S1A**). VO differentiation from hiPSCs closely follows the developmental trajectory of capillaries in vivo, with day 3, day 5, day 10, and day 15 corresponding to the lateral plate mesoderm (LPM), vascular progenitor specification (emergence of endothelial and mural progenitors), vascular sprouting, and finally vascular remodeling and maturation stages respectively (**Fig. S1D** and **S1E**). At day 15, VOs consist of robust, well-organized capillary networks composed of endothelial cells (ECs), supported by mural cells such as pericytes (**Fig. S1D** and **S1F**).

**Figure 1.**
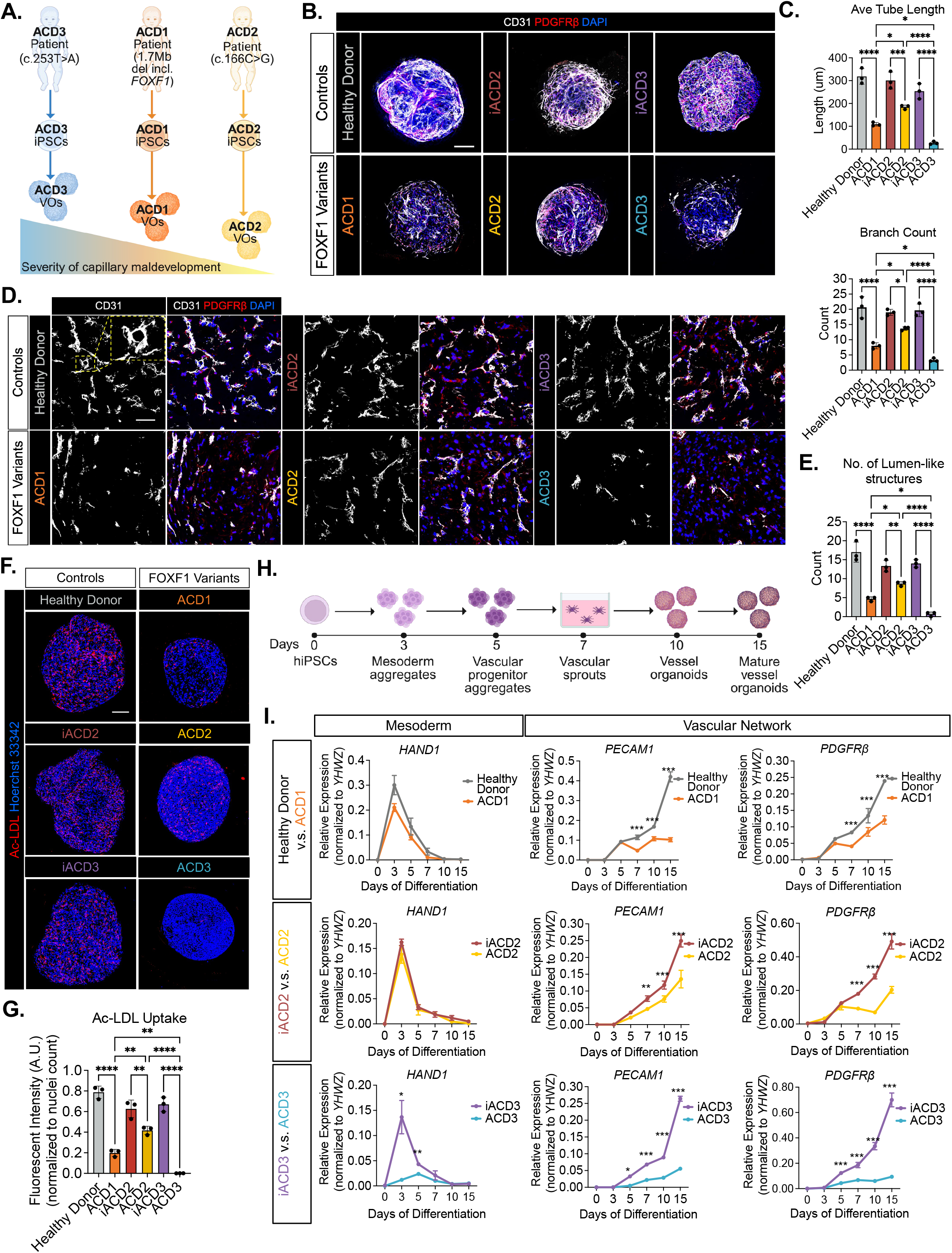
Patient-specific *FOXF1* allelic variants cause capillary maldevelopment of varying severity. (A) Schematic illustrating the varied severity of capillary maldevelopment caused by different human *FOXF1* allelic variants. (B) Representative whole-organoid immunofluorescence images showing CD31 (white), PDGFRβ (red) in D15 VOs. Nuclei are counterstained with DAPI (blue). Scale bar represents 100um. (C) Quantification of vessel parameters (mean ± S.D., n=3 organoids from 3 biological replicates, *p<0.05 ***p<0.001 ****p<0.0001 by one-way ANOVA followed by Tukey’s test). (D) Representative immunofluorescence images of D15 VO cryosections showing CD31 (white) and PDGFRβ (red). Nuclei are counterstained with DAPI (blue). Scale bar=100um. (E) Quantification of lumen-like structures (mean ± SD, n=3 organoids from 3 biological replicates, *p<0.05, **p<0.01, ****p<0.0001 by one-way ANOVA followed by Tukey’s test). (F). Representative whole-organoid live immunofluorescence images showing Ac-LDL uptake (red). Nuclei are counterstained with Hoerchst 33342 (blue). Scale bar=100um. (G) Quantification of Ac-LDL uptake (mean ± S.D., n=3 organoids, **p<0.01, ****p<0.0001 by one-way ANOVA followed by Tukey’s test). (H) Schematic illustrating the key timepoints of VO differentiation. (I) Timecourse mRNA expression profile of mesoderm marker *HAND1* and vascular markers *PECAM1* (endothelial) and *PDGFRβ* (pericyte) across VO differentiation (mean ± S.D., n= 3 biological replicates **p<0.01, ****p<0.0001 by two-way ANOVA followed by Bonferroni’s test).

We observed impaired capillary development of varying severity in ACD-VOs compared to the control-VOs as marked by endothelial marker CD31 and pericyte marker PDGFRβ, with ACD3-VOs being the most underdeveloped, followed by ACD1- and ACD2-VOs (**Fig. 1A-1C**). ACD-VOs had shorter vessels with less branching and significantly fewer endothelial cell (ECs) lined lumen-like structures (**Fig. 1D** and **1E**). Capillary function, as measured by EC uptake of acetylated low-density lipoprotein (Ac-LDL), part of the LDL scavenger pathway^22^, was also significantly lower in ACD3-VOs compared to controls, followed by ACD1-VOs, and then ACD2-VOs (**Fig. 1F** and **1G**). Consistent with reduced capillary lumens ACD-VOs also exhibited increased hypoxia (**Fig. S1G** and **S1H**). This finding coincides with increased hypoxia-inducible factor 1α signaling in ACDMPV patient samples^23^. Collectively, these data illustrate that different *FOXF1* allelic variants lead to varying severity of capillary maldevelopment and functions.

To define the onset of *FOXF1* variant-driven pathogenesis, the expression of mesoderm marker *HAND1* and vascular markers *PECAM1* and *PDGFRβ* was assessed throughout VO differentiation (**Fig. 1H**). At day 3, when *HAND1* levels normally peak and vascular markers begin to increase in control VOs, *HAND1* expression was significantly reduced in ACD3 relative to iACD3 (**Fig. 1I**), indicating impaired mesoderm development. This defect, however, was not observed in ACD1 and ACD2 (**Fig. 1I**), but we found that these did exhibit decreased vascular marker expression by day 5 in ACD3, and by day 7 (**Fig. 1I**).

Collectively, these findings demonstrate *FOXF1* variant-specific, developmental-stage-dependent disruptions in capillary development, where some variants stifle early mesoderm differentiation, while others primarily affect later vascular specification, revealing the temporal origins of capillary defects.

### Molecular outcomes of FOXF1 variants are distinct from FOXF1 LOF models

FOXF1 haploinsufficiency (HI) has been suggested to be the main mechanism by which *FOXF1* variants cause ACDMPV^7^. To understand if the *FOXF1* allelic variants in ACD1, ACD2, and ACD3 result in FOXF1 HI, FOXF1^−/+^ (HI), and FOXF1^−/−^ (null) LOF hESC lines were generated (DBD indel mutations) and differentiated into VOs. (**Fig. 2A**). As expected, a genotype-dependent reduction of FOXF1 protein levels was observed, with ∼50% loss in FOXF1^−/+^ and a complete loss in FOXF1^−/−^ in day 3 and day 5 VOs, which correspond to mesoderm and vascular progenitor stages of capillary development (**Fig. 2B** and **C**). We observed a dose-dependent reduction in FLK1 protein expression (also known as VEGFR2/ KDR), a known downstream target of FOXF1^24^, in FOXF1-LOF VOs consistently at days 3 and 5 (**Fig. 2B** and **C**), which also showed a significant loss of capillary networks and lumen-like structures at day 15 (**Fig. 2D-2F**). Despite the complete loss of FOXF1 expression in day 5 FOXF1^−/−^-VOs, FLK1 expression remained detectable, likely due to VEGFA-induced^25^ FOXF1-independent regulation (**Fig. 2B** and **C**).

**Figure 2.**
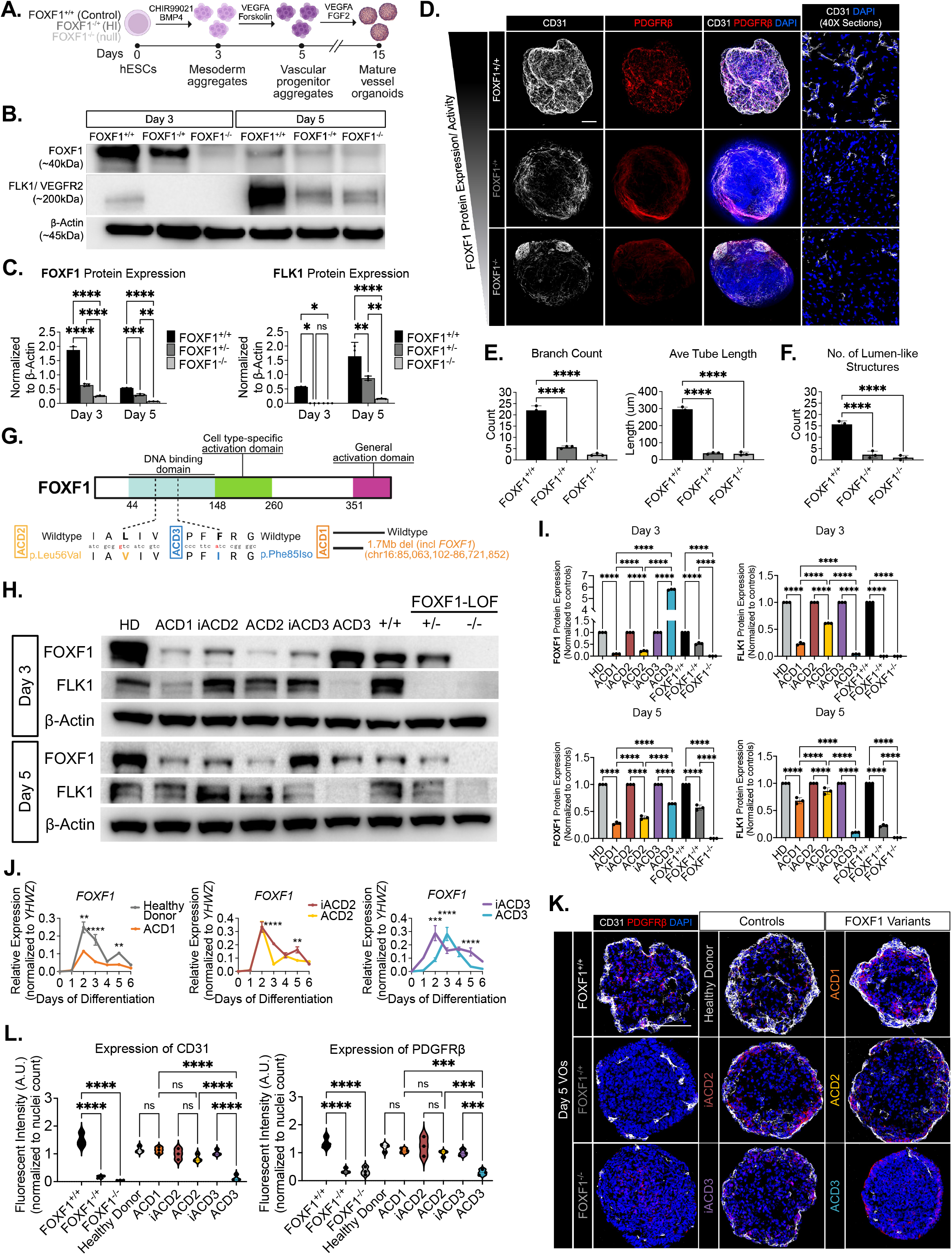
Different *FOXF1* variants elicit unique molecular outcomes that are distinct from FOXF1 LOF. (A) Schematic illustrating the differentiation of FOXF1 LOF hESC lines (FOXF1^−/+^, FOXF1^−/−^) and the respective control (FOXF1^+/+^). (B) Representative western blot image showing FOXF1, FLK1, and B-ACTIN. (C) Quantification of FOXF1 and FLK1 protein expression (mean ± SD, n=3 biological repeats, *p<0.05, **p<0.01, ***p<0.001, ****p<0.0001 by two-way ANOVA followed by Bonferroni’s test). (D) Whole-organoid images: Representative whole-organoid immunofluorescence images showing CD31 (white), PDGFRβ (red). Nuclei are counterstained with DAPI (blue). Scale bar represents 100um. Immunostained sections: Representative immunofluorescence images of D15 VO cryosections showing CD31 (white) and PDGFRβ (red). Nuclei are counterstained with DAPI (blue). Scale bar=100um. (E) Quantification of vessel parameters (mean ± SD, n=3 organoids from 3 biological replicates, ****p<0.0001). (F) Quantification of lumen-like structures (mean ± SD, n=5 organoids, ****p<0.0001, n=3 organoids from 3 biological replicates). (G) Schematic illustrating the *FOXF1* allelic variants in the three patient-derived iPSC lines ACD1, ACD2, and ACD3. (H) Representative western blot image showing FOXF1 and FLK1 protein expression in day 3 and 5 VOs across all patient-derived and FOXF1 LOF lines (including their respective controls). (I) Quantification of FOXF1 and FLK1 protein expression. Protein expression was normalized to loading control β-Actin and protein expression in ACD1, ACD2, ACD3, FOXF1 ^−/+^, and FOXF1^−/−^ were normalized to their respective controls (mean ± SD, n=3 biological repeats, *p<0.05, **p<0.01, ***p<0.001, ****p<0.0001 by two-way ANOVA followed by Bonferroni’s test). (J) Timecourse mRNA expression profile of *FOXF1* across the early stages of VO differentiation (mean ± S.D., n= 3 biological replicates **p<0.01, ***p<0.001, ****p<0.0001 by two-way ANOVA followed by Bonferroni’s test). (K) Representative immunofluorescence images of D5 VO cryosections showing CD31 (white) and PDGFRβ (red). Nuclei are counterstained with DAPI (blue). Scale bar=100um (L) Quantification of CD31 and PDGFRβ fluorescent intensity normalized to nuclei count (mean ± SD, n=3 organoids from 3 biological replicates, ***p<0.001, ****p<0.0001 by one-way ANOVA followed by Tukey’s test).

Next, we directly compared VOs from FOXF1-LOF to patient lines. The *FOXF1* missense variants in ACD2 and ACD3 correspond to p.L56V and p.F85I substitutions, respectively (**Fig. 2G**), both located in the DBD of FOXF1 (**Fig. S2A**). FOXF1 and FLK1 protein expression were assessed on days 3 and 5 (**Fig. 2H** and **2I**). Protein expression levels of ACD1, ACD2, ACD3, and FOXF1-LOF were normalized to their respective controls to correct for variability among control samples, which likely reflects differences in genetic background. At days 3 and 5, FOXF1 expression is significantly reduced in ACD1 and ACD2 compared to their respective controls, with FLK1 showing a similar pattern (**Fig. 2H** and **2I**). Unlike in the FOXF1-LOF, FLK1 expression, although reduced in ACD1 and ACD2 relative to their respective controls, was still preserved (**Fig. 2H** and **2I**). This suggests that FOXF1 variant proteins in ACD1 and ACD2 retained some function. Surprisingly, FOXF1 expression in ACD3 at day 3 is markedly increased compared to iACD3 (**Fig. 2H** and **2I**), but then by day 5 was significantly lower than in iACD3 (**Fig. 2H** and **2I**). Despite FOXF1 being expressed in ACD3 at both timepoints, FLK1 expression was almost completely abolished, suggesting a loss of protein function (**Fig. 2H** and **2I**). Altogether, FOXF1 protein expression and function vary substantially between *FOXF1* variants and differ from FOXF1-LOF.

We further examined dynamic *FOXF1* mRNA expression over the first 6 days of VO differentiation (**Fig. 2J**). In controls, *FOXF1* mRNA levels peaked at day 2, then gradually declined overtime before increasing again at day 5 (**Fig. 2J**), reflecting its roles in both mesoderm and vascular development stages. *FOXF1* expression was significantly reduced in both ACD1 and ACD2 at both day 3 and day 5 when compared to controls (**Fig. 2J**), consistent with protein expression (**Fig. 2H** and **2I**). Interestingly, in ACD3, *FOXF1* expression appeared to exhibit a temporal shift, peaking on day 3 rather than day 2 as observed in iACD3 and the other cell lines, which could explain the unexpected upregulation of FOXF1 protein observed at day 3 (**Fig. 2H** and **2I**). Moreover, in ACD3, we did not observe the second peak of expression on day 5 as we did in other lines (**Fig. 2H** and **2I**). This observation emphasizes the importance of a temporal analysis as distinct *FOXF1* allelic variants appear to differentially alter the dynamics of *FOXF1* expression.

To evaluate the impact of this altered FOXF1 expression profile on early vascular lineage specification, we compared day 5 VOs from FOXF1-LOF to patient lines. FOXF1-LOF VOs exhibited a dramatic loss of both endothelial progenitor cell (EPC) and mural progenitor cell (MPC) populations at day 5 (**Fig. 2K** and **2L**). Likewise, a significant loss of vascular progenitors was also observed in day 5 ACD3-VOs (**Fig. 2K** and **2L**). However, EPC and MPC populations remain relatively unchanged in ACD1 and ACD2-VOs (**Fig. 2K** and **2L**), further illustrating *FOXF1* variant-specific effects on vascular development.

Taken together, these findings demonstrate that *FOXF1* variants do not all act through a simple haploinsufficiency mechanism. Instead, they exert complex, variant-specific, and temporally dynamic effects during human development that are not fully captured by FOXF1 LOF mouse models.

### Differential DNA-binding and flexibility prediction profiles of F85I and L56V FOXF1

To further dissect how different variants contribute to changes in FOXF1 protein structure and DNA binding profiles, the effects of F85I and L56V amino acid substitutions were computationally modeled (**Fig. 3A**). Computational tools AlphaMissense^26^ and PolyPhen^27^ predicted that both F85I and L56V FOXF1 likely have compromised protein functions (**Fig. 3B**). To determine the biophysical properties underlying these predictions, molecular dynamics simulations of WT, F85I, and L56V FOXF1 were performed. Unbound/ APO F85I FOXF1 was significantly more flexible compared to WT or L56V FOXF1 (**Fig. 3C** and **3D**). When bound to double-stranded DNA (dsDNA) (bound/COMPLEX), the flexibility of L56V FOXF1 appears to be higher compared to WT and F85I FOXF1 (**Fig. 3C** and **S2B**). The core region of the protein became more flexible in the native state upon the introduction of F85I and L56V amino acid substitutions. This increased flexibility persists in the dsDNA-bound state, with the F85I variant showing a more pronounced effect in that region (**Fig. 3E** and **3F**). Interestingly, the F85I variant resulted in weaker protein-DNA interactions, as evidenced by the loss of hydrogen bonds and salt bridges (**Fig. 3G** and **3H**). By contrast, the L56V variant had a mild effect on protein-DNA interactions, with the same number of hydrogen bonds and salt bridges formed between the L56V FOXF1 and dsDNA, comparable to WT FOXF1 and dsDNA (**Fig. 3G** and **3H**). The total solvent accessibility area (SASA) for each of the WT, L56V, and F85I FOXF1 protein simulations exhibited a bimodal Gaussian distribution, with L56V FOXF1 emphasizing a larger SASA peak, while F85I and WT FOXF1 emphasizing a smaller SASA peak, suggesting altered flexibility of L56V FOXF1 (**Fig. 3I**). These shifts in the number of salt bridges, hydrogen bonds, and SASA suggest different DNA-binding profiles for each variant compared to WT FOXF1 and can be attributed to changes in the dynamics of the Wing 2 region of the FOXF1 protein binds dsDNA (**Fig. 3J** and **S2C**). Therefore, our predictions suggest that L56V preserves FOXF1 DNA-binding capacity but alters protein flexibility, whereas F85I disrupts both DNA binding and protein flexibility, which could explain the differential molecular and phenotypic outcomes of L56V and F85I FOXF1.

**Figure 3.**
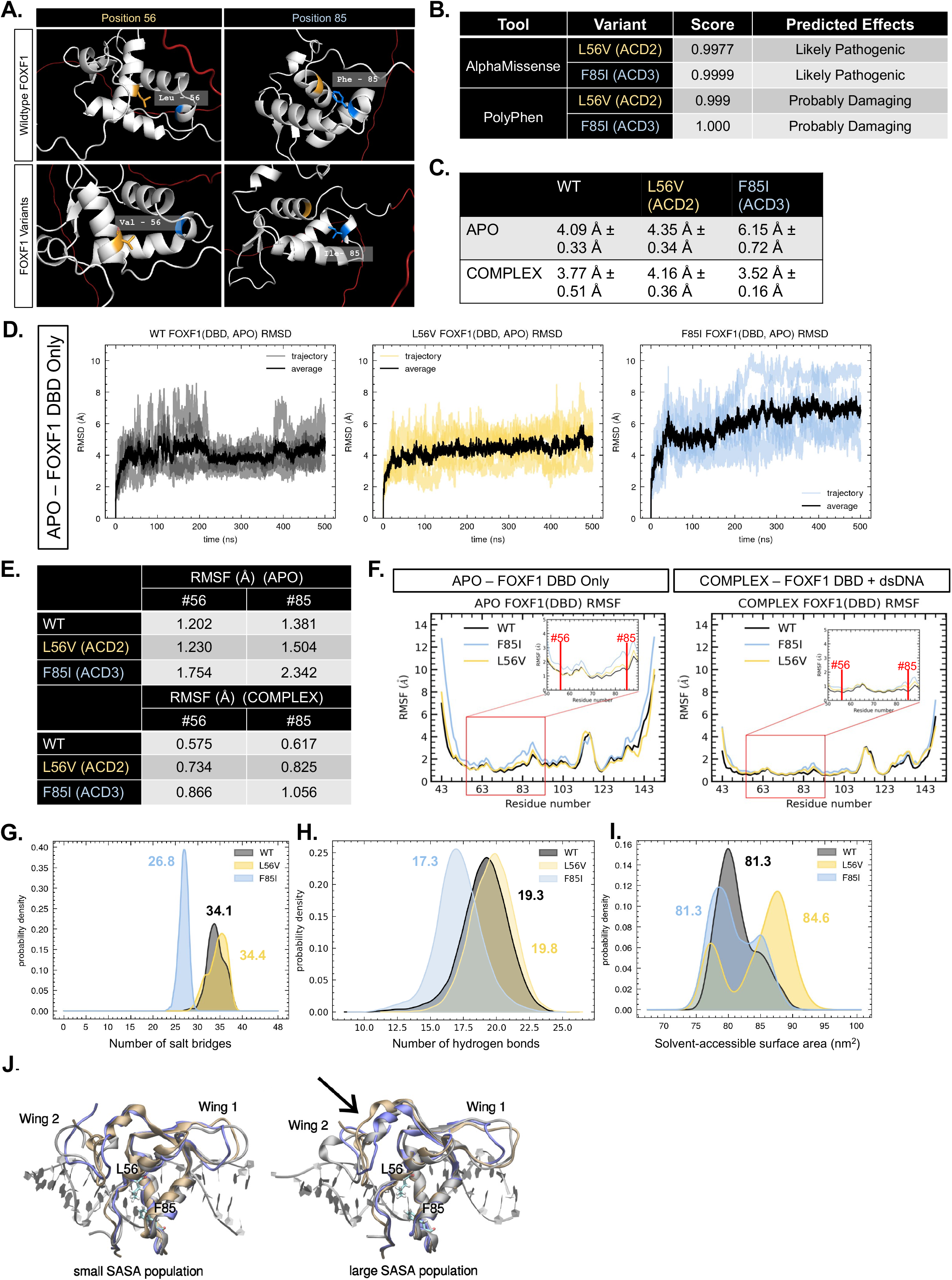
Computational analysis of DNA-binding and flexibility profiles of L56V and F85I FOXF1. (A) AlphaFold model of FOXF1 with 56^th^ and 85^th^ amino acid residue positions denoted in yellow and blue colors, respectively. (B) Predicted effects of missense variants using AlphaMissense and PolyPhen. (C) Average RMSD (Å) values (± SEM) of unbound/ APO and bound/ COMPLEX WT, L56V, and F85I FOXF1 proteins (D) RMSD (Å) plot of APO/unbound WT, L56V, and F85I FOXF1 proteins over simulation time of 500ns. (E) Average RMSF (Å) values at 56^th^ and 85^th^ positions of unbound/ APO and bound/ COMPLEX WT, L56V, and F85I FOXF1 proteins. (F) RMSF (Å) plot of APO/unbound and bound/ COMPLEX WT, L56V, and F85I FOXF1 proteins across individual amino acid residues. G) Gaussian kernel density estimation (KDE) plots of the average number of salt bridges between WT, L56V, and F85I FOXF1 proteins and dsDNA across trajectories. (H) KDE plots of the average number of hydrogen bonds between WT, L56V, and F85I FOXF1 proteins and dsDNA across trajectories. (I) KDE plots of the average solvent-accessible surface area between WT, L56V, and F85I FOXF1 proteins and dsDNA across trajectories. (J) Representative structures of WT (grey), L56V (ochre), and F85I (blue) FOXF1 proteins with low (left) and high (right) total solvent-accessible surface area. The structures for L56V and F85I FOXF1 are best fit to the corresponding WT structure. The arrow points to the Wing 2 region in the large SASA conformation that displays the largest difference in relative solvent accessibility from the small SASA population.

### Single-nucleus multiomic analysis revealed early severe developmental abnormalities in ACD3

To identify cellular and molecular mechanisms underlying the phenotypic variability observed in ACD-VOs, single-nucleus multiome sequencing (snMultiome-seq) was performed on control and ACD-VOs at both day 3 (mesoderm) and day 5 (vascular progenitor specification) stages, the two developmental peaks in *FOXF1* expression (**Fig. S3A**). Day 3 VOs (n=57043) consist of cells identified as lateral plate mesoderm (LPM), nascent mesoderm (NasM), neuroectoderm (NE), and epithelial (Epi) cells (**Fig. 4A**). LPM and NasM annotations were benchmarked against published human fetal and hiPSC-2D gastruloids datasets^28,29^ (**Fig. 4B**), and annotations of NE and Epi cells were supported by marker gene expression (**Fig. S3B**) and Gene Ontology (GO) enrichment (**Fig. S3C**). Day 3 cells are mostly void of definitive endoderm, paraxial mesoderm, pluripotent, or primitive streak cells (**Fig. S3D**). In day 5 VOs (n=46367), three main cell clusters were identified – endothelial progenitor cells (EPC), mural progenitor cells (MPC), and neuroectodermal progenitor cells (NEP) (**Fig. 4A** and **S3E**). GO analysis of enriched differentially expressed genes (DEGs) (BP: Biological Process) in ACD3-NEP showed terms associated with neuronal identity (**Fig. S3F**).

**Figure 4.**
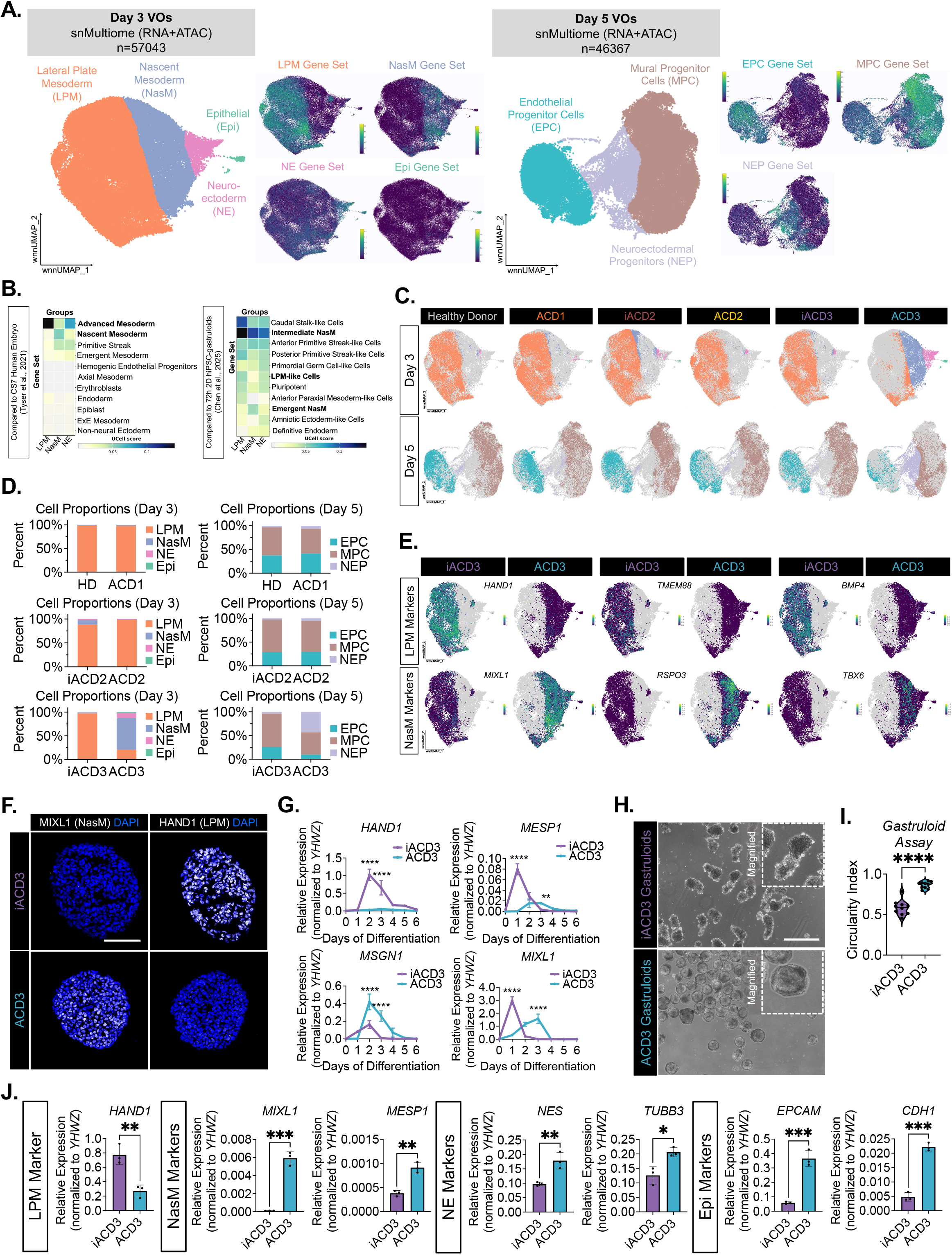
Single nucleus-multiomic analysis revealed developmental abnormalities unique to ACD3-VOs. (A) Left: Uniform manifold approximation and projection (UMAP) visualization of D3 (n=57043) and D5 (n=46367) VOs. Right: UMAP visualization of gene set module scores for each cell type annotation. (B) Heatmap comparing D3 ‘NasM’ and ‘LPM’ populations to cell annotations from Tyser et al., 2021 and Chen et al., 2025’s datasets. (C) UMAP visualization of D3 and D5 VOs across all cell lines. (D) Cell proportion plots of D3 and D5 VOs across all cell lines. (E) Feature plots showing expression of LPM and NasM markers in D3 iACD3 and ACD3-VOs. (F) Representative immunofluorescence images of D3 VO cryosections showing MIXL1 and HAND1 expression. Nuclei are counterstained with DAPI (blue). Scale bar=100um. (G) Timecourse mRNA expression profile of *HAND1, MESP1, MIXL1,* and *MSGN1* across the early stages of VO differentiation (mean ± S.D., n= 3 biological replicates **p<0.01, ****p<0.0001 by two-way ANOVA followed by Bonferroni’s test). (H) Representative brightfield images of ACD3 and iACD3-gastruloids. Scale bar=500um. (I) Quantification of gastruloid circularity (mean ± SD, n=10 organoids from 3 biological replicates, ****p<0.0001 by paired student’s t-test). (J). mRNA expression of LPM, NasM, NE, and Epi markers in ACD3 and iACD3-gastruloids (mean ± SD, n= 3 biological replicates, *p<0.05, **p<0.01, ***p<0.001 by paired student’s t-test).

Interestingly, the most drastic cell population changes were observed in ACD3-VOs – at day 3, there is a significant loss of LPM cells but an expansion of NasM, NE, and Epi cell populations compared to iACD3 (**Fig. 4C-4E** and **S3G**). These population changes, however, are not observed in ACD1 and ACD2 (**Fig. 4C, 4D,** and **3SH**), suggesting that the developmental abnormalities are unique to ACD3 pathology. Consistent with the snMultiome-seq data, bulk qPCR analysis of ACD3-VOs confirmed loss of LPM and enrichment of NasM cells. (**Fig. S3I**). Similarly, FOXF1-LOF VOs showed a significant decrease in LPM marker expression and a dramatic increase in NasM, NE, and Epi gene marker expression (**Fig. S4A**). Immunostaining of MIXL1 (NasM) and HAND1 (LPM) further confirmed those cell population changes in day 3 ACD3-VOs and FOXF1-LOF VOs (**Fig. 4F** and **S4B**), suggesting that ACD3 pathogenesis resembles that of FOXF1-LOF.

At day 5, ACD3-VOs showed a significant loss of both EPC and MPC populations and an expansion of NEP cells compared to iACD3 (**Fig. 4C** and **4D**), which was likely due to insufficient LPM differentiation at day 3. The drastic population changes in ACD3 during mesoderm and vascular development were further confirmed in additional biological replicates (ACD3, n=3; iACD3, n=2) for both day 3 and 5 (**Fig. S3J** and **S3K**). In contrast, the overall population sizes of EPCs and MPCs at day 5 remained relatively unchanged in ACD1 and ACD2 (**Fig. 4C** and **4D**). To further characterize the dynamic changes during NasM and LPM development in ACD3, additional timepoints were examined daily from days 1 to 7 via RT-qPCR. In controls, mRNA expression of NasM markers *MIXL1, MSGN1,* and *MESP1* peaked at day 1, followed by LPM differentiation (*HAND1)* at day 2 (**Fig. 4G**). In ACD3, NasM genes remain highly expressed at day 3 when compared to iACD3 (**Fig. 4G**), indicating delayed NasM differentiation that may have impeded the transition into LPM. Collectively, these data suggest that the c.253T>A *FOXF1* variant (p.F85I) in ACD3 impairs NasM-to-LPM differentiation.

Since the VO differentiation process relies on directed mesoderm differentiation, we sought to further dissect the role of the c.253T>A *FOXF1* variant (p.F85I) in ACD3 within a broader early human developmental context by generating iACD3 and ACD3 gastruloids^30^. iACD3 gastruloids were able to elongate along the anterior-posterior axis by 96h, which was indicative of proper gastrulation. However, ACD3 gastruloids remained largely circular, suggesting gastrulation defects (**Fig. 4H** and **4I**). ACD3 gastruloids mirrored D3 ACD3-VOs, with significant downregulation of LPM marker *HAND1*, while NasM (*MIXL1, MESP1*), NE (*TUBB3, NES*), and Epi (*EPCAM, CDH1*) markers were significantly upregulated (**Fig. 4J**). Additionally, FOXF1-LOF gastruloids also displayed gastrulation defects illustrated by increased circularity (**Fig. S4C** and **S4D**), coupled with a significant loss of *HAND1* expression and elevated expression of NasM, NE, and Epi genes (**Fig. S4E**). Overall, our data suggest that the c.253T>A *FOXF1* variant (p.F85I) severely impairs NasM-to-LPM differentiation and appears to promote the mis-patterning of cells towards other lineages such as NE and Epi cells.

### MECOM is a direct FOXF1 target that can account for endothelial dysfunction in ACD3-EPCs

It is likely that impaired LPM differentiation in ACD3 resulted in diminished proportion of both EPCs and MPCs in day 5 ACD3-VOs (**Fig. 2K, 2L, 4C, 4D, 5A, S5A,** and **S5B**). To examine the role of FOXF1 in vascular cell fate specification, independent of mesoderm differentiation, we compared control and ACD3 EPCs and MPCs at day 5 to assess transcriptomic and functional defects beyond the population loss resulting from impaired LPM differentiation. GO analysis of downregulated DEGs (n=211) (BP) in ACD3-EPCs showed terms associated with endothelial development and function (**Fig. 5B**). Transcripts downregulated in ACD3-EPCs include known FOXF1 targets, *FLK1* and *FLT1*, as well as *TLL1, TFPI,* and *CALCRL* (**Fig. 5C**). *FLT1* is involved in vascular growth and function^31^, while the roles of *TFPI*, *TLL1*, and *CALCRL* in endothelial identity and function remain unclear. Interestingly, the most downregulated transcript in ACD3-EPCs was *MECOM*, which encodes a zinc finger TF known to be essential for endothelial development and function^32,33^ (**Fig. 5C**). To determine if *MECOM* was a direct target of FOXF1, we first examined ATAC-seq data to identify putative enhancers (n=3556) with chromatin accessibility that changed in ACD3-EPCs relative to iACD3-EPCs. Then, we associated these differential ATAC peaks with downregulated transcripts to determine putative direct targets of FOXF1 (n=1566) (**Fig. 5D**). We identified both established FOXF1 target genes, such as *FLK1* and *FLT1*, as well as *MECOM* (p-value=0.016) (**Fig. 5D**).

**Figure 5.**
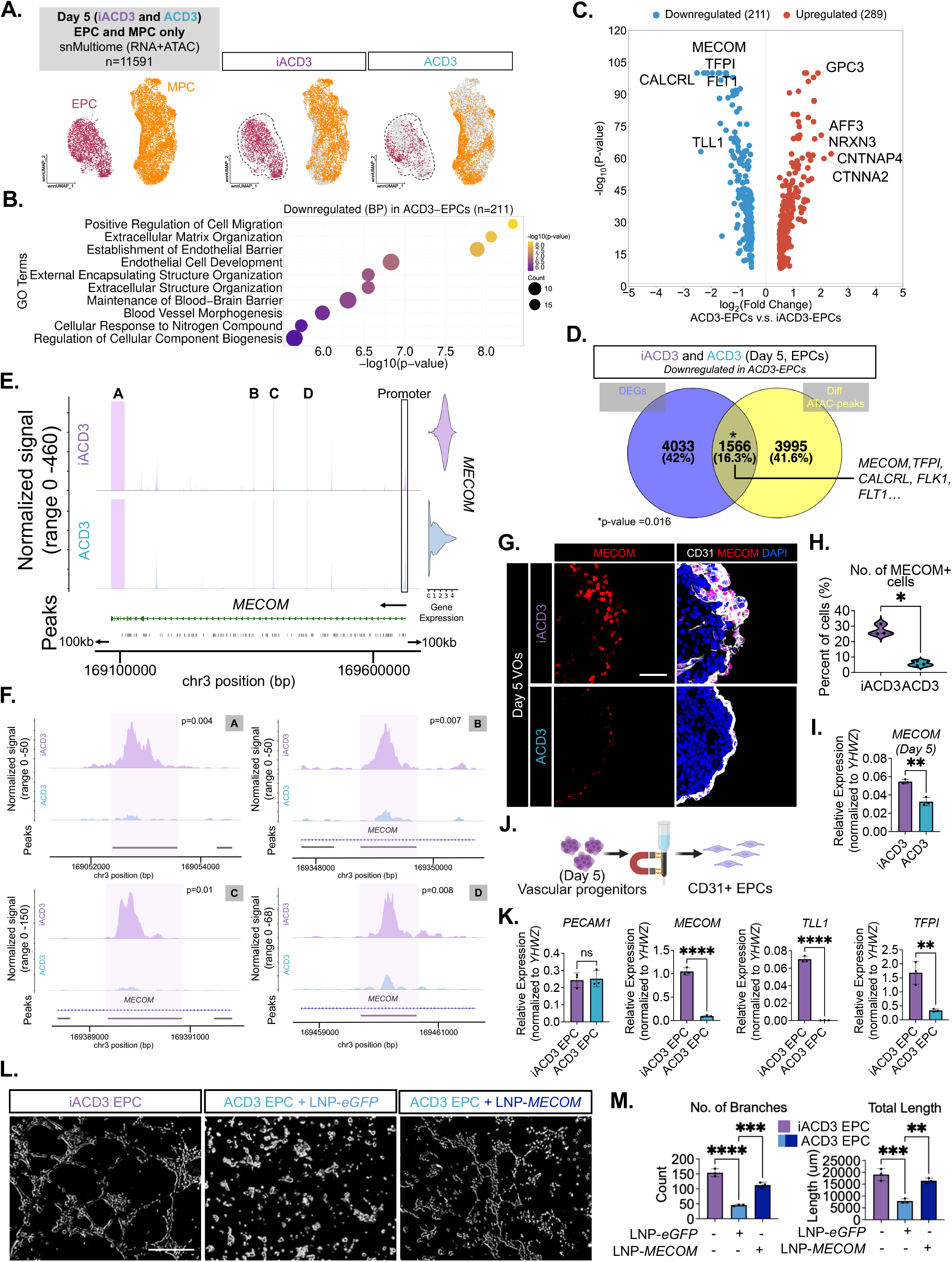
Loss of MECOM expression in ACD3-EPCs is associated with endothelial dysfunction. (A) UMAP visualization of D5 iACD3 and ACD3 VOs, EPCs and MPCs only (n=11591). (B) GO term analysis (BP) of downregulated DEGs in ACD3-EPCs (n=214). (C) Volcano plot showing top DEGs in ACD3-EPCs compared to iACD3-EPCs (log2FC>0.5, p<0.05). (D) Venn diagram showing common genes between downregulated DEGs and downregulated differential ATAC-peaks (log2FC>0.5, p<0.05) in ACD3-EPCs compared to iACD3-EPCs (*p=0.016). DEGs and differential ATAC-peaks were obtained from n=2 iACD3 and n=3 ACD3 samples. (E) Coverage plot showing chromatin accessibility and mRNA expression of *MECOM* in D5 iACD3 and ACD3-VOs. Normalized signal is represented as an average of n=3 for ACD3 and n=2 for iACD3. (F) Normalized signal of four putative FOXF1-binding enhancers of *MECOM*. Normalized signal is represented as an average of n=3 for ACD3 and n=2 for iACD3. (G) Representative immunofluorescence images of D5 VO cryosections showing MECOM (red) and CD31 (white). Nuclei are counterstained with DAPI (blue). Scale bar=50um. (H) Quantification of number of MECOM+ cells (mean ± SD, n= 3 biological replicates, *p<0.05 by paired student’s t-test). (I) mRNA expression of MECOM in D5 ACD3 and iACD3-VOs (mean ± SD, n= 3 biological replicates, **p<0.01 by paired student’s t-test). (J) Schematic diagram illustrating the isolation CD31+ EPCs from D5 VOs using magnetic sorting. (K) mRNA expression of *PECAM1*, *MECOM, TLL1, and TFPI* in CD31+ ACD3 and iACD3-EPCs (mean ± SD, n= 3 biological replicates, **p<0.01, ****p<0.0001 by paired student’s t-test). (L) Representative brightfield images of endothelial tubes. Scale bar=400um. (M) Quantification of tube formation parameters (mean ± SD, n= 3 biological replicates, **p<0.01, ***p<0.001, ****p<0.0001 by one-way ANOVA followed by Tukey’s test).

Given the known role of MECOM in EC development, we reasoned that it might be a direct FOXF1 target that could account for the ACD3 phenotypes observed at day 5. A closer examination of the ATAC-seq data in EPCs at the *MECOM* locus revealed four chromatin accessible peaks with FOXF1 binding motifs that exhibited dramatically reduced accessibility in ACD3-EPCs (**Fig. 5E** and **F**). Upon re-examination of published Foxf1-ChIP-seq dataset from murine lung ECs^34^, we observed a similar association of Foxf1 and *Mecom* (**Fig. S5C**). Immuno-staining and bulk RT-qPCR of day 5 ACD3-VOs confirmed the severe loss of MECOM expression in ACD3-EPCs (**Fig. 5G**, **5H, 5I,** and **S5D**). Expression of *MECOM* was unaffected in ACD1 or ACD2-EPCs, suggesting that the pathology is unique to ACD3 (**Fig. S5E**). Interestingly, day 5 FOXF1 LOF-VOs also showed loss of MECOM when compared to controls, again suggesting that ACD3 pathology resembles FOXF1-LOF (**Fig. S5F** and **S5G**).

To assess the effects of MECOM loss on ACD3-EPC function, CD31+ EPCs were magnetically isolated from day 5 ACD3 and iACD3-VOs for endothelial functional tests (**Fig. 5J** and **S5H**). First, downregulation of *MECOM* expression was confirmed in the isolated ACD3-EPCs and not ACD1 or ACD2-EPCs (**Fig. 5K** and **S5I**). Then, we observed that ACD3-EPCs demonstrated a marked reduction in migration (**Fig. S5J** and **S5K**) and exhibited reduced tube formation capacity relative to iACD3-EPCs (**Fig. 5L** and **5M**). Overexpression of *MECOM* via lipid nanoparticle-mediated delivery of *MECOM* mRNA (LNP-*MECOM*) (**Fig. S5L** and **S5M**) significantly restored endothelial tube formation capacity in ACD3-EPCs compared to controls (ACD3-EPCs treated with LNP-*eGFP*), though still not to the extent observed in iACD3-EPCs. (**Fig. 5L** and **M**). Thus, ACD3-EPC function was markedly compromised, partially due to the loss of MECOM.

To determine the transcriptional profile of these dysfunctional ACD3-EPCs, GO analysis of upregulated DEGs (BP) in ACD3-EPCs was performed. Enrichment of several neural-related terms was observed (**Fig. S5N**), suggesting that ACD3-EPCs may have acquired aberrant neuronal-like transcriptomic features, potentially due to MECOM deficiency-mediated loss of endothelial identity.

### Diminished mural cell identity and collagen signaling network in ACD3-MPCs

During vascular development, EPCs interact closely with MPCs to form, stabilize, and remodel vascular networks^35^. MPCs, which are precursors of vascular mural cells such as pericytes and smooth muscle cells, are necessary for proper vasculogenesis, angiogenesis, vascular maturation, and function. Thus, the loss of capillary development in ACD3-VOs could also be, in part, attributed to the loss of MPC population, which was reduced by 50% (**Fig. S6A**). Notably, expression of collagen genes *COL3A1*, *COL6A3*, *COL6A2*, and *COL5A2* were also reduced in the ACD3-MPCs (**Fig. S6B,** and **S6C**). GO analysis of downregulated transcripts in ACD3-MPCs showed BP terms associated with extracellular matrix (ECM) organization, while the upregulated transcripts in ACD3-MPCs were enriched for GO terms associated with neuronal identity (**Fig. S6D**), similar to the transcriptional profile of ACD3-EPCs. Collectively, this suggest that ACD3-MPCs lacked crucial ECM-related transcriptome and instead possess unusual ‘neuronal-like’ features, potentially negatively impacting regular MPC function. CellChat^36^ analysis was subsequently performed to infer cell-cell interactions from the transcriptome of day 5 VOs (**Fig. S6E**). In control VOs, MPCs serve as the main senders of the collagen signals such as COL4A5, COL4A1, and COL4A2, while EPCs primarily serve as signal receivers (**Fig. S6F**). However, when compared to iACD3, day 5 ACD3-VOs showed a dramatic reduction in the numbers and strength of MPC-EPC interactions, specifically with MPCs as senders and EPCs as receivers (**Fig. S6G**). Instead, interactions between MPC and NEP were increased (**Fig. S6G**). In day 5 ACD3-VOs, NEP and EPCs, instead of MPCs, became the main senders of the collagen signaling components (**Fig. S6H**). This shift was likely due to reduced MPC identity, coupled with the abnormal expansion of the NEP population in day 5 ACD3-VOs, which became active signaling hubs (**Fig. 4C, 4D,** and **S6I**). Additionally, there were significantly fewer total ligand-receptor (L-R) pairs in the collagen signaling pathway network across all cells in day 5 ACD3-VOs (n=16 in ACD3-VOs v.s. n=27 in iACD3-VOs) (**Fig. S6J**), suggesting that in addition to the significant reduction of MPC population in ACD3-VOs (**Fig. 4C** and **4D**), ACD3-MPCs collagen signaling network was markedly diminished in ACD3-MPCs.

### ‘Moderate’ FOXF1 variants differentially rewired EPC and MPC states and functions

The overall proportions of EPCs and MPCs in day 5 ACD1 and ACD2-VOs were relatively unchanged, unlike day 5 ACD3-VOs (**Fig. 2K, 2L, 4C, 4D**, and **S7A**). This is consistent with a more ‘moderate’ capillary phenotype observed in day 15 ACD1 and ACD2-VOs compared to ACD3-VOs (**Fig. 1B-1F, 2K, 2L**, and **S7A**). To address why EPCs in ACD1 and ACD2-VOs, although abundant, exhibit reduced capacity to reconstitute capillary networks, we assessed EPC transcriptional states, functional properties, and EPC–MPC interactions in more detail. Firstly, to better define cell states, we performed unintegrated, paired comparisons between day 5 Healthy donor and ACD1-VOs, as well as iACD2 and ACD2-VOs. Three main EPC sub-clusters, c1, c2, and c3, were observed in day 5 Healthy donor and ACD1-EPCs (**Fig. 6A**). Strikingly, EPC c1 was absent in ACD1-EPCs, while EPC c2 was absent in Healthy donor-EPCs (**Fig. 6A** and **6B**). GO term (BP) analysis of DEGs in EPC c1 revealed terms like ‘endothelial cell development’ which reflect a typical EPC signature, while c2 showed terms associated with cartilage development, suggesting that c2 could represent an aberrant cell state (**Fig. 6C**). This abnormal shift in EPC transcriptional state in ACD1-EPCs was found to be associated with EPC dysfunction as illustrated by reduced tube formation (**Fig. 6D** and **6E**).

**Figure 6.**
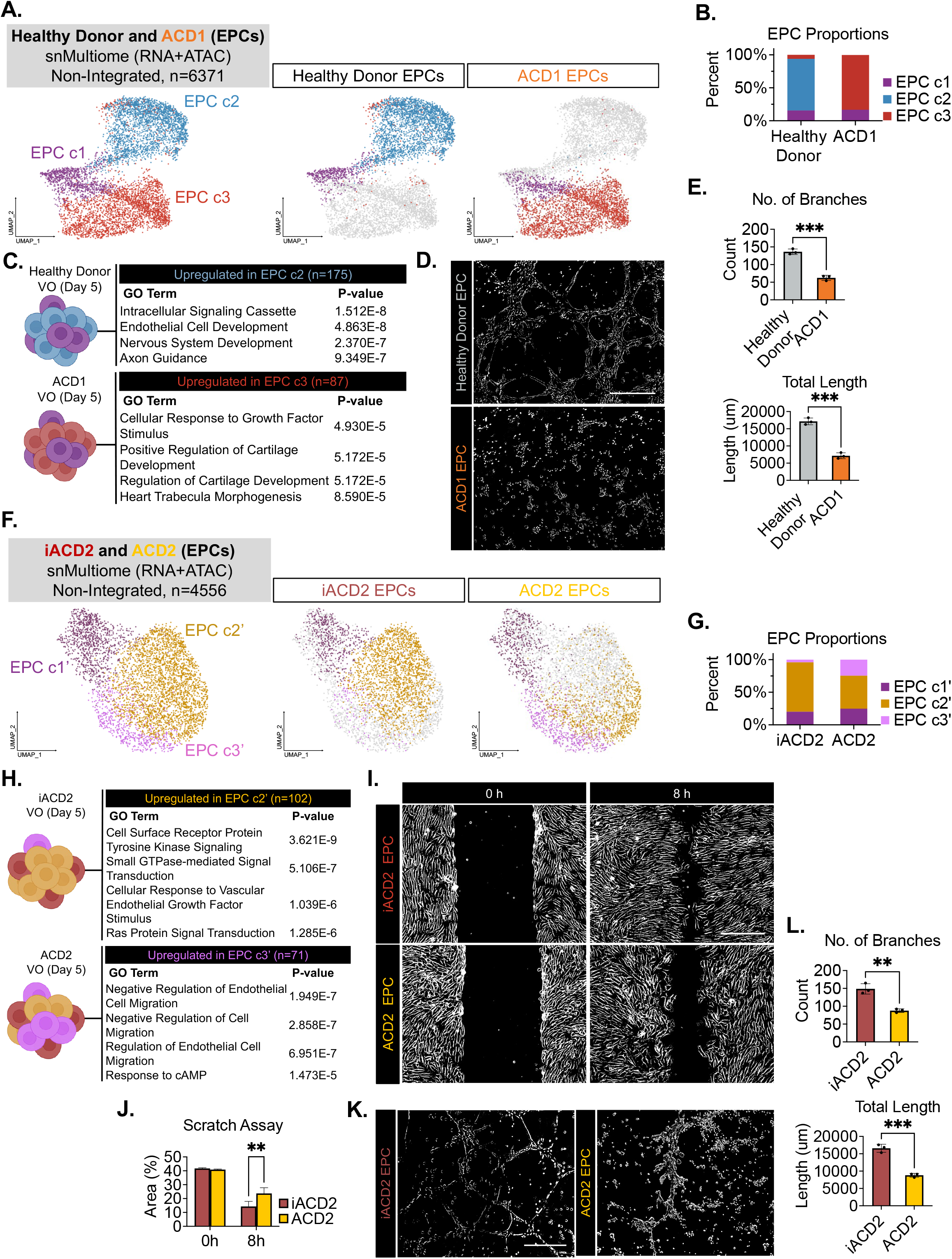
Differential rewiring of EPC transcriptional states and function in ACD1 and ACD2-EPCs. (A) UMAP visualization of EPCs of D5 Healthy Donor and ACD1-VOs (non-integrated, n=6371). Clustering was performed using the Louvain algorithm at a resolution of 0.4. (B) Cell proportion plots of EPC subclusters in Healthy Donor and ACD1-EPCs. (C) GO terms (BP) for DEGs associated with EPC c1 and EPC c2. (D) Representative brightfield images of endothelial tubes. Scale bar=400um. (E) Quantification of tube formation parameters (mean ± SD, n= 3 biological replicates, ***p<0.001 by student’s t-test). (F) UMAP visualization of EPCs of D5 iACD2 and ACD2-VOs (non-integrated, n=4556). Clustering was performed using the Louvain algorithm at a resolution of 0.4. (G) Cell proportion plots of EPC subclusters in iACD2 and ACD2-EPCs. (H) GO terms (BP) for DEGs associated with EPC c2’ and EPC c3’. (I) Representative brightfield images illustrating migration of EPCs at 0 and 8h. Scale bar=400um. (J) Quantification of area of scratch at 0 and 8h (mean ± SD, n= 3 biological replicates, ***p<0.001 by student’s t-test). (K) Representative brightfield images of endothelial tubes. Scale bar=400um. (L) Quantification of tube formation parameters (mean ± SD, n= 3 biological replicates, **p<0.01, ***p<0.001 by student’s t-test).

In day 5 iACD2 and ACD2-EPCs, three EPC sub-clusters were identified – c1’, c2’, and c3’ (**Fig. 6F**). Gene markers that defined EPC c1’ were expressed in EPC c1 identified in Healthy Donor and ACD1-EPCs (**Fig. S7B** and **S7C**), suggesting the presence of a common EPC sub-population. ACD2-EPCs showed altered proportions of EPC sub-clusters - the proportion of EPC c2’ was reduced in ACD2, while the proportion of EPC c3’ was increased in ACD2 (**Fig. 6F** and **6G**). GO terms identified in c3’ include ‘negative regulation of EC migration’ (**Fig. 6H**). To validate this, a scratch assay was performed, revealing that ACD2-EPCs showed reduced migration capacity (**Fig. 6I** and **6J**), along with significant loss of endothelial tube formation capacity (**Fig. 6K** and **6L**).

Transcriptomic changes were also observed in ACD1-MPCs, where genes such as *BST2, FTL, HPSE2, NTRK2*, and *VEGFC* were significantly upregulated while genes such as *UTY, NLGN4Y, KHDRBS2, NRG3,* and *LRP1B* were significantly downregulated (**Fig. S7D**). A dramatic loss of MPC c3, defined by GO terms associated with cell cycle regulation, coupled with the expansion of MPC c4, defined by GO terms associated with ECM remodeling, was observed in ACD1-MPCs (**Fig. S7F** and **S7G**), possibly hinting at a loss of a proliferative progenitor pool and an enrichment of a possible pro-fibrotic MPC population. Transcriptomic changes were observed in ACD2-MPCs when compared to controls (**Fig. S7H**), specifically genes including *PDE5A, MAGI2, AC007091.1, EPHA7*, and *LUM* were significantly downregulated in ACD2-MPCs, while *LINC02511, FTL, C5orf17, PDE3A*, and *SLC8A1* were significantly upregulated (**Fig. S7H**). ACD2 MPCs showed reduced proportions of MPC clusters c1 and c4, which showed GO terms associated with ECM organization and cell cycle regulation, respectively (**Fig. S7I**), suggesting impaired MPC function. Additionally, CellChat analysis revealed changes in EPC-MPC interactions and signaling networks in ACD1 and ACD2-VOs (**Fig. S8A-S8D**). These data suggest that different *FOXF1* allelic variants in ACD1 and ACD2 altered EPC and MPC states through variant-specific mechanisms, resulting in EPC dysfunction, potential impairment of MPC function, and altered EPC-MPC interactions that may collectively contribute to capillary maldevelopment seen in ACD1 and ACD2-VOs.

### Variant- and developmental-stage-specific delivery of LNP-FOXF1 rescue capillary loss

The possibility of restoring wild-type FOXF1 levels via nanoparticle delivery of *FOXF1* mRNA (LNP-*FOXF1*) in a patient-specific manner was assessed (**Fig. 7A**). So far, our results revealed that ACD3 presents early developmental abnormalities at day 3, while vascular anomalies in ACD1 and ACD2 do not emerge until day 5. Therefore, we attempted to restore wild-type FOXF1 levels at different time points. LNP-*FOXF1* was delivered to ACD3-VOs (48h treatment) at different differentiation stages – days 0, 3, and 5 (**Fig. S9A**). mRNA expression of vascular markers *PECAM1* and *PDGFRβ* in D6 VOs was observed to be significantly increased upon LNP-*FOXF1* delivery at days 0 and 3, but not at day 5, when compared to controls (ACD3-VOs treated with *LNP-eGFP*) (**Fig. 7B**). More strikingly, capillary formation was significantly increased in day 0 and day 3 treatment groups, with day 0 treatment showing the most pronounced increase (**Fig. 7C** and **7D**). However, day 5 treatment failed to rescue capillary formation (**Fig. 7C** and **7D**).

**Figure 7.**
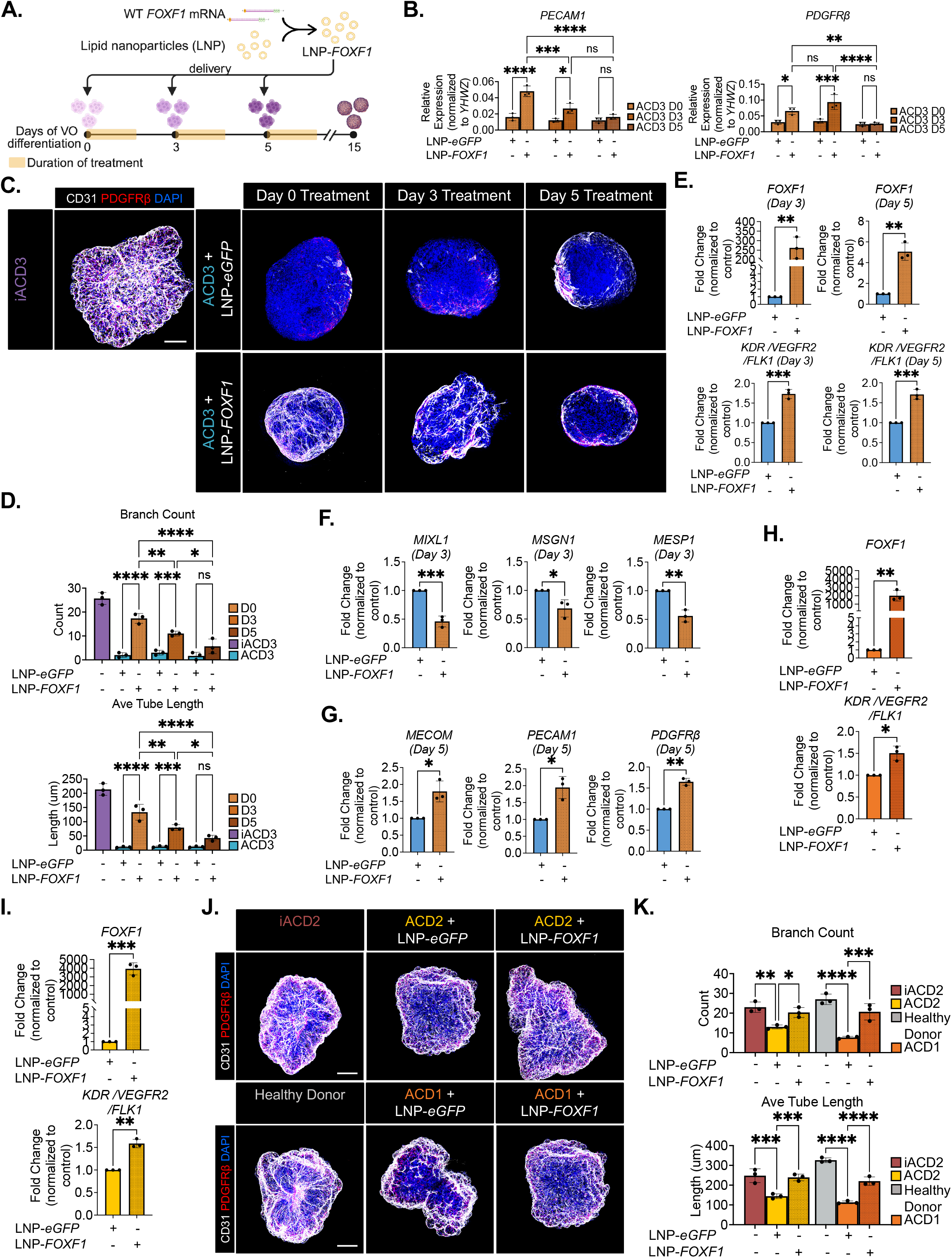
Variant- and developmental-stage-specific LNP-*FOXF1* delivery to ACD-VOs promoted capillary formation. (A) Schematic diagram illustrating the delivery of LNP-*FOXF1* into differentiating ACD3-VOs. (B) mRNA expression of *PECAM1* and *PDGFRβ* in treated (LNP-*FOXF1*) and control (LNP-*eGFP*) ACD3-VOs at D6 across different treatment conditions. (C) Representative whole-organoid immunofluorescence images showing CD31 (white), PDGFRβ (red). Nuclei are counterstained with DAPI (blue). Scale bar represents 100um. (D) Quantification of vessel parameters (mean ± SD, n=3 organoids from 3 biological replicates, *p<0.05, **p<0.01, ***p<0.001 by two-way ANOVA followed by Bonferroni’s test) (E) mRNA expression of *FOXF1* and *FLK1* in treated (LNP-*FOXF1*) and control (LNP-*eGFP*) ACD3-VOs at D3 and D5 (mean ± SD, n= 3 biological replicates, **p<0.01, ***p<0.001 by one-way ANOVA followed by Tukey’s test). (F) mRNA expression of NasM markers in treated (LNP-*FOXF1*) and control (LNP-*eGFP*) ACD3-VOs at D3 (mean ± SD, n= 3 biological replicates, *p<0.05, **p<0.01, ***p<0.001 by one-way ANOVA followed by Tukey’s test). (G) mRNA expression of *MECOM, PECAM1,* and *PDGFRβ* in treated (LNP-*FOXF1*) and control (LNP-*eGFP*) ACD3-VOs at D5 (mean ± SD, n= 3 biological replicates, *p<0.05, **p<0.01, ***p<0.001 by one-way ANOVA followed by Tukey’s test). (H) mRNA expression of *FOXF1* and *FLK1* in treated (LNP-*FOXF1*) and control (LNP-*eGFP*) ACD1 and ACD2-VOs at D6 (mean ± SD, n= 3 biological replicates, **p<0.01, ***p<0.001 by one-way ANOVA followed by Tukey’s test). (I) Representative whole-organoid immunofluorescence images showing CD31 (white), PDGFRβ (red). Nuclei are counterstained with DAPI (blue). Scale bar represents 100um. (J) Quantification of vessel parameters (mean ± SD, n=3 organoids from 3 biological replicates, *p<0.05, **p<0.01, ***p<0.001, ****p<0.0001 by two-way ANOVA followed by Bonferroni’s test).

The effects of D0 LNP-*FOXF1* treatment were scrutinized further. At day 3, there was a 250-fold increase in F*OXF1* mRNA expression, and at day 5, *FOXF1* mRNA expression remains significantly upregulated at around 5-fold (**Fig. 7E**), illustrating successful uptake of LNP-*FOXF1* and persistence of *FOXF1* mRNA for up to 5 days. Consequently, FLK1 mRNA levels were increased by approximately 1.7-fold at both days 3 and 5, indicating that a functionally sufficient level of FOXF1 protein was produced (**Fig. 7E**). Day 0 LNP-*FOXF1* treatment was also able to result in a significant downregulation of NasM, NE, and Epi markers in day 3 ACD3-VOs (**Fig. 7F** and **S9B**), illustrating successful reversal of developmental abnormalities. Moreover, day 0 LNP-*FOXF1* treatment was further shown to drastically reduce NasM and NE markers in day 3 FOXF1-LOF VOs (**Fig. S9C**). At day 5, day 0 LNP-*FOXF1* treatment was also able to promote a significant increase in *MECOM, PECAM1,* and *PDGFRβ* expression (**Fig. 7G**).

As ACD1 and ACD2 exhibited abnormalities at a later time point, LNP-*FOXF1* was administered to differentiating ACD1 and ACD2 VOs on day 3 (48-hour treatment) (**Fig. S9D**). LNP-*FOXF1* treatment significantly induced FOXF1 mRNA expression - ∼2000-fold increase in ACD1 and ∼4000-fold in ACD2 compared to *LNP-eGFP* treated VOs (**Fig. 7H** and **7I**). Correspondingly, *FLK1* mRNA expression was increased by around 1.5-fold in both ACD1 and ACD2 VOs (**Fig. 7H** and **7I**). Of note, mRNA expression of vascular markers *PECAM1* and *PDGFRβ* remained relatively unchanged in ACD1 and ACD2-VOs upon LNP-*FOXF1,* suggesting that the overall number of vascular progenitors was not altered (**Fig. S9E**). Nonetheless, day 3 LNP-*FOXF1* treatment was observed to significantly increase capillary formation in both ACD1 and ACD2 (**Fig. 7J** and **7K**), thereby suggesting that elevating wild-type FOXF1 levels can act to promote proper vascular specification and, consequently, capillary development in ACD1 and ACD2-VOs, potentially by exerting effects on vascular progenitor states and/or function rather than overall progenitor population size.

## DISCUSSION

Our study employs patient-specific organoid models to investigate a fundamental question in mesoderm and vascular development: how allelic variants of a TF give rise to divergent developmental outcomes in humans. Here, we show that distinct FOXF1 allelic variants produce divergent developmental trajectories during mesoderm and vascular development, resulting in capillary maldevelopment through variant-specific molecular mechanisms. This reflects known phenotypic variability present in ACDMPV patients^10–13^. More specifically, we identified a ‘severe’ c.253T>A *FOXF1* variant (p.F85I) found in ACD3, which resulted in the most severe extent of vascular maldevelopment. The c.166C>G (p.L56V) and 16q24.1del *FOXF1* variants found in ACD2 and ACD1, respectively, however, exhibited a comparatively ‘moderate’ phenotype. Interestingly, although the ACD2 and ACD3 variants are both missense mutations located within the FOXF1 DNA-binding domain, they lead to strikingly different degrees of capillary maldevelopment. Time-course single-cell transcriptomics revealed variant-specific disease onset, with abnormalities emerging in ACD3 as early as the mesoderm differentiation stage, whereas vascular defects in ACD1 and ACD2 appeared after the vascular specification stage.

Previous studies in Foxf1^−/+^ mice demonstrated a dose-dependent requirement for Foxf1 during pulmonary vascular development^4,5,37^, with lower FOXF1 levels associated with more severe vascular and lung developmental defects. Consistent with these findings, FOXF1-LOF VOs showed a correlation between FOXF1 expression and vascular formation. FOXF1 and FLK1 expression were reduced in both ACD1 and ACD2 at days 3 and 5, with higher levels in ACD2 than in ACD1, corresponding to the milder capillary phenotype observed in ACD2-VOs. In contrast, ACD3 exhibited the most severe capillary defects despite expressing higher FOXF1 protein levels than ACD1 and ACD2. Transcriptomic analysis revealed delayed *FOXF1* expression in ACD3 at day 3, suggesting that disruption of the temporal window of FOXF1 activity impaired activation of FLK1 and downstream programs required for NasM-to-LPM differentiation. Consistent with this model, FLK1 expression was nearly absent in ACD3. Alternatively, the c.253T>A *FOXF1* variant in ACD3 may produce a functionally impaired FOXF1 protein, resulting in loss of downstream target activation despite preserved protein expression. Together, these findings suggest that both the activity and timing of FOXF1 are critical for proper pulmonary vascular development.

Structural modeling and molecular dynamics simulations suggest that the L56V and F85I amino acid substitutions may differentially alter FOXF1 DNA-binding behavior and conformational dynamics. Notably, the increased solvent-accessible surface area predicted for L56V FOXF1 could modify interactions with transcriptional co-regulators while preserving partial FOXF1 activity, providing a potential explanation for its comparatively milder phenotype. Others have found that FOXF1 cooperates with ETS transcription factors in pulmonary ECs^38,39^. FOXF1 have also been shown to bind to co-activators such as serum response factor and myocardin during smooth muscle development^40^. Therefore, it is possible that the altered flexibility of L56V FOXF1 compromises interactions with other TFs or co-activators that are necessary for proper transcriptional regulation. Unlike L56V FOXF1, a significant reduction in DNA-binding interactions was predicted in F85I FOXF1, which could account for the impaired FOXF1 function in ACD3. Indeed, the ability of F85I FOXF1 to bind to DNA has been shown by others to be severely attenuated^16^. Regardless, our findings illustrate the diverse effects of allelic variants on the molecular and biophysical attributes of TFs such as FOXF1.

At present, FOXF1 HI is widely accepted to be the main cause of ACDMPV^7,41^. Therefore, we compared the *FOXF1* variants with FOXF1-LOF models. We found that the molecular and phenotypic outcomes of the *FOXF1* variants differ substantially from those observed in FOXF1 haploinsufficiency models, suggesting that ACDMPV cannot be universally explained by a simple model of FOXF1 haploinsufficiency. Unlike ACD1 and ACD2, FOXF1-LOF and ACD3-VOs exhibited impaired NasM-to-LPM differentiation at day 3. While NasM was able to form in FOXF1-LOF and ACD3 at day 3, these precursors appeared unable to further differentiate into LPM, illustrating that NasM differentiation is likely to be independent of FOXF1^42^ and that FOXF1 is necessary for NasM-to-LPM differentiation. While our study identified a novel role of FOXF1 in regulating NasM-to-LPM differentiation, further studies are needed to define the FOXF1-mediated gene regulatory network (GRN) underlying this process in greater detail.

The reduced LPM population in ACD3 likely resulted in the reduced proportion of EPCs and MPCs in day 5 ACD3-VOs, which are derivatives of LPM. To determine why the remaining EPCs and MPCs in day 5 ACD3-VOs failed to form well-developed capillary networks, we examined regulators of vascular progenitor specification and function. We found that critical vascular lineage regulator *MECOM*, which is a direct FOXF1 target, was significantly downregulated in ACD3-EPCs and that the function of ACD3-EPCs was severely compromised. Others have shown that knockout of *MECOM* not only resulted in loss of both ECs and mural cells in mature VOs, but also dramatic changes in EPC and MPC states^32^. Additionally, MECOM is necessary for maintaining endothelial identity and function^33^. Therefore, the loss of MECOM in ACD3-EPCs likely impaired cellular function and further development into capillary networks. Overexpressing *MECOM* in ACD3-EPCs, although significantly improved endothelial function, offers only a partial rescue, suggesting that additional FOXF1 target genes that were downregulated in ACD3-EPCs, such as *TLL1, TFPI, FLT1,* and *CALCRL* could also be contributing to reduced EPC function. Further studies could be done to determine the roles of TLL1, TFPI, and CALCRL in EPC function and development.

In tandem, the MPC population was also found to be diminished on day 5 ACD3-VOs, and ACD3-MPCs were found to exhibit loss of mural identity and collagen signaling in ACD3-MPCs. Mural cells typically produce collagen, an essential component of vascular ECM that supports vasculogenesis and angiogenesis during vascular development^43,44^. Therefore, the loss of mural cell fate identity and collagen production in ACD3-MPCs, collectively, could impair proper vessel formation, therefore contributing to capillary maldevelopment in ACD3-VOs. Vascular development also relies on close interactions between EPCs and MPCs^45^; thus, the combined loss of EPC and MPC populations and expansion of NEPs, which significantly altered EPC-MPC crosstalk, coupled with impaired EPC and MPC functions, likely drive the severe capillary maldevelopment phenotype. Overall, we propose a ‘two-hit’ model for ACD3 - First, delayed FOXF1 expression and/or impaired FOXF1 activity disrupts NasM-to-LPM differentiation at day 3. Second, loss of FOXF1-mediated MECOM expression compromises vascular progenitor specification and function at day 5, ultimately impairing capillary formation.

Although LPM differentiation was unaffected in ACD1 and ACD2-VOs, and overall EPC and MPC population sizes were unchanged, we uncovered pathogenic alterations to EPC transcriptional states that are associated with reduced EPC function. *DOCK4* is one of the top downregulated DEGs in ACD1-EPCs. Interestingly, DOCK4 was found to be important in endothelial barrier function^46^ and blood vessel lumen morphogenesis during angiogenesis. Future studies should determine if FOXF1 acts to regulate DOCK4 in EPCs, and if restoring DOCK4 levels could restore transcriptional and functional programs in ACD1-EPCs and promote proper capillary development in ACD1-VOs. Additionally, we also uncovered an increase in anti-migratory EPCs in day 5 ACD2-VOs. Endothelial migration is a fundamental process during vasculogenesis^48^, and Foxf1 was previously found to regulate cell migration in mouse fetal lung ECs^49^. Hence, the anti-migratory EPCs in ACD2-VOs could possibly have stifled proper capillary formation and maturation. It is plausible that reduced migration capacity, coupled with altered MPC states and EPC-MPC interactions, collectively contributed to capillary maldevelopment in ACD2-VOs.

The ACD hiPSC lines used in this study were derived from three distinct ACDMPV patients and therefore retain their respective genetic backgrounds, which may contribute to the observed phenotypes. Although isogenic controls (iACD2 and iACD3) were generated to isolate the effects of the corresponding FOXF1 variants, patient-specific genetic modifiers may influence transcriptomic and epigenomic programs and thereby affect capillary development. Consequently, some differences observed among ACD1, ACD2, and ACD3 may reflect interactions between FOXF1 variants and the broader genetic background. Future studies incorporating additional patient-derived and isogenic iPSC models will be important for determining the extent to which genetic background modifies FOXF1-dependent developmental outcomes.

Presently, ACDMPV remains incurable. Patients typically receive cardiorespiratory support before eventually succumbing to respiratory failure^9^. Lung transplantation promotes long-term survival but is only feasible for atypical/ late-presenting cases^50,51^. Therefore, we sought to determine if restoring WT FOXF1 levels at early developmental stages could promote capillary formation. Specifically, we showed that introducing LNP-*FOXF1* at developmental stages that preceded pathogenesis, i.e., before LPM and vascular progenitor stages in ACD3 and ACD1/ACD2, respectively, was successful in rescuing capillary maldevelopment to a certain extent. However, dose and timing of LNP-*FOXF1* should be carefully considered since Foxf1 overexpression also leads to pulmonary hypoplasia and vascular defects^52^. Although the effectiveness of LNP-*FOXF1* in treating ACDMPV has not been assessed pre-clinically, others have recently shown that LNP-*Foxf1* treatment was effective in restoring EC proliferation and function in a rodent model of neonatal sepsis^53^, and safety of LNP-*Foxf1* has been demonstrate in postnatal wildtype mice^54^. Recently, we also showed the use of LNP-mediated mRNA delivery to reverse vascular defects associated with bronchopulmonary dysplasia^55^. Our data suggest that introducing LNP-*FOXF1* may potentially correct early developmental abnormalities caused by the *FOXF1* variants. Although the feasibility of embryonic or fetal LNP-mediated mRNA therapies has not been determined clinically, delivery of gene editing therapies has been done in animal models^56^, and more recently, in human embryos^57^. Additionally, others have shown that LNP-mRNA can be effectively introduced prenatally via in utero and intravenous delivery methods^56,58^. Collectively, these findings suggest that LNP-*FOXF1* warrants further investigation as a potential therapeutic strategy for ACDMPV. However, the ethical implications of therapeutic interventions in developing human embryos warrant careful consideration.

In summary, these findings establish a framework for understanding how allelic variation in FOXF1 gives rise to divergent developmental outcomes and variable disease severity in ACDMPV. More broadly, our study demonstrates how patient-derived organoid models and single-cell multiomic approaches can uncover variant-specific mechanisms that may inform precision therapeutic strategies for developmental disorders.

### Limitations of the study

We acknowledge several limitations of this study. Firstly, one of the main clinical features of ACDMPV is the severe loss of alveolar capillaries, particularly the alveolar capillary endothelial populations (CAP1 and CAP2), which are transcriptionally and functionally distinct from ECs found in other organs^10,59,60^. ECs in the hiPSC-derived VOs do not possess any lung-specific features^32^. Therefore, VO differentiation does not represent alveolar capillary EC development and specification. Currently, there are no established protocols for generating pure populations of alveolar capillary ECs from hiPSCs. Nonetheless, VOs are still suitable models to study the effects of *FOXF1* variants on early developmental events such as LPM differentiation and vascular progenitor specification, which precede the acquisition of tissue-specific identities.

Secondly, the pathogenic effects of *FOXF1* variants cannot be completely decoupled from the genetic background in which they arise. Other genetic modifiers may be present in these patients that could either sensitize an already developmentally compromised state, thus further exacerbating disease phenotypes, or conversely, provide protective effects. CRISPR/Cas9-mediated knock-in of *FOXF1* variants into normal hiPSCs could disentangle variant-intrinsic effects from the potential influence of underlying genetic modifiers.

Third, while our findings revealed important roles of FOXF1 in mesoderm and capillary development, the FOXF1 GRN remains incompletely defined. Further characterization of FOXF1-dependent regulatory programs will help identify the molecular mechanisms governing these developmental transitions and establish how distinct *FOXF1* variants perturb downstream pathways.

Lastly, while we showed that variant- and developmental*-*stage-specific *FOXF1*-LNP treatment was able to promote capillary formation in ACD-VOs. The clinical feasibility of delivering LNP-*FOXF1* to human patients during early developmental stages will require further investigations. Regardless, our study emphasizes the need to consider variant-specific therapeutic strategies. Overall, addressing these limitations will further our endeavor in understanding and treating FOXF1-associated pathologies such as ACDMPV.

## RESOURCE AVAILABILITY

### Lead contact

Requests for further information and resources should be directed to and will be fulfilled by the lead contact, Mingxia Gu.

### Materials availability

The FOXF1-LOF hESC lines are available from Aaron Zorn, and the isogenic controls iACD2 and iACD3 hiPSCs are available from lead contact upon request under a transfer agreement.

### Data and code availability

- Processed and raw 10x single nucleus multiome-seq data is available at GEO: GSE332842 (Token: sniruysotnwzxgp)
- This paper does not report any original code.
- Codes used for data processing and analysis are available at https://github.com/ZornLab/vessel_organoid_analysis and https://doi.org/10.5281/zenodo.20736501
- Original western blot images will be deposited at Mendeley and will be publicly available as of the date of publication. Microscopy data reported in this paper will be shared by the lead contact upon request.

## ACKNOWLEDGMENTS

We thank Dr. Jim Wells, Dr. Aaron Zorn, and Dr. Makiko Iwafuchi, as well as members of their labs, for providing valuable insights, suggestions, and technical aid. Particularly, we thank Dr. Nicole Edwards, Dr. Bailey Weatherbee, Dr. Scott Rankin, Liu Peiyao, and Dr. Chen Jichao for their feedback and suggestions. We also thank Dr. Anne Karina-Perl and Dr. Ya-Wen Chen for their input. We would like to acknowledge Dr. Anmin Wang, Dr. Yifei Miao, Dr. Ziyi Liu, Jessica Babros, Edward Baker, and Owen Focht for their contributions to the study. We thank the following Research Shared Facilities at CCHMC – the Single Cell Genomics Facility, Genomics Sequencing Facility, and the Bio-Imaging and Analysis Facility. We thank the staff of the Transgenic Animal and Genome Editing Facility and Pluripotent Stem Cell Facility at CCHMC, particularly Joseph Shiley, Isheunouya Mazani, and Anushkaa Parwade, for their expertise and assistance in generating the FOXF1 LOF hESC lines. The iACD2 and iACD3 hiPSC lines were generated by the Stem Cell Research Facility at Memorial Sloan Kettering Cancer Center (RRID: SCR_027808). This work was supported by NIH/NHLBI R01HL166283 (M. Gu), CCHMC CURE Award (A.M.Z.), R01HL095993 and N01:75N92025D00035 (D.N.K.), NSF-BSF grant MCB-2136816 (G.S.), and the allocation TG-MCB170020 (G.S.) for supercomputer resources from the ACCESS program, which is supported by National Science Foundation grants #2138259, #2138286, #2138307, #2137603, and #2138296. N.P. received the American Heart Association Pre-doctoral Fellowship (121890), the Albert J. Ryan Fellowship, the University of Cincinnati Graduate Student Government Research Fellowship, and the Genscript Life Science Research Grant. Schematic figures were generated using BioRender.com.

## AUTHOR CONTRIBUTIONS

N.P. and M. Gu conceptualized the project. A.Z. assisted with the conceptualization of the project and the interpretation of the results. K.T. and M. Guo performed bioinformatics analysis. H.D., T.M., and G.S. performed molecular dynamics simulation experiments, and the associated data analysis and interpretation. K.K. and A.Z. generated the FOXF1-LOF hESC lines. R.R. and D.N.K. generated the ACDMPV patient-derived iPSCs. N.P. designed, performed experiments, obtained and interpreted the results. N.P and M. Gu wrote the manuscript. All authors reviewed, provided feedback, and approved the manuscript.

## SUPPLEMENTAL INFORMATION

Document S1 – Figures S1-S9, Tables S1-S5

## KEY RESOURCES TABLE

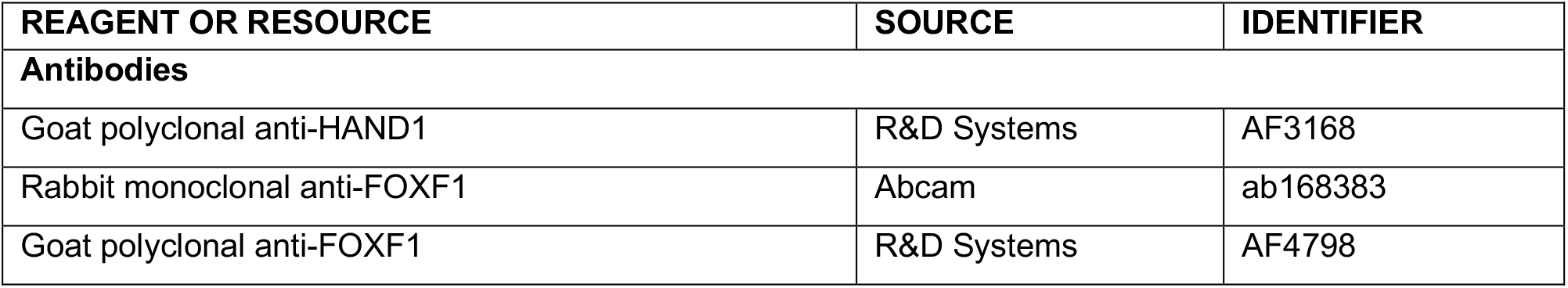
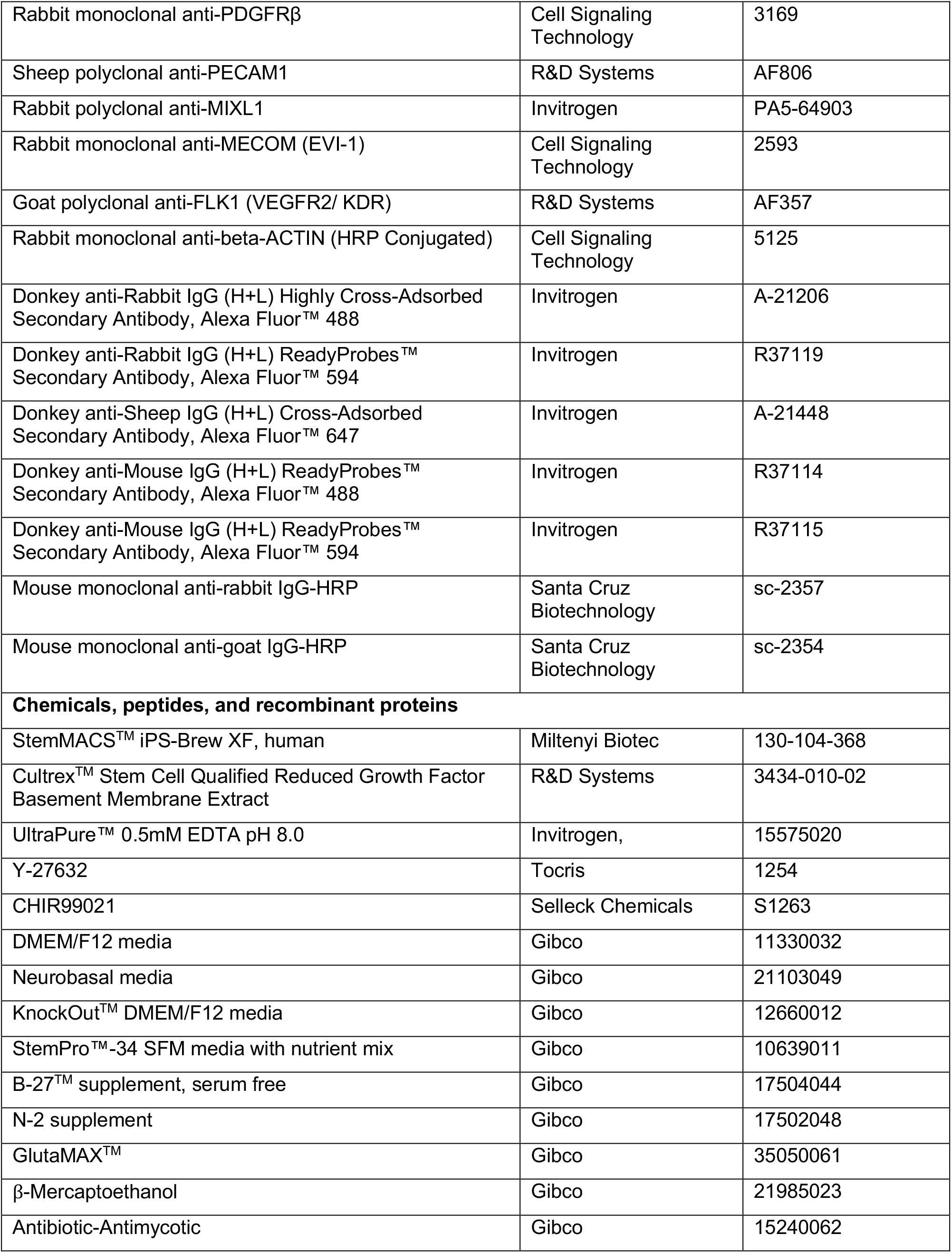
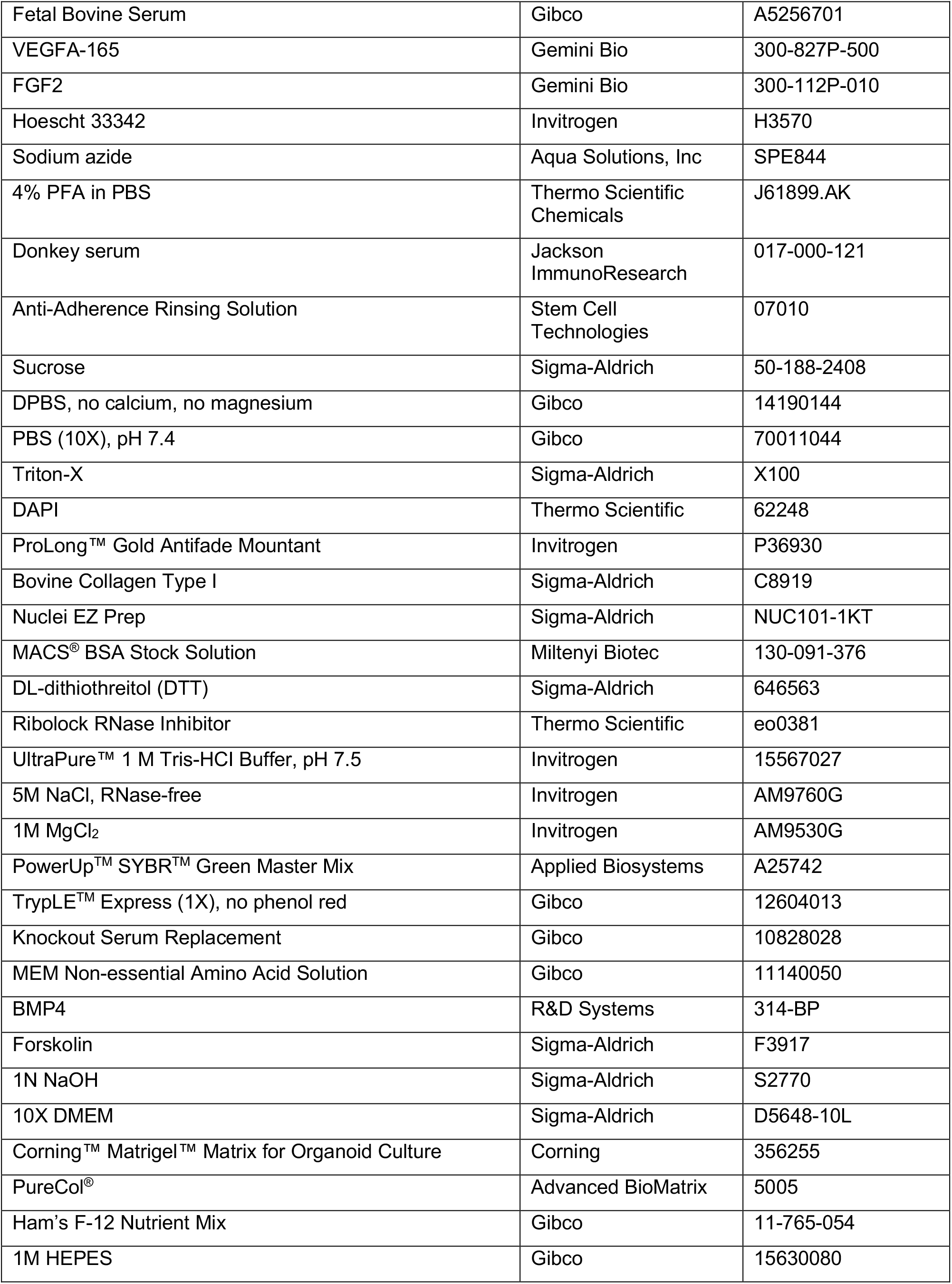
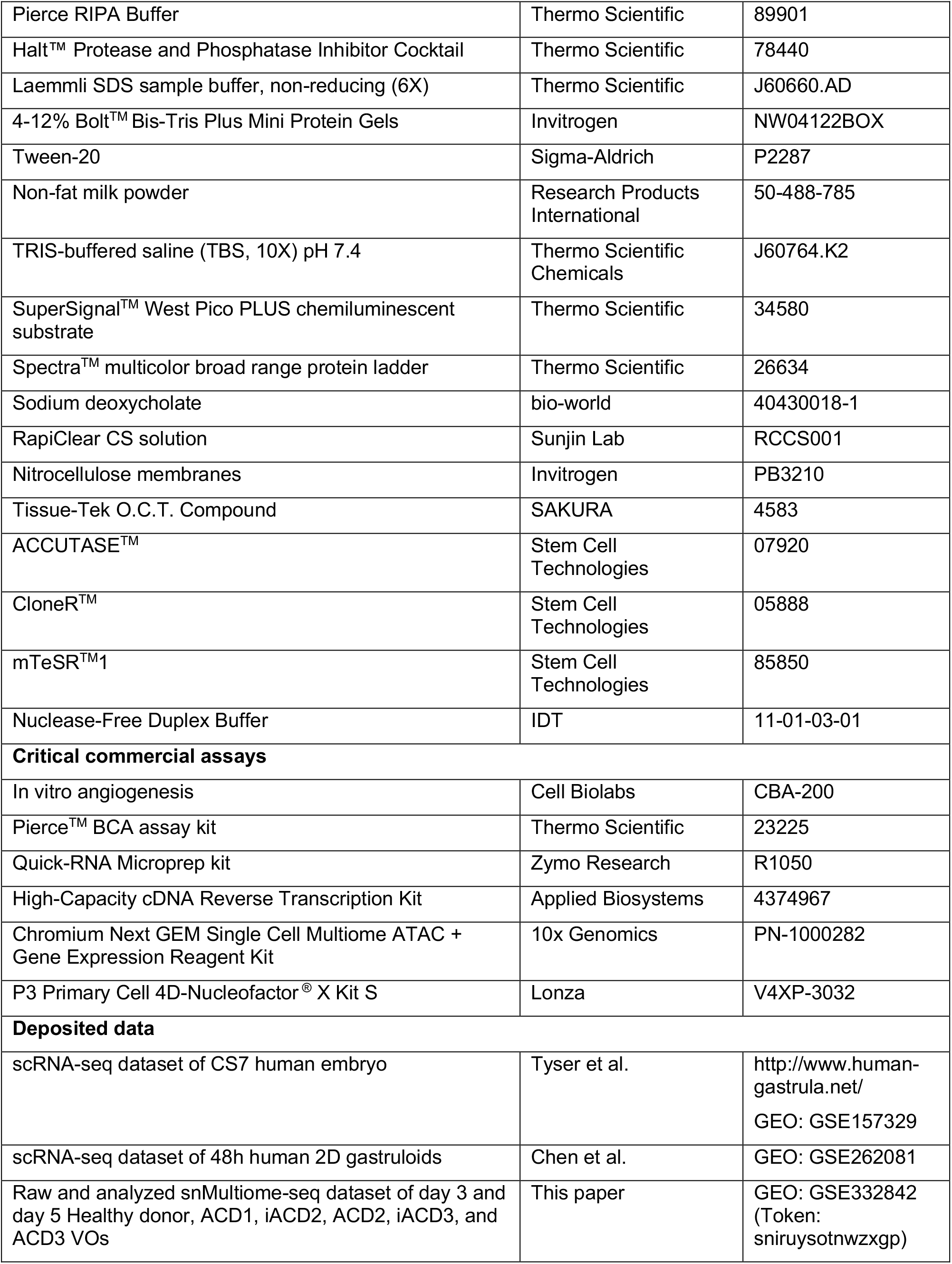
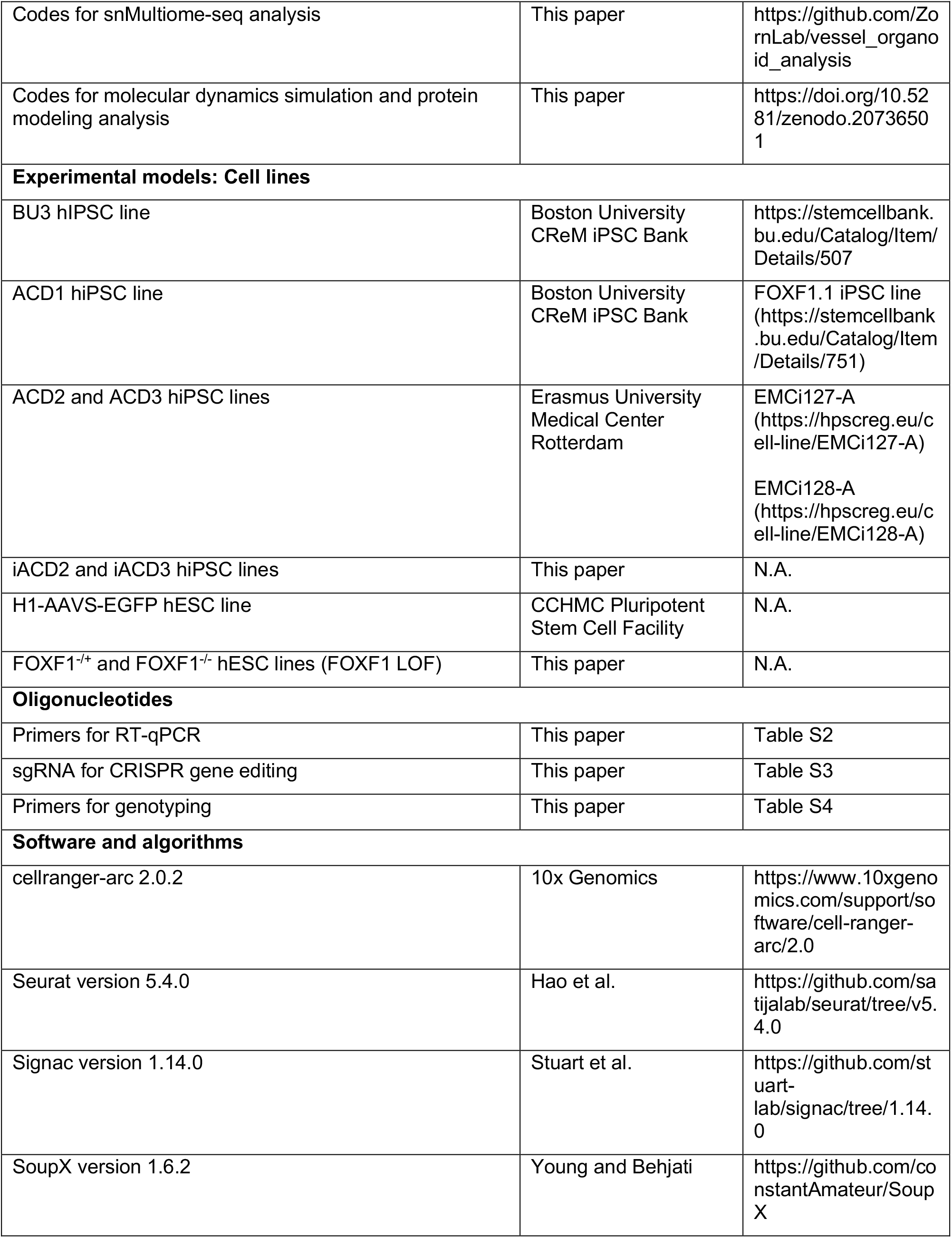
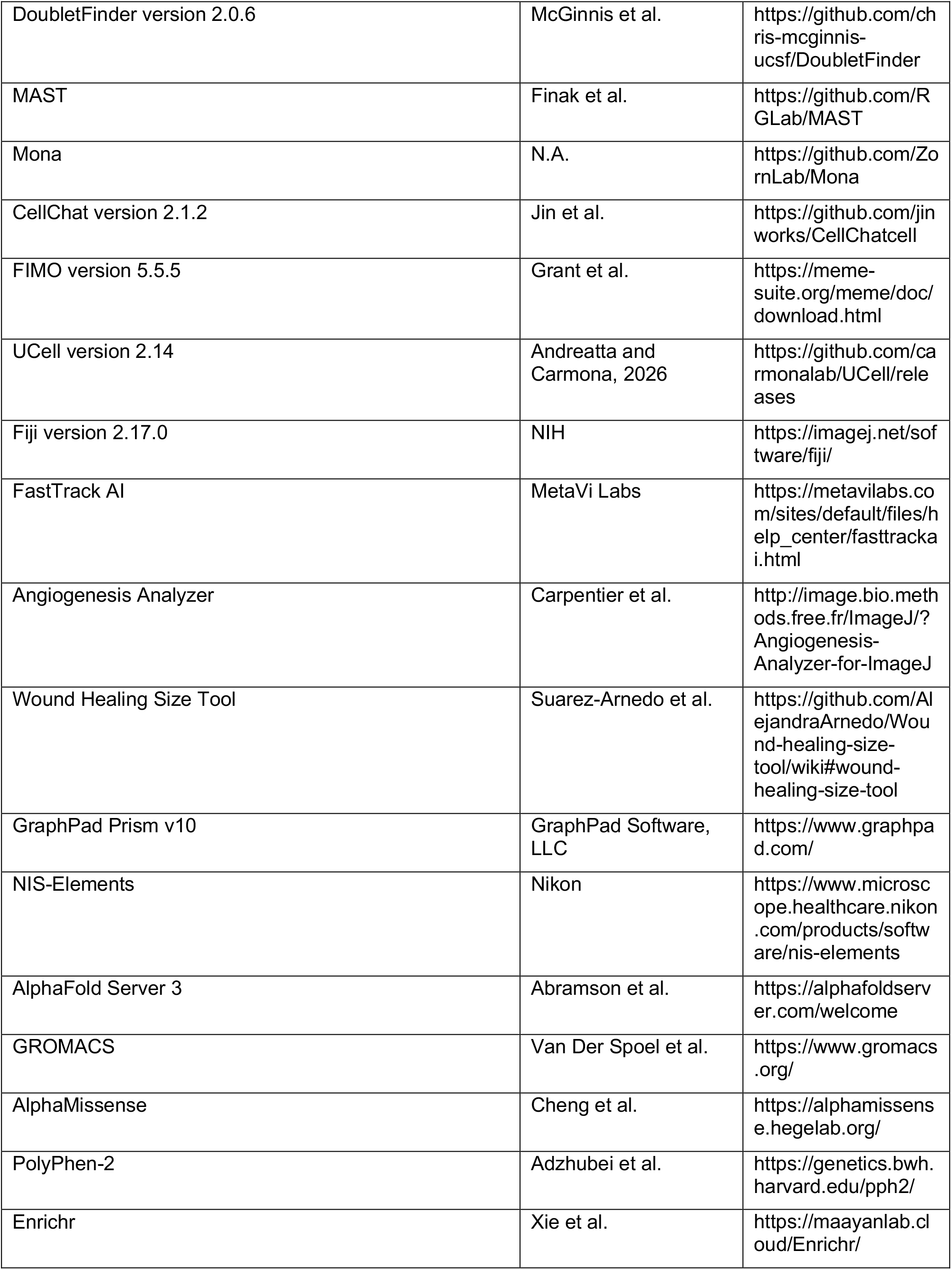
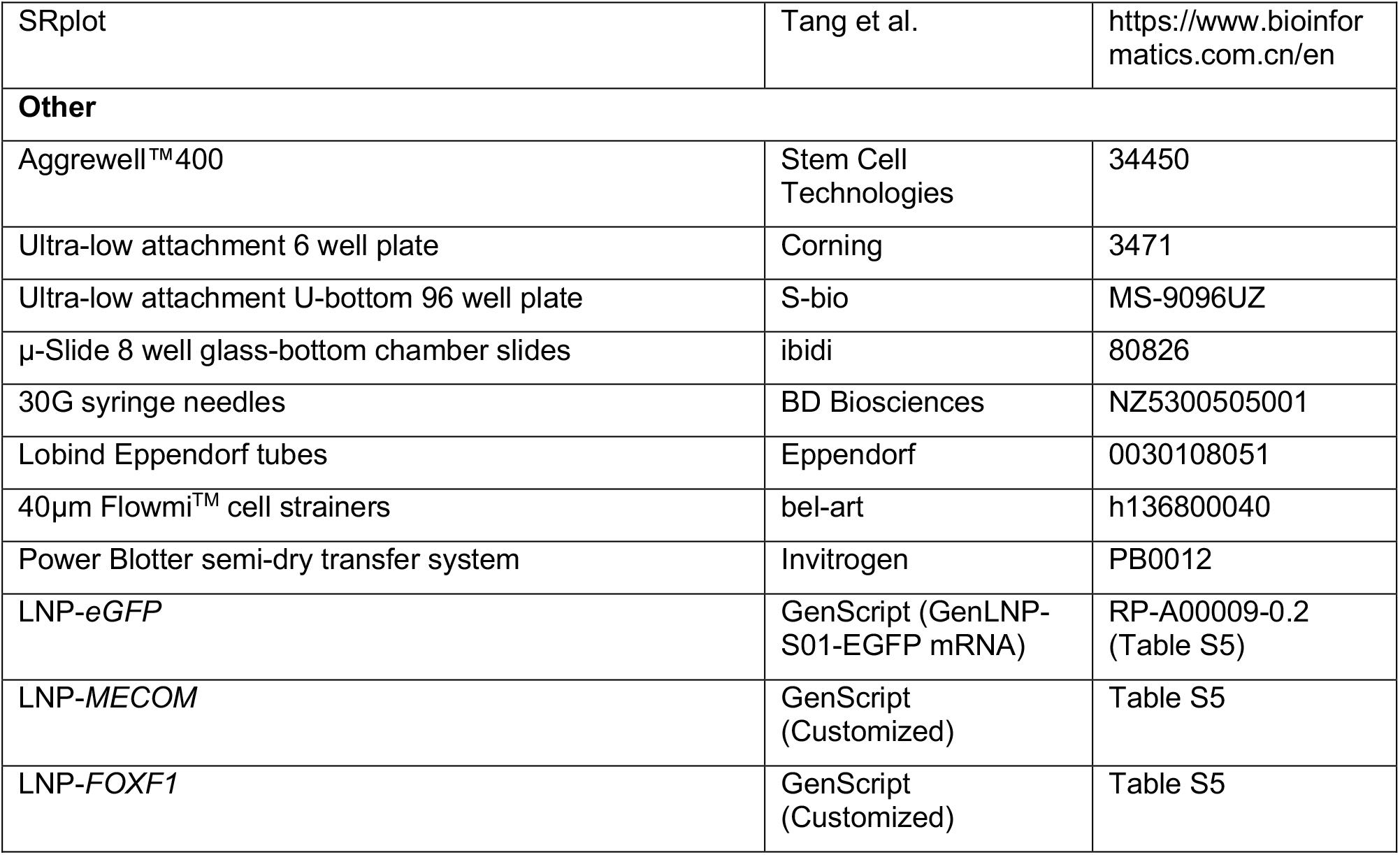

## EXPERIMENTAL MODEL AND STUDY PARTICIPANT DETAILS

### CELL LINES

#### Human Induced Pluripotent Stem Cells (hiPSCs) and Human Embryonic Stem Cells (hESCs)

**Table S1** summarizes ACDMPV patient information. Healthy Donor (BU3, male, 32-year-old) and ACD1 (FOXF1.1) iPSCs were acquired from Boston University^17^. ACD2 (EMC127i-A:ACD871C4) and ACD3 (EMC128i-A:ACD874C9) iPSCs were acquired from Erasmus University Medical Center^19^. iACD2 and iACD3 isogenic controls were generated from ACD2 and ACD3 lines respectively. All human iPSC work was approved by Institutional Review Boards at Cincinnati Children’s Hospital, Boston University, and the Daily Board of the Medical Ethics Committee (METC) Erasmus University Medical Center Rotterdam, The Netherlands. FOXF1-LOF lines – FOXF1^−/+^ and FOXF1^−/−^ were generated using the H1-AAVS-EGFP hESC line^61^.

hiPSCs and hESCs were hPSCs were cultured and maintained based on previously published studies^18,55^. StemMACS™ iPS-Brew XF media (Miltenyi Biotec, 130-127-865) in Cultrex Stem Cell Qualified Reduced Growth Factor Basement Membrane Extract (R&D Systems, 3434-010-02)-coated 6 well plates (Corning) in 37°c under humidified atmosphere of 5% CO_2_. Cells were passaged every 3-5 days using UltraPure™ 0.5mM EDTA pH 8.0 (Invitrogen, 15575020) diluted in DPBS. During passaging, clumps were seeded on Cultrex-coated wells in StemMACS™ iPS-Brew XF supplemented with 10μM of Y-27632 (Tocris, 1254). Fresh media change was performed daily. hiPSC and hESC lines were routinely authenticated for karyotype, pluripotency markers, and mycoplasma contamination.

### METHOD DETAILS

#### Vessel organoid differentiation

Vessel organoids (VOs) were generated based on a protocol described by Wimmer et al.^20,21^ with some slight modifications. hPSCs were first dissociated into single cells and resuspended in Aggregation media consisting of Knockout DMEM/F12 (Gibco, 12660012), 20% Knockout Serum Replacement (Gibco, 10828028), 1% GlutaMAX™ (Gibco, 35050061), 1% MEM Non-essential Amino Acid Solution (Gibco, 11140050), 55µM β-Mercaptoethanol (Gibco, 21985023), and 1% Antibiotic-Antimycotic (Gibco, 15240062) supplemented with 20µM Y-27632 (Tocris, 1254). Filter cells through 40μM Flowmi^TM^ Cell Strainers (Bel-Art, H136800040). 1.2×10^6^ cells were added to each well of the Aggrewell™400 (Stem Cell Technologies, 34450) that was already prepped with Anti-Adherence Rinsing Solution (Stem Cell Technologies, 07010). The Aggrewell™400 consisting of cells was then spun down at 100g for 3 mins to evenly distribute cells into each microwell (1000 cells per microwell). Cells were then left to form embryoid bodies (EBs) overnight at 37°C, 5% CO_2_. The next day, EBs were transferred to N2B27 media supplemented with 12μM CHIR99021 (Selleck Chem, S1263) and 30ng/ml BMP4 (R&D Systems, 314-BP) and cultured in ultra-low attachment 6 well plates on an orbital shaker at 37°C, 5% CO_2_ for three days. Subsequently, the media was changed to N2B27 supplemented with 2μM Forskolin (Sigma-Aldrich, F3917) and 100ng/ml VEGFA-165 (Gemini Bio, 300-827P-500) for two days. Then, on day 5, the aggregates were embedding into a Collagen I-Matrigel sandwich consisting mixture consisting of 11.3% 0.1N NaOH (Sigma-Aldrich, S2770), 4.7% 10x DMEM (Sigma-Aldrich, D5648-10L), 0.9% 1M HEPES (Gibco, 15630080), 0.7% NaHCO_3_ (Gibco, 25080094), 0.5% GlutaMAX (Gibco, 35050061), 6.9% Ham’s F-12 Nutrient Mix (Gibco, 11-765-054), 50% PureCol^®^ (Advanced BioMatrix, 5005), and 25% Organoid Matrigel (Corning, 356255) to induce vessel sprouting. BVI media was added to the collagen-Matrigel sandwiches. Fresh BVI media was changed every 2 days. On day 10, the vessel sprouts were then micro-dissected from the collagen-Matrigel sandwich gels using sterile 30G syringe needles (BD Biosciences, NZ5300505001) and transferred to ultra-low attachment 6 well plates containing fresh BVI media and cultured on an orbital shaker at 37°C for 24h to form VOs. Subsequently, VOs were transferred to ultra-low attachment 96 well plates and supplemented with fresh BVI for the remaining 4 days. BVI media was replaced every 2 days and VOs were harvested at day 15 of differentiation, unless otherwise stated.

#### 3D human gastruloid differentiation

hiPSCs were differentiated to 3D gastruloids as described by Moris et al., 2020^30^ with some slight modifications. 24h before gastruloid formation, 3uM of CHIR99021 (Selleck Chemicals, S1263) was added to hiPSCs (40-60% confluent) to induce primitive streak differentiation. The next day, the CHIR99021-treated hiPSCs were dissociated into single cells and resuspended in N2B27 media containing 3uM CHIR99021 and 5uM Y-27632 (Tocris, 1254). N2B27 media is composed of 50% DMEM/F12 media (Gibco, 11330032), 50% Neurobasal media (Gibco, 21103049), 1% B-27 supplement (Gibco, 17504044), 0.5% N-2 supplement (Gibco, 17502048), 0.5% GlutaMAX^TM^ (Gibco, 35050061), 55µM β-Mercaptoethanol (Gibco, 21985023), and 1% Antibiotic-Antimycotic (Gibco, 15240062). 1.5×10^6^ cells were added to each well of the Aggrewell™400 (Stem Cell Technologies, 34450) that was already prepped with Anti-Adherence Rinsing Solution (Stem Cell Technologies, 07010). The Aggrewell™400 consisting of cells was then spun down at 100g for 3 mins to evenly distribute cells into each microwell (1000 cells per microwell). Cells were left to aggregate overnight at 37°C, 5% CO_2_. Next day, the aggregates were transferred to N2B27 media and cultured in ultra-low attachment 6 well plates on an orbital shaker at 37°C, 5% CO_2_ for three additional days. The N2B27 media was replaced daily.

#### Ac-LDL uptake assay

Vessel organoids were incubated in Blood Vessel Induction (BVI) media containing 5ug/ml of Ac-LDL, Dil complex (Invitrogen, L3484) for 4h in the incubator. BVI media is composed of StemPro™-34 SFM media supplemented with StemPro™-34 nutrient mix (Gibco, 10639011), 1% GlutaMAX™ (Gibco, 35050061), 1% Antibiotic-Antimycotic (Gibco, 15240062), 15% Fetal Bovine Serum (FBS) (Gibco, A5256701), 100ng/ml VEGFA-165 (Gemini Bio, 300-827P-500), and FGF2 (Gemini Bio, 300-112P-010). Then, the nuclei were counterstained with 10ng/ml Hoescht 33342 (Invitrogen, H3570). The organoids were then washed three times with fresh BVI media before being imaged live using the confocal microscope.

#### CRISPR-correction of *FOXF1* missense mutations in ACDMPV hiPSCs

Both the *FOXF1* c.166C>6 and c.253A>T mutations in ACD2 and ACD3 hiPSCs, respectively, were corrected using prime editing. The design and execution of prime editing in iPSCs were performed as previously described^62^. Briefly, pegRNAs containing the respective spacer and 3′ extension sequences were cloned into the pU6-tmpknot-GG-acceptor backbone (Addgene #174039), while the corresponding nicking sgRNAs were cloned into a nicking sgRNA backbone (Addgene #47108). To achieve higher editing efficiency, PEmax-P2A-hP53DD (Addgene #214084), epegRNA, and nicking sgRNA plasmids were delivered into iPSCs via electroporation, followed by single-cell cloning^63^. Target loci from individual clones were amplified by PCR and analyzed by Sanger sequencing to identify correctly edited clones. sgRNA sequences can be found in **Table S3.** Sequences for primers used for sanger sequencing can be found in **Table S4.**

#### Endothelial tube formation assay

Endothelial tube formation assay (in vitro angiogenesis) (Cell Biolabs, CBA-200) was performed according to manufacturer’s instructions. In brief, 1×10^5^ CD31+ EPCs were seeded onto ECM gel-coated wells of a 96 well plate. The EPCs were cultured in BVI media and allowed to form tubes for 4-6 hours.

#### Generation of FOXF1 LOF hESC lines

H1-AAVS-GFP hESCs^61^ were used to introduce a 650 base pair length deletion in the FOXF1 gene (transcript FOXF1-201) spanning exon 1 to intron 1-2 to generate FOXF1^−/−^. A ribonucleoprotein (RNP) complex was assembled by combining 20µg Alt-R^TM^ S.p. HiFi Cas9 Nuclease V3 (Cas9) (IDT) with 4ug of each sgRNA which are constituted with Nuclease-Free Duplex Buffer (IDT, 11-01-03-01). An additional 0.4 µL Duplex Buffer was added for a final assembled RNP complex volume of 6.4 µL. 1×10^6^ hESCs were electroporated with 6.4ul RNP complex with 100ul P3 primary cell solution (Lonza, V4XP-3032) using the Lonza 4D nucleofector ® (Lonza), with the pulse code CA-137. Electroporated cells were cultured in mTeSR^TM^1 (Stem Cell Technologies, 85850) supplemented with 10% CloneR^TM^ Supplement (Stem Cell Technologies, 05888) in Cultrex SCQ-coated wells for 24 hrs. Daily media changes with mTeSR1 supplemented with CloneR^TM^ were performed until the culture achieved 80% confluency. Then, ACCUTASE^TM^ (Stem Cell Technologies, 07920) was used to obtain a single cell suspension of transfected cells, and the cells were subsequently plated at clonal density with CloneR-supplemented mTeSR1. Colonies were then manually excised, expanded, and genotyped. Clones identified to contain an amplicon indicative of a putative bi-allelic and mono-allelic *FOXF1* deletion genotype were subjected to Sanger sequencing to further confirm the deletion genotype. Clones harboring the bi-allelic FOXF1 null genotype (FOXF1^−/−)^ were identified, expanded and cryopreserved.

A clone containing the exon 1 to intron 1-2 FOXF1 deletion on one allele and a 4bp deletion on the 2nd allele was selected to generate FOXF1^+/−^. The RNP complex was assembled using 16 µg sgRNA, 20 µg Cas9, 0.4 µL Duplex buffer, and 8 µg of a single-stranded Ultramer DNA donor template for the repair of the 4bp deletion. In addition to the WT *FOXF1* sequence, the donor template contained a silent mutation to facilitate genotyping, asymmetrical 5’- and 3’-homology arms, and phosphorothioate bonds on the final three 5’ and 3’ nucleotides. Clones harboring the corrected allele were identified, sequenced, expanded and cryopreserved and used subsequently as mono-allelic FOXF1 null (FOXF1^+/−^). sgRNA sequences and donor DNA template can be found in **Table S3.** Sequences for primers used for genotyping and sanger sequencing can be found in **Table S4.**

#### Imaging

Brightfield images were obtained using the EVOS M5000 imaging system (Invitrogen). Immunofluorescent images were acquired using the Nikon A1R inverted LUNV confocal microscope (Nikon). Images were analyzed and processed using Nikon’s nis-elements 5.20.

#### Immunofluorescence of cryosections

Fixed VOs were first incubated in 30% sucrose (Sigma-Aldrich, 50-188-2408) containing 0.1% sodium azide (Aqua Solutions, Inc, SPE844) overnight at 4°C. Then, cryoblocks were made by embedding VOs in Tissue-Plus™ O.C.T. compound (Fisher Scientific, 23-730-571). The cryoblocks were left at −20°C overnight before cryosectioning. 16um VO cryosections were obtained using the Leica CM3050 S Cryostat (Leica Biosystems). To perform immunofluorescence staining, sections were brought to RT and fixed again with 4% PFA in PBS (Thermo Scientific Chemicals, J61899.AK), for 15 mins at RT, then permeabilized using 0.5% Triton-X (Sigma-Aldrich, X100) in PBS for 15 mins at RT. Then, the sections were blocked consisting of 5% donkey serum (Jackson ImmunoResearch, 017-000-121) in PBS for 1h at RT. Then, sections were incubated with primary antibodies diluted in blocking buffer overnight at 4°C. The primary antibodies used: CD31 (1:500, R&D Systems, AF806), PDGFRβ (1:100, Cell Signaling Technology, 3169), FOXF1 (1:100, R&D Systems, AF4798), HAND1 (1:100, R&D Systems, AF3168), MIXL1 (1:100, Thermo Fisher Scientific, PA5-64903), and MECOM (1:100, Cell Signaling Technology, 2593). The next day, the sections were wash thrice with DPBS at RT for 5 min on the rocking shaker. The sections were then incubated with secondary antibodies diluted in blocking buffer for 1.5h at RT. Secondary antibodies used: Donkey anti-Rabbit IgG (H+L) Highly Cross-Adsorbed Secondary Antibody, Alexa Fluor™ 488 (Invitrogen, A-21206), Donkey anti-Rabbit IgG (H+L) ReadyProbes™ Secondary Antibody, Alexa Fluor™ 594 (Invitrogen, R37119), Donkey anti-Sheep IgG (H+L) Cross-Adsorbed Secondary Antibody, Alexa Fluor™ 647 (Invitrogen, A-21448), Donkey anti-Mouse IgG (H+L) ReadyProbes™ Secondary Antibody, Alexa Fluor™ 488 (Invitrogen, R37114), Donkey anti-Mouse IgG (H+L) ReadyProbes™ Secondary Antibody, Alexa Fluor™ 594 (Invitrogen, R37115). Then, sections were counter-stained with DAPI (Thermo Fisher Scientific, 62248) (1:1000 in PBS) for 15 mins at RT. Sections were subsequently washed thrice with DPBS at RT for 5 min on the rocking shaker and mounted using ProLong™ Gold Antifade Mountant (Invitrogen, P36930) before imaging.

#### LNP-mRNA delivery to vessel organoids

Lipid-nanoparticles containing *FOXF1*, *MECOM*, or *eGFP* mRNA (GenScript) are first incubated with FBS (Gibco, A5256701) in a 1:1 ratio for 30 mins at 37°C before adding to organoid cultures. For LNP-*FOXF1* experiments, 1ug of LNP-*FOXF1* were utilized. For LNP-*MECOM* experiments, 2ug of LNP-*MECOM* were utilized. Accordingly, equal amounts of LNP-*eGFP* were utilized. mRNA sequences used for those LNP formulations can be found in **Table S5**.

#### Magnetic isolation of CD31+ EPCs

CD31+ ECs were isolated using the human CD31 MicroBead Kit (Miltenyi Biotec, 130-091-935) per manufacturer’s recommendations.

#### Migration/ scratch assay

8×10^4^ CD31+ EPCs were seeded into 0.01% Bovine Collagen Type I (Sigma-Aldrich, C8919)-coated 35mm µ-Dish containing 2 well culture inserts (ibidi, 81176). The EPCs were cultured in BVI overnight. The next day, the BVI media was replaced, and the culture inserts were removed to promote cell migration.

#### Molecular dynamics simulations

The structure of FOXF1 bound to DNA was modeled using AlphaFold Server 3^64^, as no experimental crystal structure is currently available. Both DNA-bound (COMPLEX) and DNA-unbound (APO) conformations were generated for the wild-type FOXF1 protein. Two mutants (L56V and F85I) were modeled with Visual Molecular Dynamics package^65^, resulting in three structural models for each condition. The DNA sequence used in the simulations was taken from a previous crystallographic study employing a segment of the transthyretin promoter^66^. All-atom molecular dynamics simulations were performed using GROMACS 2024^67,68^ with the CHARMM36 force field^69^. Each system was placed in a cubic simulation box and solvated with water molecules described by the TIP3P model. Mg²⁺ and Cl^−^ ions were added to neutralize the system with a final Mg²⁺ concentration of 47. mM for the COMPLEX systems and 61. mM for the APO systems. Energy minimization was carried out using the steepest descent algorithm for 50,000 steps, with a convergence criterion of a maximum force below 1000 kJ mol⁻¹ nm⁻¹. The minimized systems were then equilibrated sequentially under NVT and NPT ensembles. Temperature equilibration was performed for 100 ps at 298 K using the velocity-rescale thermostat, followed by pressure equilibration for 100 ps at 1 bar using the C-rescale barostat. Production simulations consisted of five independent unbiased trajectories of 500 ns for each system with a time step of 2 fs, yielding a total aggregate simulation time of 12 μs. For all analyses, only the FOXF1 DNA-binding domain (residues 43–148) was considered^3^. Data frames were saved every 0.1 ns after discarding the first 20 ns of each trajectory as equilibration. Analysis code and molecular dynamics data is available at https://doi.org/10.5281/zenodo.20736501.

#### 10X single-nucleus Multiome (RNA+ATAC) profiling of vessel organoids

Day 3 and 5 VOs were dissociated into single cells prior to nuclei isolation. First, VOs were wash thrice with DPBS in 5ml tubes. Then, DPBS were aspirated and 2ml of 1X TrypLE™ Express (Gibco, 12604013) was added. The tubes were placed on a rocking shaker in the 37°C incubator for 5-10 minutes. Once single-cell suspension was attained, the digestion reaction was quenched with ice-cold KnockOut^TM^ DMEM/F12 (Gibco, 12660012) media containing 10% FBS (Gibco, A5256701). Cells were centrifuged at 300g for 5 mins at 4°C. Supernatant was removed, and cell pellet was resuspended in cold DPBS containing 2% FBS (Gibco, A5256701). Cell mixture was filtered through a 40um filter and centrifuged again at 300g for 5 mins at 4°c. Supernatant was removed and resuspended in cold DPBS containing 2% FBS (Gibco, A5256701) before performing nuclei isolation.

Nuclei was isolated from the single cell suspension using the Nuclei EZ prep kit (Sigma-Aldrich, nuc101) based on a protocol detailed by Bessonett et al., 2022^70^. Transfer cells into BSA (Miltenyi Biotec, 130-091-376)-coated Lobind Eppendorf tubes (Eppendorf, 0030108051). Spin cells down in a swing-bucket centrifuge at 300g for 5 mins (4°c). Remove supernatant and add 200ul EZ lysis buffer supplemented with 1mM dl-dithiothreitol (DTT) (Sigma-Aldrich, 646563) and 1U/μl of Ribolock RNase Inhibitor (Thermo scientific, eo0381). Leave cells on ice for 5 mins. Add 1ml of ice-cold wash buffer to quench the lysis process. Spin cells down immediately at 500g for 5 mins (4°c). The wash buffer consists of 10mM TRIS-HCl (Invitrogen, 15567027), 10mM NaCl (Invitrogen, AM9760G), 3mM MgCl_2_ (Invitrogen, AM9530G), 1% BSA (Miltenyi Biotec, 130-091-376), 1U/μl Ribolock RNase inhibitor (Thermo Scientific, eo0381). Remove the supernatant and resuspend nuclei in 200ul 1x Nuclei Buffer (10x Genomics, 2000207). Filter nuclei samples using 40μm Flowmi^TM^ cell strainers (bel-art, h136800040). 10, 000 nuclei were used for chromium next GEM single cell Multiome ATAC + gene expression sequencing (10x Genomics, PN-1000282).

#### 10X single-nucleus Multiome (RNA+ATAC) data analysis

All datasets were aligned using 10x Genomics cellranger-arc 2.0.2^71^ against the GRCh38 “2020-A” genomic reference. All subsequent analyses were performed using the R packages Seurat version 5.4.0^72^ and Signac version 1.14.0^73^. The RNA assay of each dataset was corrected for ambient RNA using the package SoupX version 1.6.2^74^. Ribosomal and mitochondrial genes were removed from the counts, SCTransform was used for the normalization and scaling, and standard PCA, clustering, and UMAP steps are also performed. Lastly, doublets are predicted and removed using DoubletFinder version 2.0.6^75^. The ATAC assay is added using a unified set of peaks generated from all datasets, and the resulting multiomic dataset is further filtered based on nCount_RNA between 1000 and 25000, nCount_ATAC between 1000 and 100000, nucleosome signal <2, and TSS enrichment >2. For RNA integration, objects are combined and processed with the same SCTransform approach, then CCA was implemented. For ATAC integration, TFIDF normalization and SVD reduction are performed, then embeddings from reciprocal LSI are calculated and integrated. To visualize the assays together, we also generate a UMAP based on the multimodal weighted nearest neighbors.

Figure generation, finding marker genes, and cell annotation were performed through Mona (https://github.com/ZornLab/Mona), an interactive visualization tool. Further downstream analysis of ATAC data including differential accessibility, coverage plots, and linking peaks to genes were done using Signac. Importantly, the MAST^76^ model was used when calculating differential genes, whereas logistic regression was used for differential peaks. For both, default values of 0.1 for fold change and 0.01 for min.pct were used as cutoffs, as well as a 0.05 adjusted p-value cutoff. FOXF1 motif scanning within the differentially accessible peaks was performed using FIMO version 5.5.5^77^. Once cell populations were identified, cell-cell interactions were inferred and visualized with the package CellChat version 2.1.2^36^. Scoring of gene sets was performed using UCell version 2.14^78^. Codes associated with single-nucleus Multiome data analysis are available at https://github.com/ZornLab/vessel_organoid_analysis.

#### Pathogenicity predictions

AlphaMissense^26^ and PolyPhen-2^27^ were used to predict the effects of *FOXF1* missense mutations on protein function. In AlphaMissense, scores close to 1 denote ‘likely pathogenic’ while scores close to 0 are ‘likely benign’. In PolyPhen-2, scores close to 1 denote ‘probably damaging’ while scores close to 0 are ‘benign’.

#### Quantitative Reverse-transcription (qRT-PCR)

The method used to perform qRT-PCR was previously-described^79,80^. Briefly, total RNA from VO samples were isolated and purified using the Quick-RNA Microprep kit (Zymo Research, R1050). RNA was then converted to cDNA using the High-Capacity cDNA Reverse Transcription Kit (Applied Biosystems, 4374967) according to manufacturer’s recommendations. 5ng of cDNA and PowerUp^TM^ SYBR^TM^ Green Master Mix (Applied Biosystems, A25742) was used to perform qPCR. Sequences of primers used can be found in **Table S2**.

#### Western Blot

Protein from VOs was extracted using Pierce RIPA Buffer (Thermo Scientific, 89901) supplemented with 1X of Halt™ Protease and Phosphatase Inhibitor Cocktail (Thermo Scientific, 78440). Concentration of protein lysates was determined using the Pierce^TM^ BCA assay kit (Thermo Scientific, 23225). 1X non-reducing Laemmli SDS sample buffer (Thermo Scientific, J60660.AD) was added to protein samples and denatured at 70°c for 10mins. Protein samples were then separated using the 4-12% Bolt^TM^ Bis-Tris Plus Mini Protein Gels (Invitrogen, NW04122BOX) and transferred onto nitrocellulose membranes (Invitrogen, PB3210) using the Power Blotter semi-dry transfer system (Invitrogen, PB0012). Then, the membranes were blocked with 5% non-fat milk in TBST. TBST is made up of 0.1% Tween-20 (Sigma-Aldrich, P2287) in 1X TBS (Invitrogen, J60764.K2) for 1h at RT before overnight incubation with primary antibodies diluted in 5% non-fat milk in TBST. The following primary antibodies were used: FOXF1 (1:100, Abcam, ab168383), FLK1 (1:1000, R&D Sytems, AF357), beta-ACTIN (HRP-conjugated) (1:1000, Cell Signaling Technology, 5125). The next day, the membranes were washed three times with TBST for 5-10 mins at RT before incubating with secondary antibodies. Secondary antibodies used include mouse anti-rabbit IgG-HRP (1:1000, Santa Cruz Biotechnology, sc-2357) and rabbit anti-goat (1:1000) diluted in 5% low fat milk in TBST. Subsequently, the membranes were washed three times with TBST for 5-10 mins at RT before adding the SuperSignal^TM^ West Pico PLUS chemiluminescent substrate (Thermo Scientific, 34580) to measure chemiluminescence signals. Chemiluminescence images were acquired using the ChemiDoc MP Imaging System (BioRad).

#### Whole-organoid immunofluorescence

mVOs were fixed overnight at 4°C in 4% PFA in PBS (Thermo Scientific Chemicals, J61899.AK). After fixation, mVOs were washed thrice with DPBS at RT. Fixed mVOs were incubated in blocking buffer consisting of 3% FBS (Gibco, A5256701), 1% BSA (Miltenyi Biotec, 130-091-376), 0.5% Triton-X (Sigma Aldrich, X100), 0.5% Tween-20 (Sigma Aldrich, P2287), and 0.01% sodium deoxycholate (bio-world, 40430018-1) solution. at RT for 4h on a rocking shaker. Subsequently, mVOs were incubated in primary antibodies diluted in blocking buffer overnight at 4°C. The primary antibodies used: CD31 (1:100, RnD Systems, AF806), PDGFRβ (1:100, Cell Signaling Technology, 3169), and FOXF1 (1:100, RnD Systems, AF4798). The next day, mVOs were washed thrice with PBS-T (DPBS with 0.05% Tween-20) at RT before incubation with secondary antibodies and DAPI (Thermo Scientific, 62248) diluted in blocking buffer for 4h at RT on the rocking shaker. The dilution of secondary antibodies and DAPI were 1:200 and 1:300 respectively. Then, the mVOs were washed thrice with PBS-T at RT and mounted in an µ-Slide 8 well glass-bottom chamber slides (ibidi, 80826) immersed in RapiClear CS optical clearing solution (Sunjin Lab, RCCS001). Confocal microscopy was performed using the Nikon A1R Inverted Confocal Microscope (Nikon).

### QUANTIFICATION AND STATISTICAL ANALYSIS

Vessel parameters such as average tube length and branch counts were quantified using FastTrack AI™ (MetaVi Labs). For all other image quantification (i.e. western blot bands, gastruloid circularity), ImageJ was used. Additional ImageJ plugins used: Tube formation assay images were quantified using the ‘Angiogenesis Analyzer’ plugin (http://image.bio.methods.free.fr/ImageJ/?Angiogenesis-Analyzer-for-ImageJ) and the migration assay images were quantified using the ‘Wound Healing Size Tool’ plugin^81^ (https://github.com/AlejandraArnedo/Wound-healing-size-tool/wiki#wound-healing-size-tool). Gene ontology analysis was performed using Enrichr^82^ (https://maayanlab.cloud/Enrichr/). Volcano plots and GO enrichment bubble plots were generated using SRplot^83^ (https://www.bioinformatics.com.cn/en).

Statistical analyses were performed using GraphPad Prism 10.0. All data represents mean ± S.D. Statistical significance were determined via parametric tests - paired two-tailed t-test (two groups), one-way ANOVA coupled with post-hoc Tukey’s test (more than 2 groups), or two-way ANOVA coupled with post-hoc Bonferroni test (more than 2 groups across multiple conditions). p-values <0.05 were considered statistically significant. To calculate the p-value of whether the differential genes are overrepresented in the differential peaks, the hypergeometric test was used.

Quantification of molecular dynamics metrices is as follows:

#### Root-mean-square deviation (RMSD)

RMSD quantifies the structural deviation of atoms from a reference conformation and is commonly used to monitor equilibration in Cartesian space. In this work, we calculated the RMSD for the backbone atoms of DNA binding domain of FOXF1.

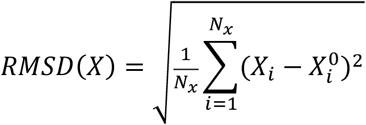

where RMSD(X) RMSD of conformation X, *X_i_* represents the Cartesian coordinate of atom *i* in conformation X, *N_X_* is the number of atoms included in the calculation, and 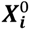 denotes the corresponding coordinate of atom *i* in the reference structure at t=0 ns.

#### Root-mean-square fluctuation (RMSF)

RMSF measures the extent of positional fluctuations of residues around their average structure and provides a residue-level description of protein flexibility. RMSF values were computed for the Cα atom of each residue for each data frame as

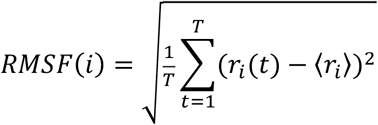

where ***r_i_(t)*** is the Cartesian coordinate of atom *i* at time t, 〈***r_i_***〉 is its average position over the trajectory, and T is the number of data frames.

#### Analysis of hydrogen bonds

Hydrogen bonds between the FOXF1 DNA-binding domain and double-stranded DNA were calculated using the gmx hbond module of GROMACS. A geometric criterion of a donor–acceptor distance ≤ 0.35 nm (3.5 Å) and a hydrogen–donor–acceptor angle ≤ 30°.

#### Analysis of solvent-accessible surface area

The solvent-accessible surface area quantifies the exposure of each residue of a protein or nucleic acid to the solvent. We calculate the solvent-accessible area of each residue using the Python package mdtraj^84^ which uses the Shrake-Rupley algorithm^85^ and then sum the values for the protein and nucleic acid separately to derive a total value for each species.

#### Analysis of salt bridges

Salt bridges are an electrostatic interaction between a basic atom within a residue such as an oxygen on glutamate or phosphate and an acidic atom within a residue such as a nitrogen in lysine^86^. Salt bridges between the FOXF1 protein and DNA strand were computed for each data frame with the Visual Molecular Dynamics^65^ software using the criterion that the distance between the negative and positive atoms must fall within 4 Ångstroms in at least one data frame and the distance between the center of masses of the two residues must fall within 10 Ångstroms for at least 50% of the simulation time.

## Supplemental Information

### SUPPLEMENTARY FIGURE LEGENDS

**Figure S1.**
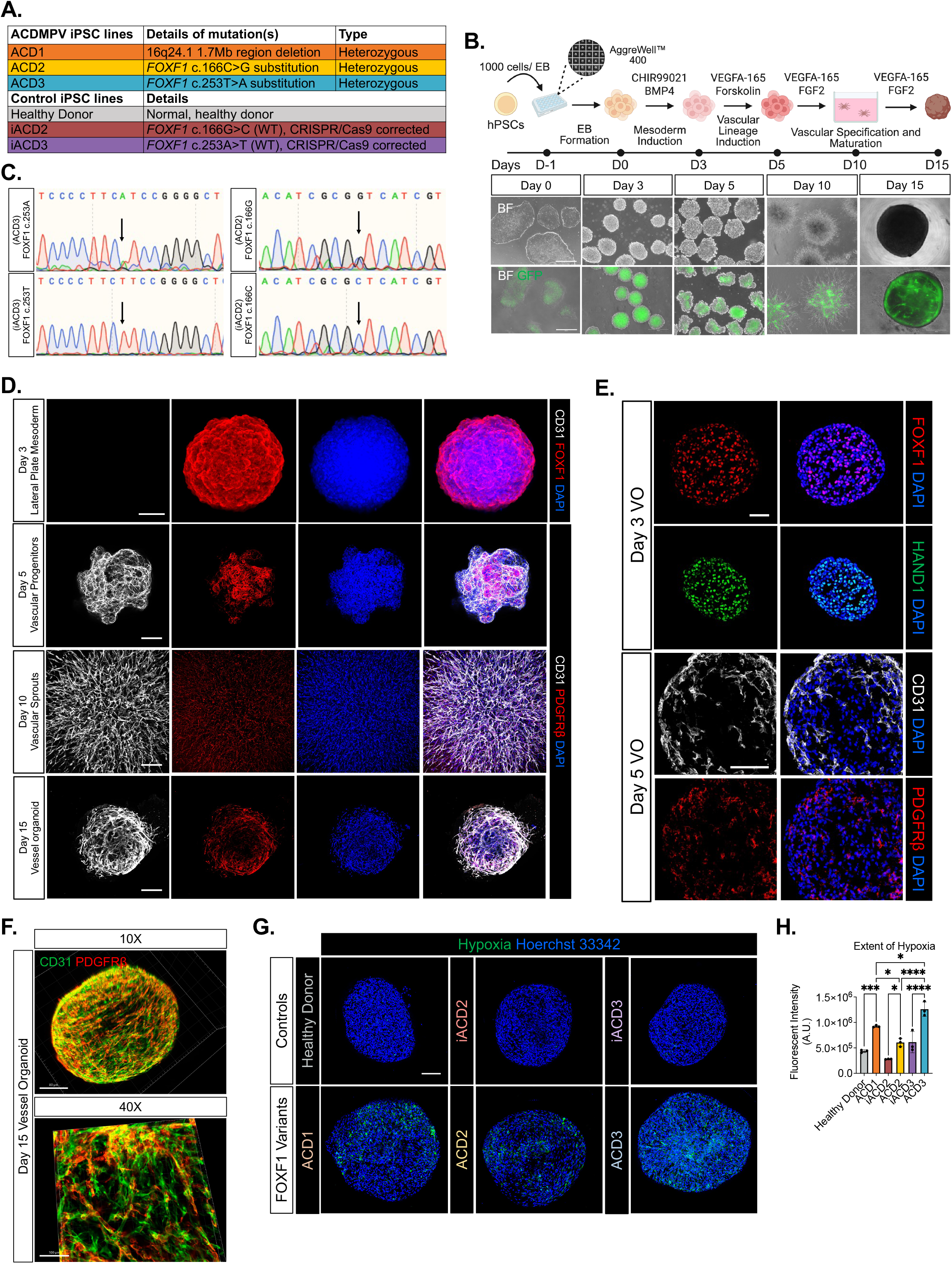
Generation and characterization of human iPSC-derived vessel organoids from ACDMPV patient and isogenic control lines, related to Figure 1. (A) Table summarizing the ACDMPV and control iPSC lines. (B) Top: Schematic diagram illustrating the differentiation of hiPSCs into VOs. Bottom: Representative brightfield and fluorescence images showing the morphologies of VO throughout different stages of VO differentiation. Scale bar= 500um. (C) Sanger sequencing confirming the correction of the c.166C>G and c.253T>A variants in the isogenic control lines iACD2 and iACD3, respectively. (D) Representative whole-organoid immunofluorescence images of D3, D5, D10, and D15 VOs. Nuclei are counterstained with DAPI (blue). Scale bar=100um. (E) Representative immunofluorescence images of D3 and D5 VO cryosections. Nuclei are counterstained with DAPI (blue). Scale bar=100um. (F) Representative 3D-rendered whole-organoid immunofluorescence image of a D15 VO showing CD31 (green) and PDGFRβ (red). Top: 10x magnification, scale bar=80um. Bottom: 40x magnification, scale bar=100um. (G) Representative whole-organoid live immunofluorescence images showing extent of hypoxia (green). Nuclei are counterstained with Hoerchst 33342 (blue). Scale bar=100um. (H) Quantification of extent of hypoxia (mean ± S.D., n=3 organoids, *p<0.05, **p<0.01, ****p<0.0001 by one-way ANOVA followed by Tukey’s test).

**Figure S2.**
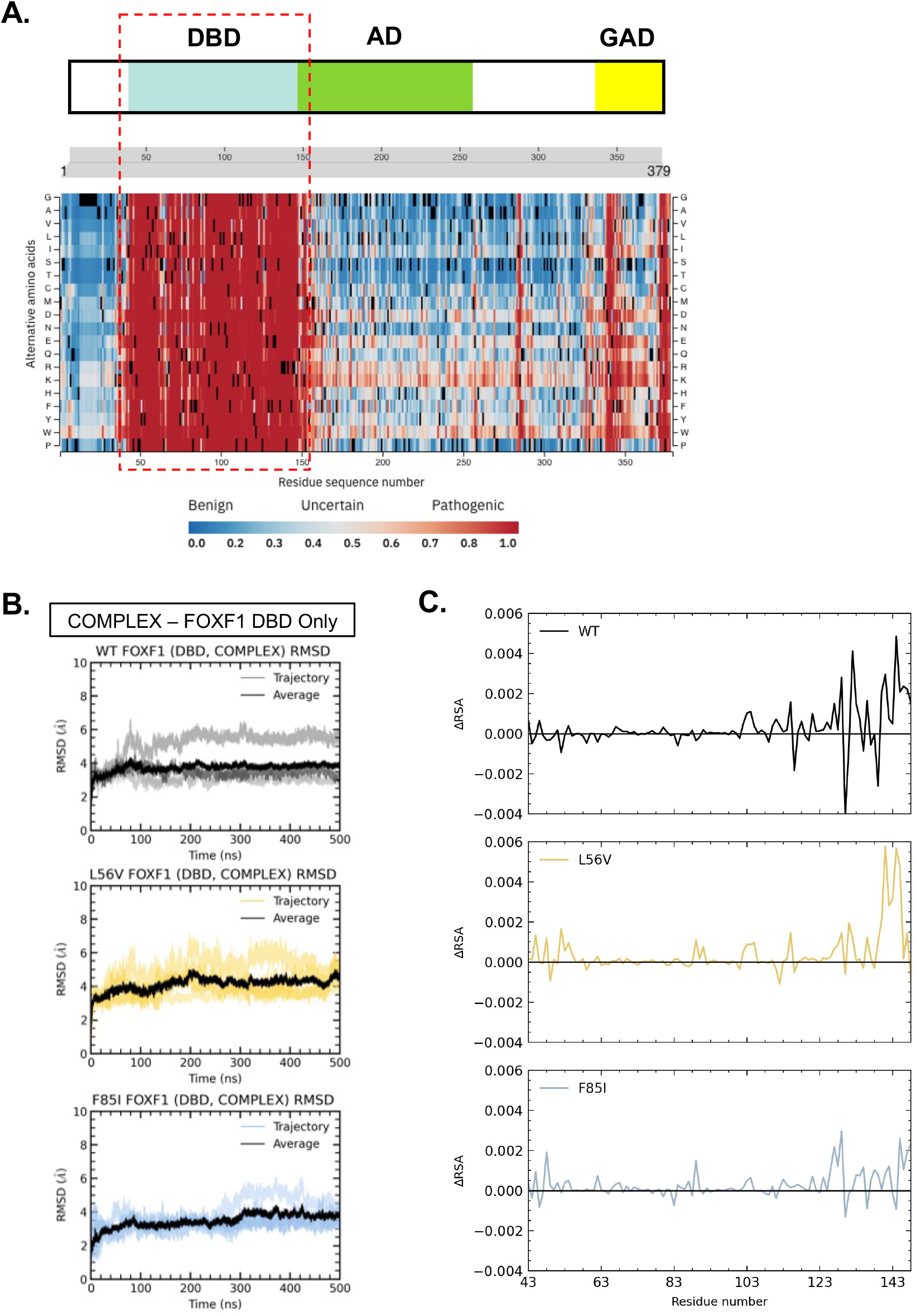
Computational analyses predict pathogenic effects of FOXF1 variants, related to Figure 3. (A) AlphaMissense pathogenicity heatmap of the FOXF1 illustrating that amino acid substitutions in the DBD region are predicted to be highly pathogenic (red). (B) RMSD (Å) plot of dsDNA-bound/ COMPLEX WT, L56V, and F85I FOXF1 proteins (DBD only) over simulation time of 500ns. (C) Residue-wise difference in relative solvent accessibility (RSA) of wild-type and variant protein conformations with large and small total SASA. Positive ΔRSA values indicate increased solvent exposure in the large-SASA structures.

**Figure S3.**
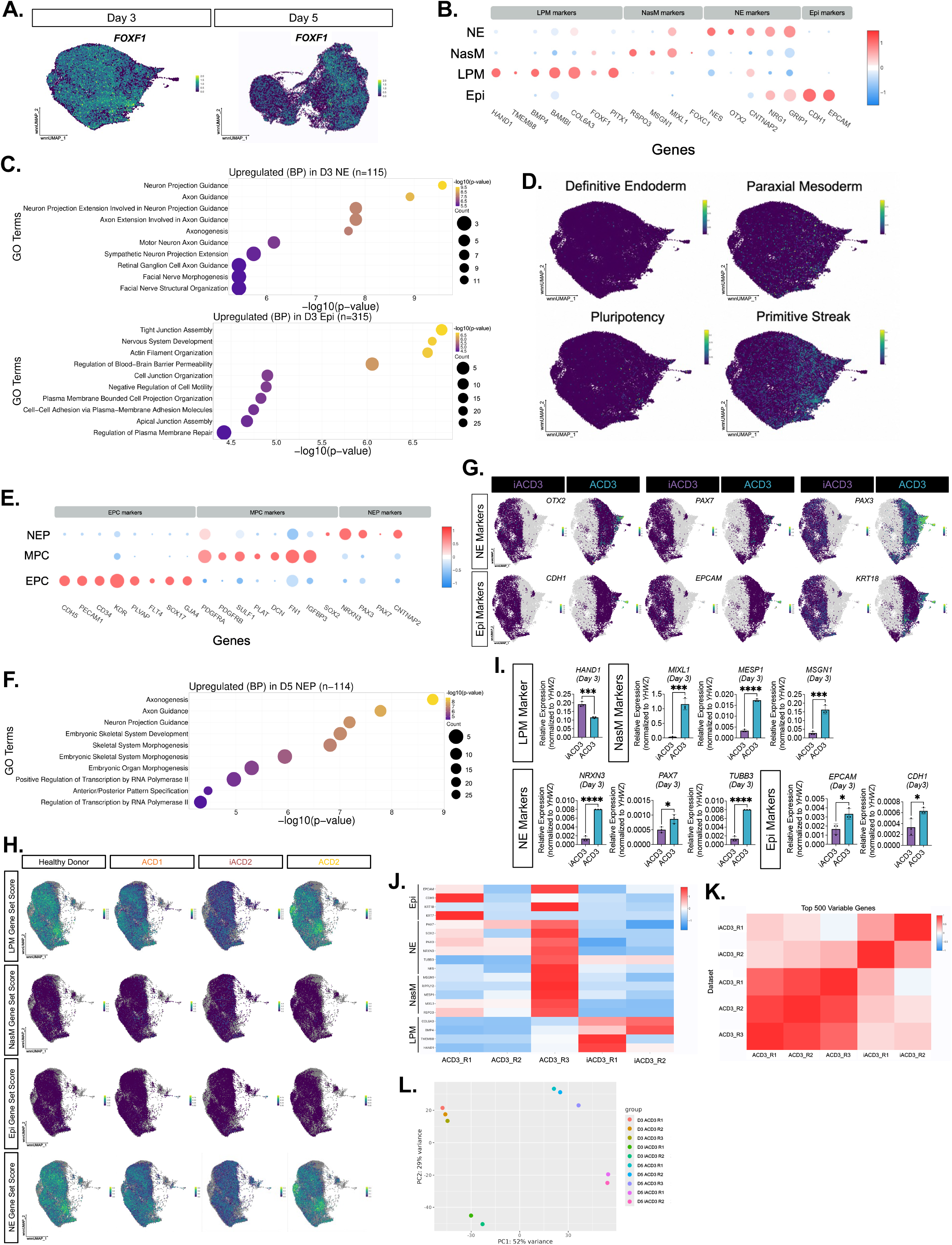
Single-nucleus multiomic analysis revealed developmental abnormalities unique to ACD3, related to Figure 4. (A) UMAP visualization of *FOXF1* expression in D3 and D5 VOs. (B) Dot plot illustrating representative LPM, NasM, NE, and Epi gene markers in D3 VOs. (C) GO analysis of top upregulated DEGs (BP) in D3 ACD3 NE compared to D3 ACD3 LPM, NasM, and Epi cells (Top) and D3 ACD3 Epi compared to D3 LPM, NasM, and NE cells (Bottom). (D) UMAP visualization of gene set scores for definitive endoderm, paraxial mesoderm, pluripotency, and primitive streak in D3 VOs. (E) Dot plot illustrating representative EPC, MPC, and NEP gene markers in D5 VOs. (F) GO analysis of top upregulated DEGs (BP) in D5 ACD3-NEP when compared to iACD3-NEP. (G) Feature plots showing expression of NE and Epi gene markers in D3 iACD3 and ACD3-VOs. (H) Feature plots illustrating LPM, NasM, Epi, and NE gene set scores in D3 Healthy Donor, ACD1, iACD2, and ACD2-VOs. (I) Bulk mRNA expression of LPM, NasM, NE, and Epi gene markers in D3 ACD3-VOs compared to D3 iACD3-VOs (mean ± SD, n= 3 biological replicates, *p<0.05, ***p<0.001, ****p<0.0001 by paired student’s t-test). (J) Heatmap showing expression of key LPM, NasM, NE, and Epi gene markers across the two iACD3 biological snMultiome-seq replicates and three ACD3 biological snMultiome-seq replicates. (K) Heatmap showing expression of top 500 variable genes across the two iACD3 biological snMultiome-seq replicates and three ACD3 biological snMultiome-seq replicates. (L) PCA plot showing transcriptomic similarity across the different biological replicates.

**Figure S4.**
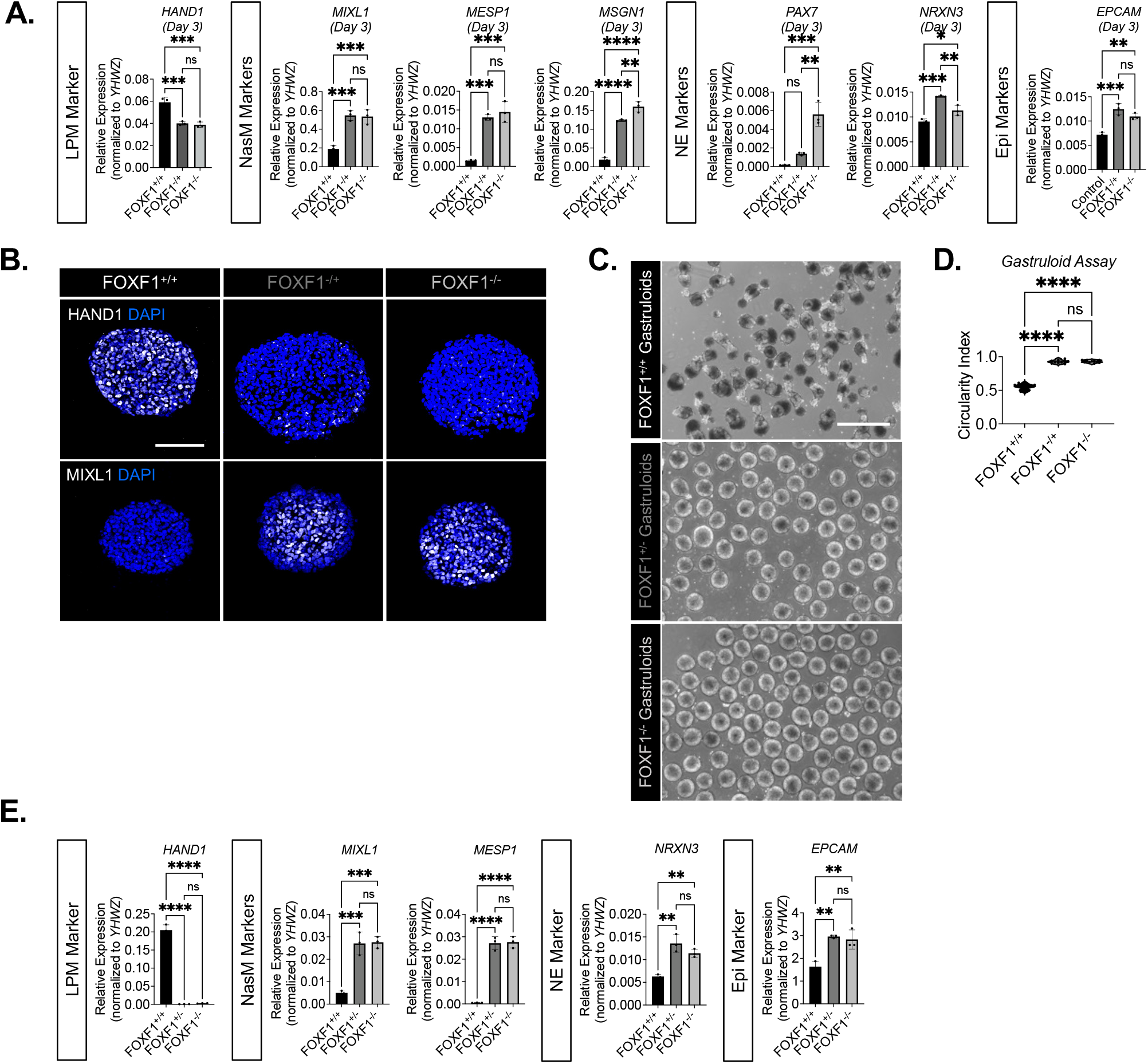
Developmental abnormalities observed in ACD3-VOs were mirrored in FOXF1-LOF-VOs, related to Figure 4. (A) Bulk mRNA expression of LPM, NasM, NE, and Epi gene markers in D3 FOXF1-LOF VOs compared to controls (mean ± SD, n=3 biological replicates, *p<0.05, **p<0.01, ***p<0.001, ****p<0.0001 by one-way ANOVA followed by Tukey’s test). (B) Representative immunofluorescence images of D3 VO cryosections stained with HAND1 and MIXL1. Nuclei are counterstained with DAPI (blue). Scale bar=100um. (C) Representative brightfield images of control and FOXF1-LOF gastruloids. Scale bar=500um. (D) Quantification of gastruloid circularity (mean ± SD, n=10 organoids from 3 biological replicates, ****p<0.0001 by one-way ANOVA followed by Tukey’s test). (E) Bulk mRNA expression of LPM, NasM, NE, and Epi gene markers in FOXF1-LOF gastruloids compared to controls (mean ± SD, n=3 biological replicates, ****p<0.0001 by one-way ANOVA followed by Tukey’s test).

**Figure S5.**
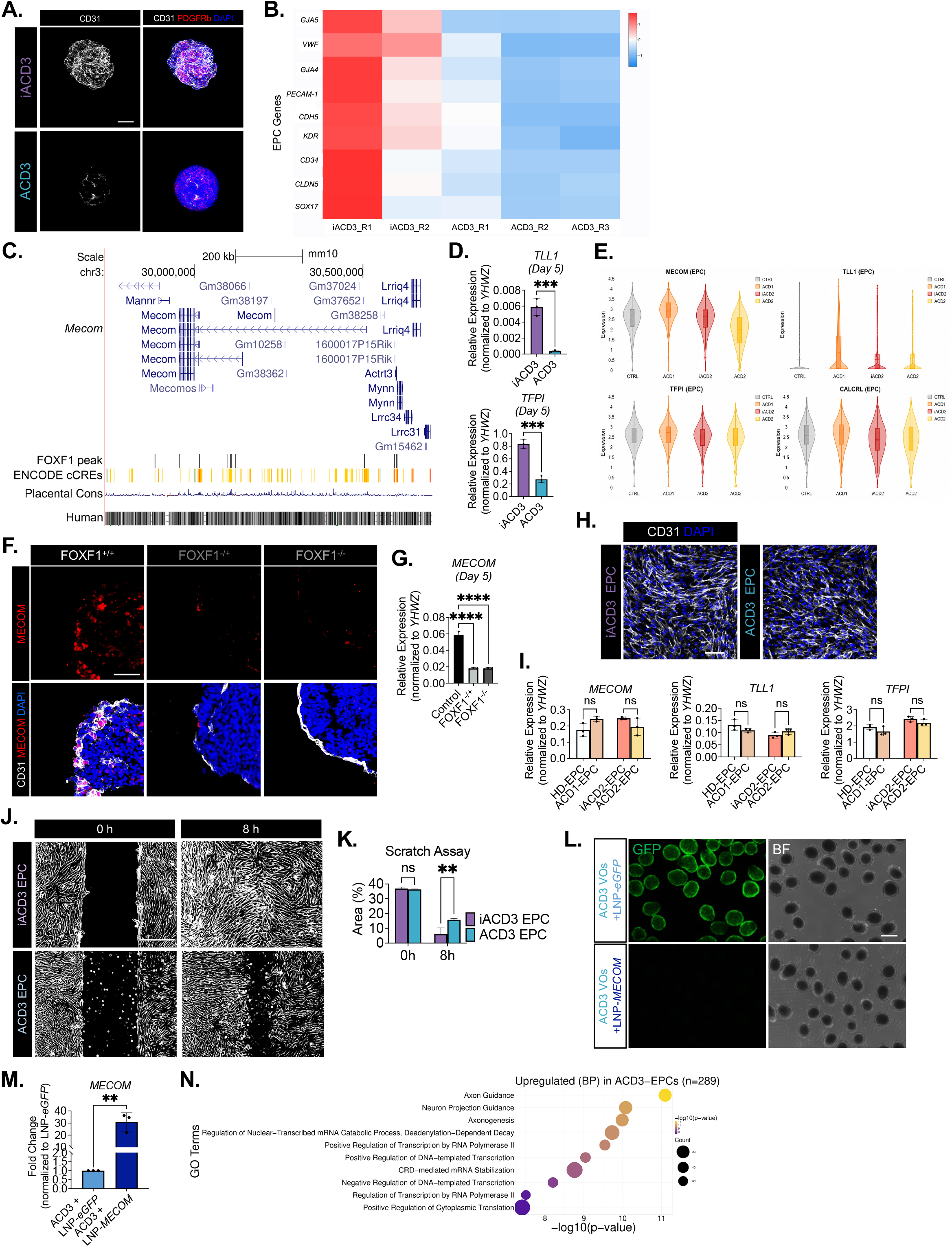
Loss of MECOM expression is associated with perturbed ACD3-EPC identity and function, related to Figure 5. (A) Representative whole-organoid immunofluorescence images showing CD31 (white), PDGFRβ (red). Nuclei are counterstained with DAPI (blue). Scale bar represents= 200um. (B) Heatmap showing expression of EPC genes across the two iACD3 biological D5 snMultiome-seq replicates and three ACD3 biological D5 snMultiome-seq replicates. (C) FOXF1 binding motifs in *MECOM* from FOXF1 ChIP-seq performed in mouse fetal lung ECs. (D) Bulk mRNA expression of *TLL1* and *TFPI* in D5 ACD3-VOs compared to D5 iACD3-VOs (mean ± SD, n= 3 biological replicates, ***p<0.001 by paired student’s t-test). (E) Violin plot showing expression of *MECOM*, *TLL1, TFPI,* and *CALCRL* in D5 Healthy Donor, ACD1, iACD2, and ACD2-EPCs. (F) Representative immunofluorescence images of D5 VO cryosections showing MECOM (red) and CD31 (white). Nuclei are counterstained with DAPI (blue). Scale bar=50um. (G) Bulk mRNA expression of *MECOM* in D5 FOXF1-LOF VOs compared to controls (mean ± SD, n=3 biological replicates, ****p<0.0001 by one-way ANOVA followed by Tukey’s test). (H) Representative immunofluorescence images of EPCs isolated from D5 iACD3 and ACD3-VOs stained for CD31 (gray). Nuclei are counterstained with DAPI (blue). (I) Bulk mRNA expression of *MECOM*, *TLL1,* and *TFPI* in EPCs isolated from D5 Healthy Donor, ACD1, iACD2, and ACD2-VOs (mean ± SD, n= 3 biological replicates, n.s.= p≥0.05 by paired student’s t-test). (J) Representative brightfield images illustrating migration of EPCs at 0 and 8h. Scale bar=400um. (K) Quantification of area of scratch at 0 and 8h (mean ± SD, n= 3 biological replicates, ***p<0.001 by student’s t-test). (L) Representative brightfield and fluorescence images of LNP-*eGFP* and LNP-*MECOM* treated D5 ACD3-VOs. Scale bar=500um. (M) Bulk mRNA expression of *MECOM* in LNP-*eGFP* and LNP-*MECOM* treated D5 ACD3-VOs. (mean ± SD, n= 3 biological replicates, **p<0.01 by paired student’s t-test). (N) GO analysis of top upregulated DEGs (BP) in D5 ACD3-EPCs compared to iACD3-EPCs.

**Figure S6.**
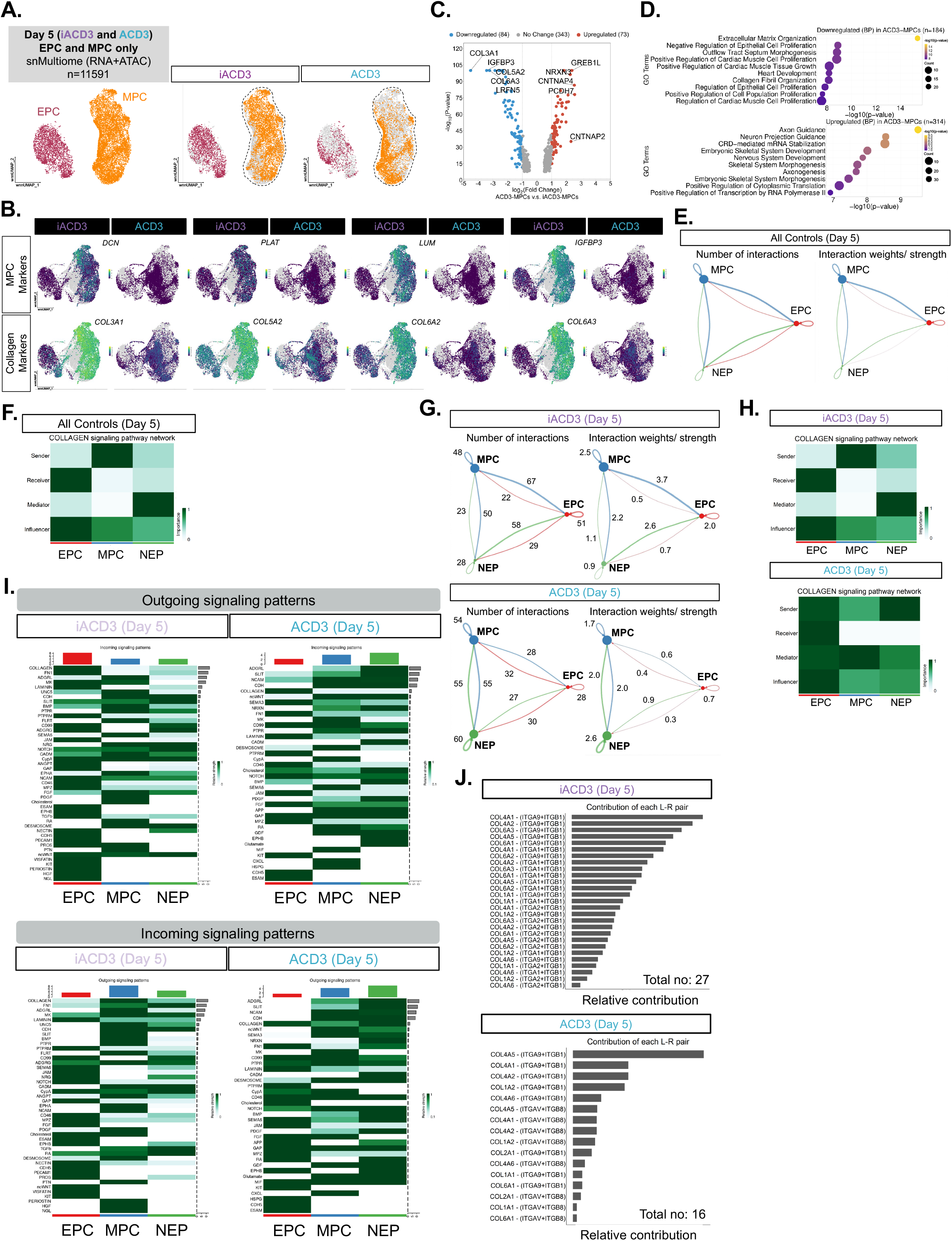
Disrupted collagen signaling network observed in ACD3-MPCs. (A) UMAP visualization of D5 iACD3 and ACD3 VOs, EPCs and MPCs only (n=11591). (B) Feature plots showing the expression of MPC markers and collagen genes in D5 ACD3-VOs compared to iACD3-VOs. (C) Volcano plot showing top DEGs in ACD3-MPCs compared to iACD3-MPCs. (D) GO term (BP) analysis downregulated (top) and upregulated (bottom) DEGs in ACD3-MPCs compared to iACD3-MPCs. (E) Number and strength of interactions between cell clusters in all D5 controls. (F) Heatmap showing dynamics of collagen signaling network among each cell type in all D5 controls (healthy donor, iACD2, and iACD3). (G) Cell-cell interaction analysis showing number of interactions and interaction strength/ weight across cell types in D5 iACD3 and ACD3-VOs. (H) Heatmap dynamics of collagen signaling network among each cell type in D5 iACD3 and ACD3-VOs. (I) Heatmap illustrating incoming and outgoing signaling patterns in D5 iACD3-VOs and ACD3-VOs. (J) Ligand–receptor interactions within the collagen signaling pathway network in D5 iACD3-VOs and D5 ACD3-VOs.

**Figure S7.**
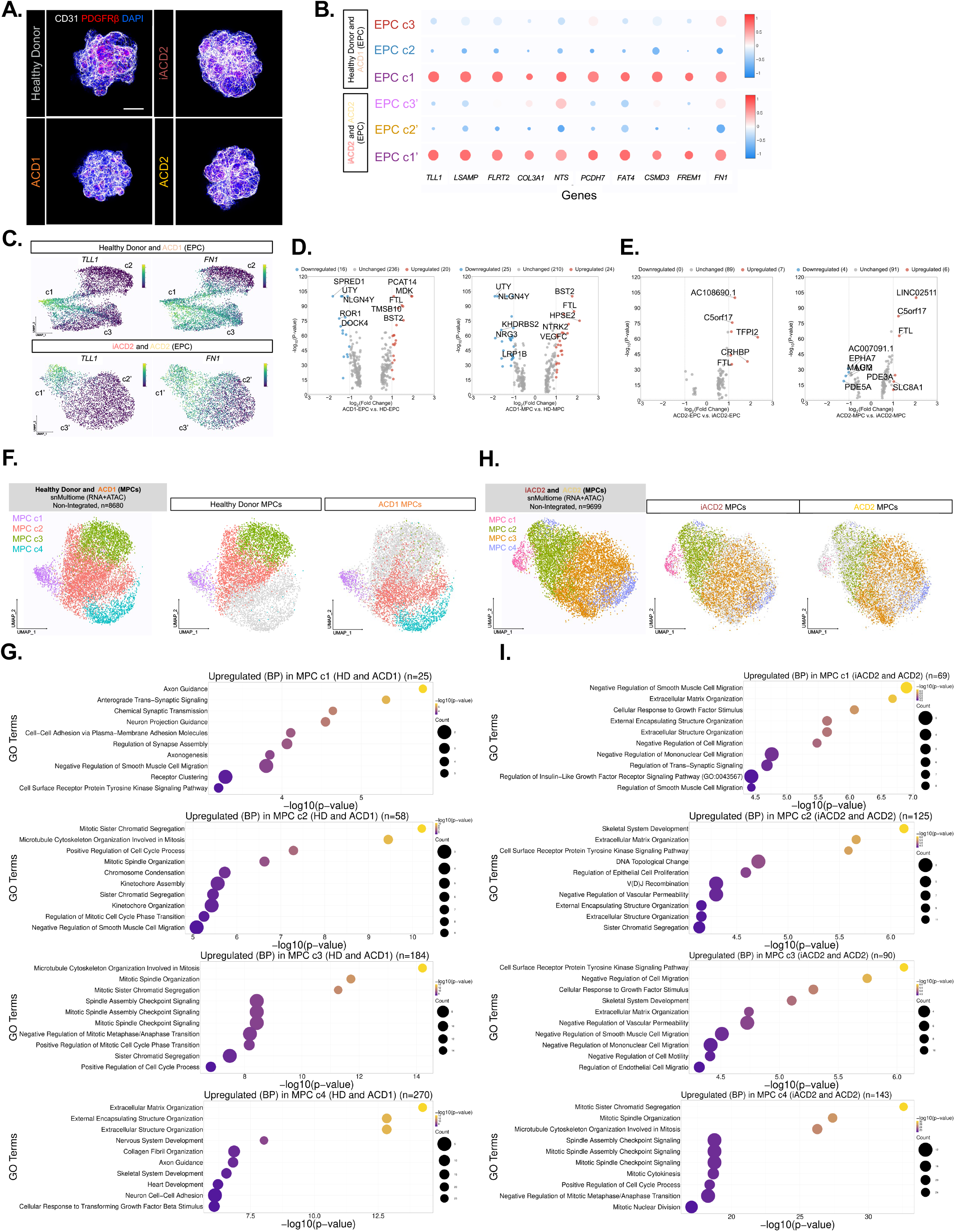
Transcriptomic and cell population changes in D5 ACD1 and ACD2-MPCs. (A) Representative whole-organoid immunofluorescence images showing CD31 (white), PDGFRβ (red). Nuclei are counterstained with DAPI (blue). Scale bar represents= 200um. (B) Left: Volcano plot showing DEGs in ACD1-EPCs compared to Healthy Donor-EPCs. Right: Volcano plot showing DEGs in ACD1-MPCs compared to Healthy Donor-MPCs. (C) Dot plot illustrating expression of representative genes of EPC c1 (Healthy Donor and ACD1 EPCs) and EPC c1’ (iACD2 and ACD2 EPCs). (D) Feature plot showing expression of *TLL1* and *FN1* in EPC c1 (Healthy Donor and ACD1 EPCs) and EPC c1’ (iACD2 and ACD2 EPCs) populations. (E) Left: Volcano plot showing DEGs in ACD2-EPCs compared to iACD2-EPCs. Right: Volcano plot showing DEGs in ACD2-MPCs compared to iACD2-MPCs. (F) UMAP visualization of MPC subclusters Healthy Donor and ACD1-MPCs (n=8680). Clustering was performed using the Louvain algorithm at a resolution of 0.8. (G) GO analysis (BP) of DEGs in each MPC subcluster relative to the other clusters. (H) UMAP visualization of MPC subclusters iACD2 and ACD2-MPCs (n=9699). Clustering was performed using the Louvain algorithm at a resolution of 0.8. (I) GO analysis (BP) of DEGs in each MPC subcluster relative to the other clusters.

**Figure S8.**
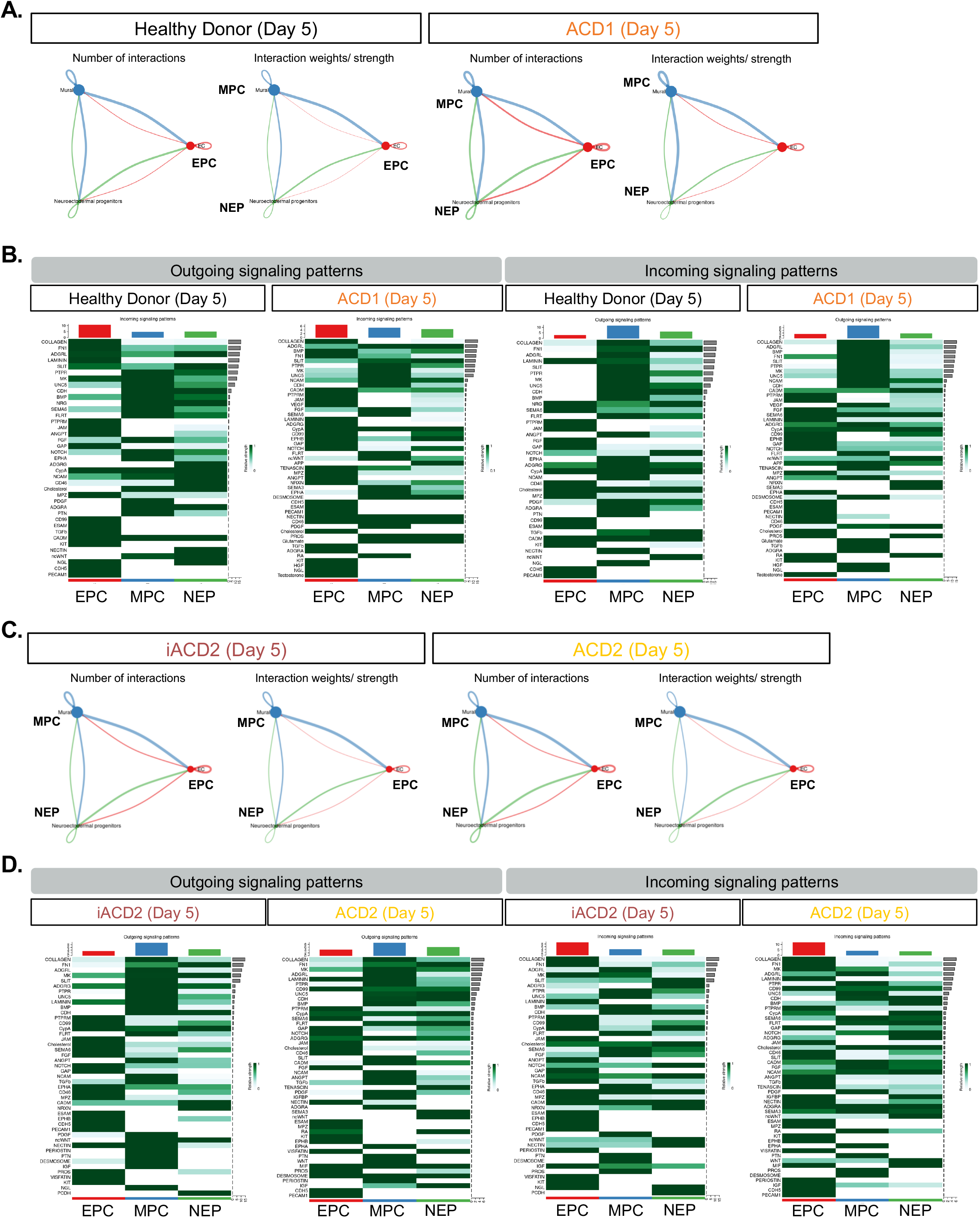
Altered in cell-cell communication in day 5 ACD1 and ACD2-VOs, related to Figure 6. (A) Number and strength of interactions between cell clusters in D5 Healthy Donor and ACD1-VOs. (B) Left: Heatmap illustrating outgoing signaling patterns in D5 HD-VOs and ACD1-VOs. Right: Heatmap illustrating incoming signaling patterns in D5 HD-VOs and ACD1-VOs. (C) Number and strength of interactions between cell clusters in D5 iACD2 and ACD2-VOs. (D) Left: Heatmap illustrating outgoing signaling patterns in D5 iACD2-VOs and ACD2-VOs. Right: Heatmap illustrating incoming signaling patterns in D5 iACD2-VOs and ACD2-VOs.

**Figure S9.**
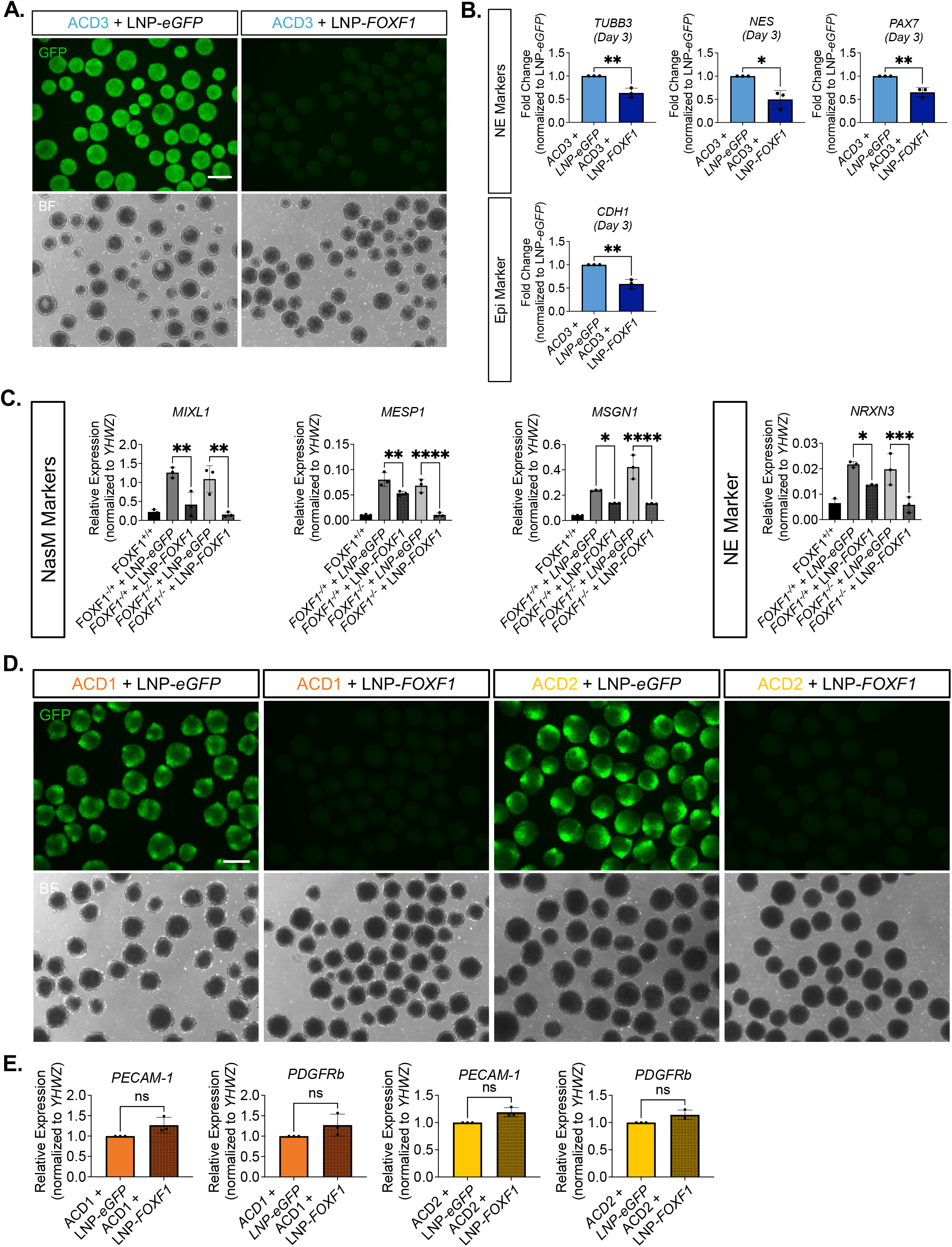
Further validation of LNP-mRNA delivery to VOs, related to Figure 7. (A) Representative brightfield and fluorescence images of LNP*-eGFP* and LNP-*FOXF1* treated D3 ACD3-VOs. Scale bar=500um. (B) Bulk mRNA expression of NE and Epi gene markers in D0 LNP*-eGFP* and LNP-*FOXF1* treated D3 ACD3-VOs (mean ± SD, n= 3 biological replicates, *p<0.05, **p<0.01 by paired student’s t-test). (C) Bulk mRNA expression of NasM and NE gene markers in D0 LNP*-eGFP* and LNP-*FOXF1* treated D3 FOXF1-LOF VOs (mean ± SD, n= 3 biological replicates, *p<0.05, **p<0.01, ****p<0.0001 by paired student’s t-test). (D) Representative brightfield and fluorescence images of LNP*-eGFP* and LNP-*FOXF1* treated D5 ACD1 and ACD2-VOs. Scale bar=500um. (E) Bulk mRNA expression of *PECAM1* and *PDGFRβ* in D5 LNP*-eGFP* and LNP-*FOXF1* treated D6 ACD3-VOs (mean ± SD, n= 3 biological replicates, n.s.= p≥0.05 by paired student’s t-test).

**Table S1.** ACDMPV Patient Information.

| Patient | Sex | Age | Ethnicity | Genotype | Co-anomalies | Age of mortality |
| --- | --- | --- | --- | --- | --- | --- |
| ACD1 | Female | Newborn | Unknown | 1.7 Mb deletion which included the FOXF1 locus (hg19 chr16:85,063,102-86,721,852). | Atrial septal defect, gall bladder agenesis, bicornuate uterus, bilateral renal dysplasia, ureteral stenosis, laryngeal stenosis, kidney dysplasia, abnormal right bronchial branching | At birth |
| ACD2 | Male | Newborn | Caucasian | *FOXF1* c.166C>G | Hirschsprung disease | 35 days |
| ACD3 | Male | Newborn | Caucasian | *FOXF1* c.253T>A | Atrial septal defect, ventricle septal defect, gall bladder agenesis, duodenal atresia, anal atresia, intestinal malrotation | 17 days |

**Table S2.**
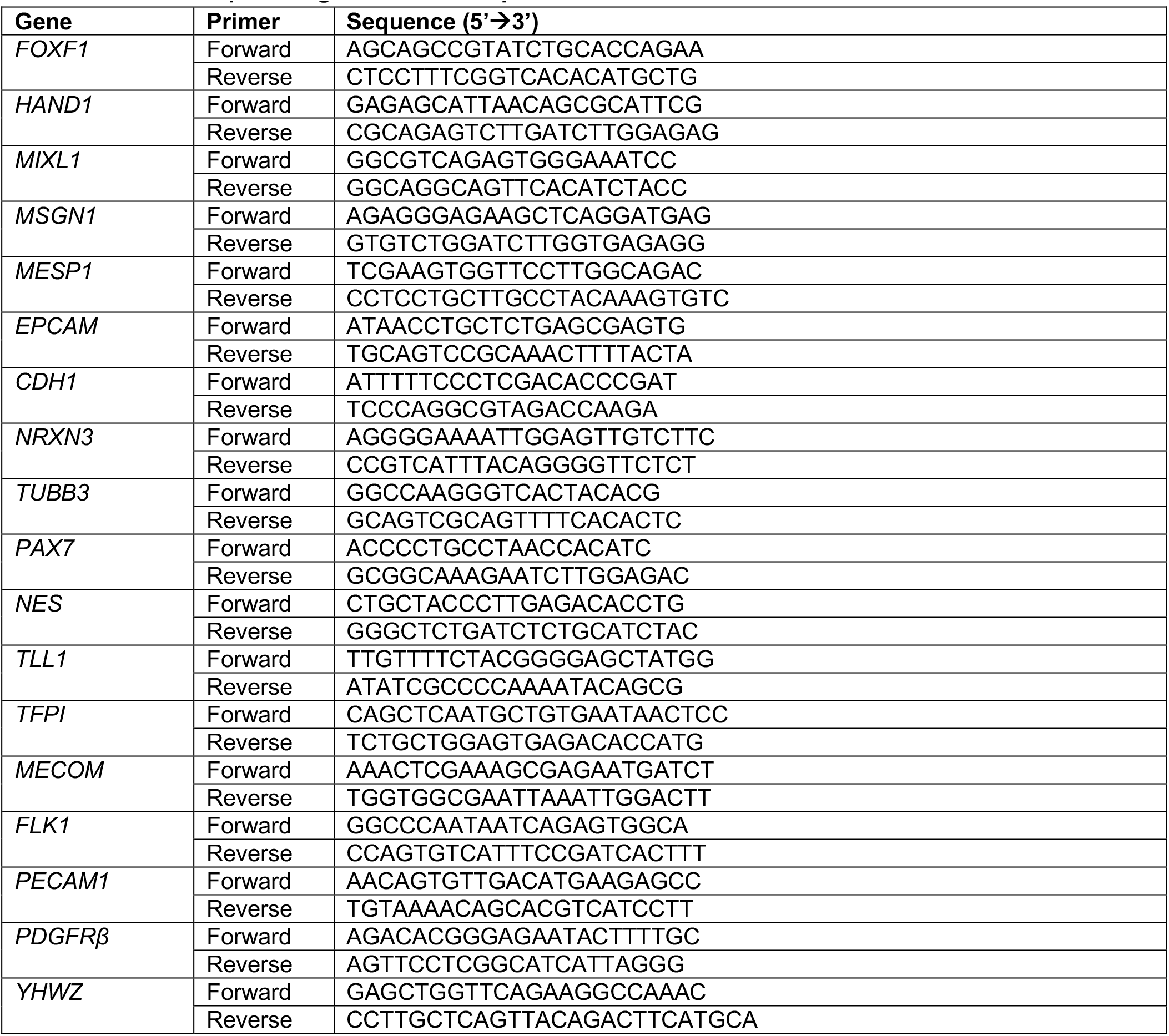
List of RT-qPCR oligonucleotide sequences.

**Table S3.**
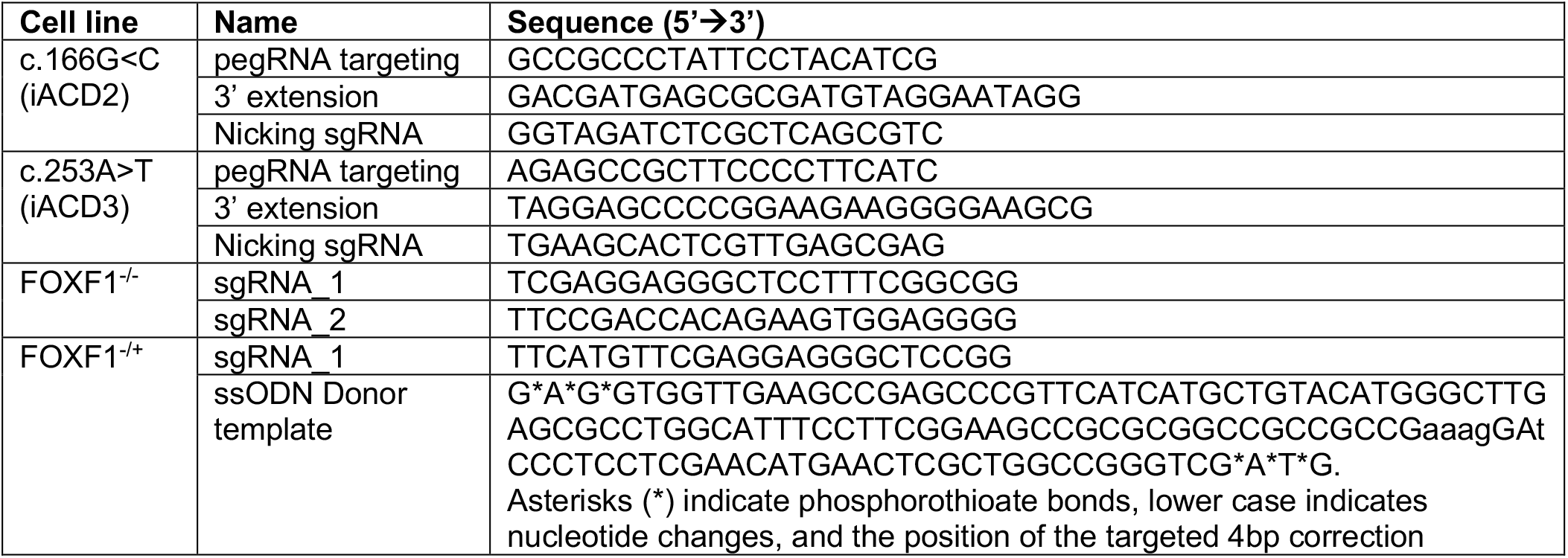
List of sgRNA and donor DNA template sequences.

**Table S4.**
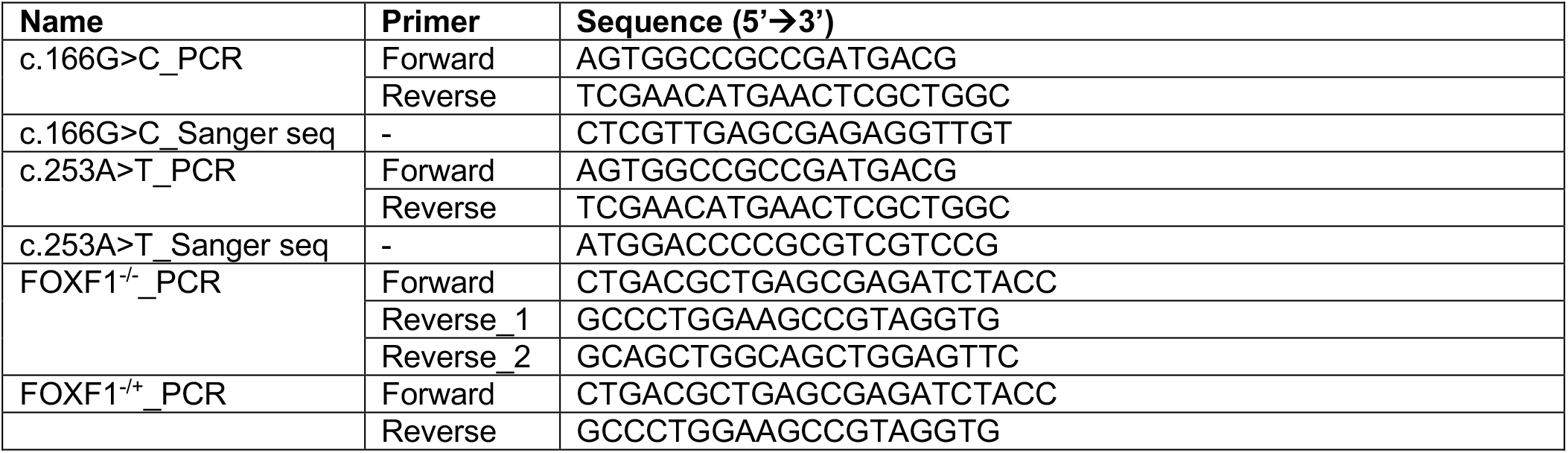
List of genotyping and sanger sequencing primers.

**Table S5.**
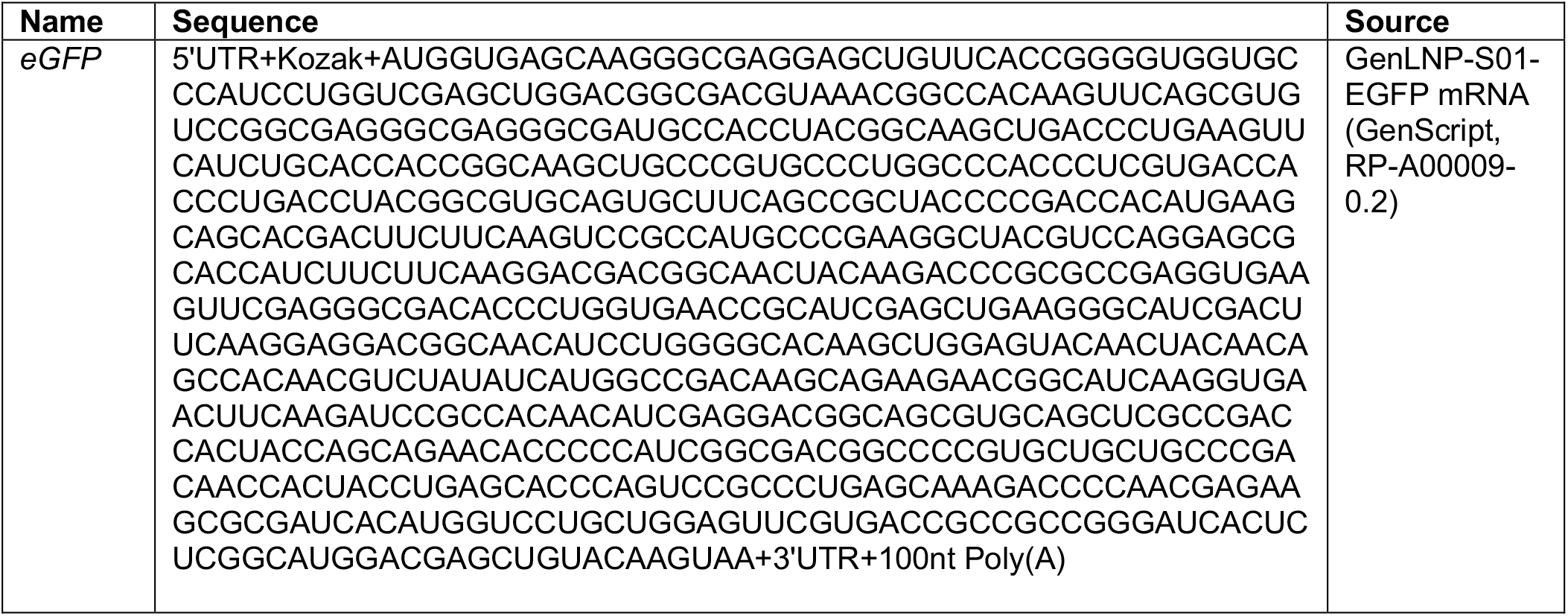
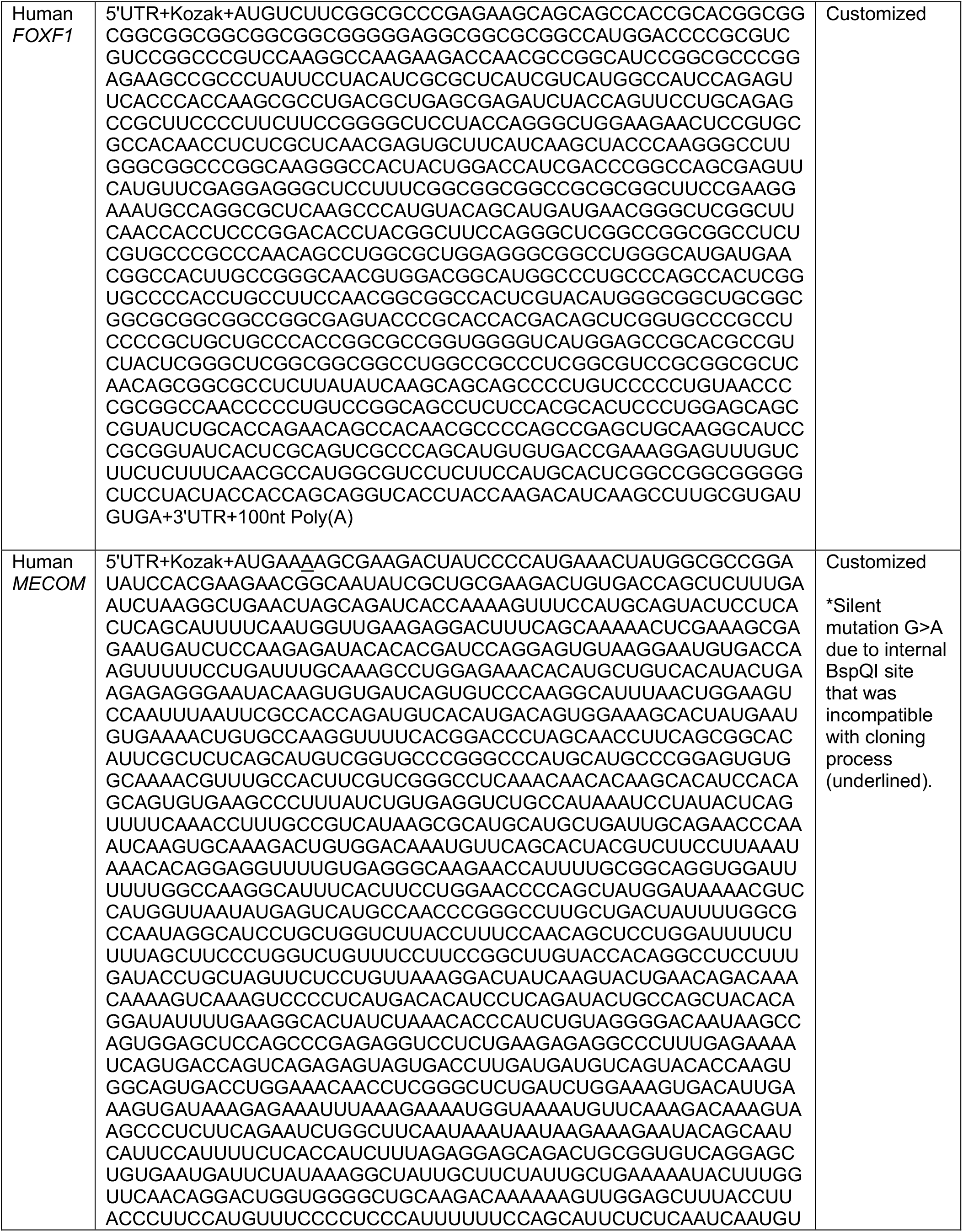
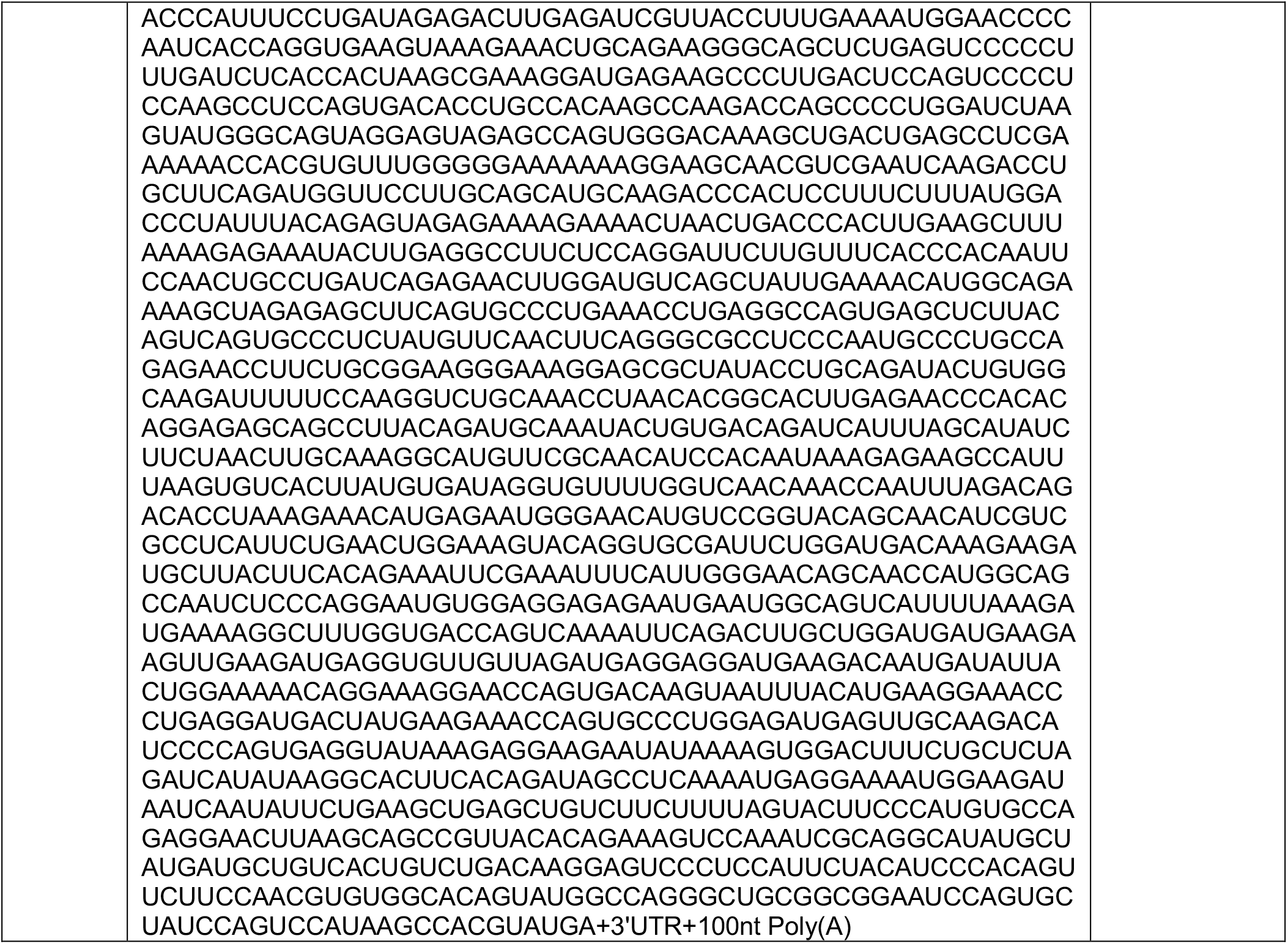
mRNA sequences for mRNA-LNPs.

## REFERENCES

1. Dharmadhikari, A.V., Szafranski, P., Kalinichenko, V.V., and Stankiewicz, P. (2015). Genomic and Epigenetic Complexity of the FOXF1 Locus in 16q24.1: Implications for Development and Disease. Curr Genomics 16, 107–116. 10.2174/1389202916666150122223252.

2. Golson, M.L., and Kaestner, K.H. (2016). Fox transcription factors: from development to disease. Development 143, 4558–4570. 10.1242/dev.112672.

3. Mahlapuu, M., Ormestad, M., Enerbäck, S., and Carlsson, P. (2001). The forkhead transcription factor Foxf1 is required for differentiation of extra-embryonic and lateral plate mesoderm. Development 128, 155–166. 10.1242/dev.128.2.155.

4. Kalinichenko, V.V., Lim, L., Stolz, D.B., Shin, B., Rausa, F.M., Clark, J., Whitsett, J.A., Watkins, S.C., and Costa, R.H. (2001). Defects in pulmonary vasculature and perinatal lung hemorrhage in mice heterozygous null for the Forkhead Box f1 transcription factor. Dev Biol 235, 489–506. 10.1006/dbio.2001.0322.

5. Lim, L., Kalinichenko, V.V., Whitsett, J.A., and Costa, R.H. (2002). Fusion of lung lobes and vessels in mouse embryos heterozygous for the forkhead box f1 targeted allele. Am J Physiol Lung Cell Mol Physiol 282, L1012–1022. 10.1152/ajplung.00371.2001.

6. Slot, E., Edel, G., Cutz, E., van Heijst, A., Post, M., Schnater, M., Wijnen, R., Tibboel, D., Rottier, R., and de Klein, A. (2018). Alveolar capillary dysplasia with misalignment of the pulmonary veins: clinical, histological, and genetic aspects. Pulm Circ 8, 2045894018795143. 10.1177/2045894018795143.

7. Sen, P., Yang, Y., Navarro, C., Silva, I., Szafranski, P., Kolodziejska, K.E., Dharmadhikari, A.V., Mostafa, H., Kozakewich, H., Kearney, D., et al. (2013). Novel FOXF1 mutations in sporadic and familial cases of alveolar capillary dysplasia with misaligned pulmonary veins imply a role for its DNA binding domain. Hum Mutat 34, 801–811. 10.1002/humu.22313.

8. Szafranski, P., Gambin, T., Dharmadhikari, A.V., Akdemir, K.C., Jhangiani, S.N., Schuette, J., Godiwala, N., Yatsenko, S.A., Sebastian, J., Madan-Khetarpal, S., et al. (2016). Pathogenetics of alveolar capillary dysplasia with misalignment of pulmonary veins. Hum Genet 135, 569–586. 10.1007/s00439-016-1655-9.

9. Bishop, N.B., Stankiewicz, P., and Steinhorn, R.H. (2011). Alveolar Capillary Dysplasia. Am J Respir Crit Care Med 184, 172–179. 10.1164/rccm.201010-1697CI.

10. Guo, M., Wikenheiser-Brokamp, K.A., Kitzmiller, J.A., Jiang, C., Wang, G., Wang, A., Preissl, S., Hou, X., Buchanan, J., Karolak, J.A., et al. (2023). Single Cell Multiomics Identifies Cells and Genetic Networks Underlying Alveolar Capillary Dysplasia. Am J Respir Crit Care Med 208, 709–725. 10.1164/rccm.202210-2015OC.

11. Alturkustani, M., Li, D., Byers, J.T., Szymanski, L., Parham, D.M., Shi, W., and Wang, L.L. (2021). Histopathologic features of alveolar capillary dysplasia with misalignment of pulmonary veins with atypical clinical presentation. Cardiovasc Pathol 50, 107289. 10.1016/j.carpath.2020.107289.

12. Reiter, J., Szafranski, P., Breuer, O., Perles, Z., Dagan, T., Stankiewicz, P., and Kerem, E. (2016). Variable phenotypic presentation of a novel FOXF1 missense mutation in a single family. Pediatric Pulmonology 51, 921–927. 10.1002/ppul.23425.

13. Abu-El-Haija, A., Fineman, J., Connolly, A.J., Murali, P., Judge, L.M., and Slavotinek, A.M. (2018). Two patients with FOXF1 mutations with alveolar capillary dysplasia with misalignment of pulmonary veins and other malformations: Two different presentations and outcomes. American Journal of Medical Genetics Part A 176, 2877–2881. 10.1002/ajmg.a.40641.

14. Pradhan, A., Dunn, A., Ustiyan, V., Bolte, C., Wang, G., Whitsett, J.A., Zhang, Y., Porollo, A., Hu, Y.-C., Xiao, R., et al. (2019). The S52F FOXF1 Mutation Inhibits STAT3 Signaling and Causes Alveolar Capillary Dysplasia. Am J Respir Crit Care Med 200, 1045–1056. 10.1164/rccm.201810-1897OC.

15. Sun, F., Wang, G., Pradhan, A., Xu, K., Gomez-Arroyo, J., Zhang, Y., Kalin, G.T., Deng, Z., Vagnozzi, R.J., He, H., et al. (2021). Nanoparticle Delivery of STAT3 Alleviates Pulmonary Hypertension in a Mouse Model of Alveolar Capillary Dysplasia. Circulation 144, 539–555. 10.1161/CIRCULATIONAHA.121.053980.

16. Edel, G.G., van Kempen, M., Munck, A.B., Huisman, C.N., Naalden, C.A.P., Brouwer, R.W.W., Koornneef, S., van IJcken, W.F.J., Wijnen, R.M.H., and Rottier, R.J. (2024). The molecular consequences of FOXF1 missense mutations associated with alveolar capillary dysplasia with misalignment of pulmonary veins. Journal of Biomedical Science 31, 100. 10.1186/s12929-024-01088-5.

17. Pradhan, A., Che, L., Ustiyan, V., Reza, A.A., Pek, N.M., Zhang, Y., Alber, A.B., Kalin, T.R., Wambach, J.A., Gu, M., et al. (2023). Novel FOXF1-Stabilizing Compound TanFe Stimulates Lung Angiogenesis in Alveolar Capillary Dysplasia. Am J Respir Crit Care Med 207, 1042–1054. 10.1164/rccm.202207-1332OC.

18. Miao, Y., Pek, N.M., Tan, C., Jiang, C., Yu, Z., Iwasawa, K., Shi, M., Kechele, D.O., Sundaram, N., Pastrana-Gomez, V., et al. (2025). Co-development of mesoderm and endoderm enables organotypic vascularization in lung and gut organoids. Cell 188, 4295–4313.e27. 10.1016/j.cell.2025.05.041.

19. Slot, E., de Klein, A., and Rottier, R.J. (2020). Generation of three iPSC lines from two patients with heterozygous FOXF1 mutations associated to Alveolar Capillary Dysplasia with Misalignment of the Pulmonary Veins. Stem Cell Res 44, 101745. 10.1016/j.scr.2020.101745.

20. Wimmer, R.A., Leopoldi, A., Aichinger, M., Wick, N., Hantusch, B., Novatchkova, M., Taubenschmid, J., Hämmerle, M., Esk, C., Bagley, J.A., et al. (2019). Human blood vessel organoids as a model of diabetic vasculopathy. Nature 565, 505–510. 10.1038/s41586-018-0858-8.

21. Wimmer, R.A., Leopoldi, A., Aichinger, M., Kerjaschki, D., and Penninger, J.M. (2019). Generation of blood vessel organoids from human pluripotent stem cells. Nat Protoc 14, 3082–3100. 10.1038/s41596-019-0213-z.

22. Voyta, J.C., Via, D.P., Butterfield, C.E., and Zetter, B.R. (1984). Identification and isolation of endothelial cells based on their increased uptake of acetylated-low density lipoprotein. J Cell Biol 99, 2034–2040. 10.1083/jcb.99.6.2034.

23. Kamp, J.C., Neubert, L., Ackermann, M., Stark, H., Plucinski, E., Shah, H.R., Janciauskiene, S., Bergmann, A.K., Schmidt, G., Welte, T., et al. (2022). A Morphomolecular Approach to Alveolar Capillary Dysplasia. The American Journal of Pathology 192, 1110–1121. 10.1016/j.ajpath.2022.05.004.

24. Ren, X., Ustiyan, V., Pradhan, A., Cai, Y., Havrilak, J.A., Bolte, C.S., Shannon, J.M., Kalin, T.V., and Kalinichenko, V.V. (2014). FOXF1 Transcription Factor Is Required for Formation of Embryonic Vasculature by Regulating VEGF Signaling in Endothelial Cells. Circulation Research 115, 709–720. 10.1161/CIRCRESAHA.115.304382.

25. Hervé, M.-A., Buteau-Lozano, H., Mourah, S., Calvo, F., and Perrot-Applanat, M. (2005). VEGF189 stimulates endothelial cells proliferation and migration in vitro and up-regulates the expression of Flk-1/KDR mRNA. Exp Cell Res 309, 24–31. 10.1016/j.yexcr.2005.05.022.

26. Cheng, J., Novati, G., Pan, J., Bycroft, C., Þemgulytë, A., Applebaum, T., Pritzel, A., Wong, L.H., Zielinski, M., Sargeant, T., et al. (2023). Accurate proteome-wide missense variant effect prediction with AlphaMissense. Science 381, eadg7492. 10.1126/science.adg7492.

27. Adzhubei, I., Jordan, D.M., and Sunyaev, S.R. (2013). Predicting Functional Effect of Human Missense Mutations Using PolyPhen-2. Curr Protoc Hum Genet 0 7, Unit7.20. 10.1002/0471142905.hg0720s76.

28. Tyser, R.C.V., Mahammadov, E., Nakanoh, S., Vallier, L., Scialdone, A., and Srinivas, S. (2021). Single-cell transcriptomic characterization of a gastrulating human embryo. Nature 600, 285–289. 10.1038/s41586-021-04158-y.

29. Chen, B., Khan, H., Yu, Z., Yao, L., Freeburne, E., Jo, K., Johnson, C., and Heemskerk, I. (2025). Extended culture of 2D gastruloids to model human mesoderm development. Nat Methods 22, 1355–1365. 10.1038/s41592-025-02669-4.

30. Moris, N., Anlas, K., van den Brink, S.C., Alemany, A., Schröder, J., Ghimire, S., Balayo, T., van Oudenaarden, A., and Martinez Arias, A. (2020). An in vitro model of early anteroposterior organization during human development. Nature 582, 410–415. 10.1038/s41586-020-2383-9.

31. Chappell, J.C., Mouillesseaux, K.P., and Bautch, V.L. (2013). Flt-1 (Vascular Endothelial Growth Factor Receptor-1) Is Essential for the Vascular Endothelial Growth Factor–Notch Feedback Loop During Angiogenesis. Arteriosclerosis, Thrombosis, and Vascular Biology 33, 1952–1959. 10.1161/ATVBAHA.113.301805.

32. Nikolova, M.T., He, Z., Seimiya, M., Jonsson, G., Cao, W., Okuda, R., Wimmer, R.A., Okamoto, R., Nikoloff, J.M., Dittrich, P.S., et al. (2025). Fate and state transitions during human blood vessel organoid development. Cell 188, 3329–3348.e31. 10.1016/j.cell.2025.03.037.

33. Lv, J., Meng, S., Gu, Q., Zheng, R., Gao, X., Kim, J., Chen, M., Xia, B., Zuo, Y., Zhu, S., et al. (2023). Epigenetic landscape reveals MECOM as an endothelial lineage regulator. Nat Commun 14, 2390. 10.1038/s41467-023-38002-w.

34. Bolte, C., Flood, H.M., Ren, X., Jagannathan, S., Barski, A., Kalin, T.V., and Kalinichenko, V.V. (2017). FOXF1 transcription factor promotes lung regeneration after partial pneumonectomy. Sci Rep 7, 10690. 10.1038/s41598-017-11175-3.

35. Armulik, A., Abramsson, A., and Betsholtz, C. (2005). Endothelial/pericyte interactions. Circ Res 97, 512–523. 10.1161/01.RES.0000182903.16652.d7.

36. Jin, S., Plikus, M.V., and Nie, Q. (2025). CellChat for systematic analysis of cell-cell communication from single-cell transcriptomics. Nat Protoc 20, 180–219. 10.1038/s41596-024-01045-4.

37. Kalinichenko, V.V., Gusarova, G.A., Kim, I.-M., Shin, B., Yoder, H.M., Clark, J., Sapozhnikov, A.M., Whitsett, J.A., and Costa, R.H. (2004). Foxf1 haploinsufficiency reduces Notch-2 signaling during mouse lung development. Am J Physiol Lung Cell Mol Physiol 286, L521–530. 10.1152/ajplung.00212.2003.

38. Wang, G., Wen, B., Guo, M., Li, E., Zhang, Y., Whitsett, J.A., Kalin, T.V., and Kalinichenko, V.V. (2024). Identification of endothelial and mesenchymal FOXF1 enhancers involved in alveolar capillary dysplasia. Nat Commun 15, 5233. 10.1038/s41467-024-49477-6.

39. Wang, G., Wen, B., Deng, Z., Zhang, Y., Kolesnichenko, O.A., Ustiyan, V., Pradhan, A., Kalin, T.V., and Kalinichenko, V.V. (2022). Endothelial progenitor cells stimulate neonatal lung angiogenesis through FOXF1-mediated activation of BMP9/ACVRL1 signaling. Nat Commun 13, 2080. 10.1038/s41467-022-29746-y.

40. Hoggatt, A.M., Kim, J.-R., Ustiyan, V., Ren, X., Kalin, T.V., Kalinichenko, V.V., and Herring, B.P. (2013). The Transcription Factor Foxf1 Binds to Serum Response Factor and Myocardin to Regulate Gene Transcription in Visceral Smooth Muscle Cells. J Biol Chem 288, 28477–28487. 10.1074/jbc.M113.478974.

41. Stankiewicz, P., Sen, P., Bhatt, S.S., Storer, M., Xia, Z., Bejjani, B.A., Ou, Z., Wiszniewska, J., Driscoll, D.J., Bolivar, J., et al. (2009). Genomic and Genic Deletions of the FOX Gene Cluster on 16q24.1 and Inactivating Mutations of *FOXF1* Cause Alveolar Capillary Dysplasia and Other Malformations. The American Journal of Human Genetics 84, 780–791. 10.1016/j.ajhg.2009.05.005.

42. Sadahiro, T., Isomi, M., Muraoka, N., Kojima, H., Haginiwa, S., Kurotsu, S., Tamura, F., Tani, H., Tohyama, S., Fujita, J., et al. (2018). Tbx6 Induces Nascent Mesoderm from Pluripotent Stem Cells and Temporally Controls Cardiac versus Somite Lineage Diversification. Cell Stem Cell 23, 382–395.e5. 10.1016/j.stem.2018.07.001.

43. Rajan, A.M., Ma, R.C., Kocha, K.M., Zhang, D.J., and Huang, P. (2020). Dual function of perivascular fibroblasts in vascular stabilization in zebrafish. PLOS Genetics 16, e1008800. 10.1371/journal.pgen.1008800.

44. Davis, G.E., and Senger, D.R. (2005). Endothelial extracellular matrix: biosynthesis, remodeling, and functions during vascular morphogenesis and neovessel stabilization. Circ Res 97, 1093–1107. 10.1161/01.RES.0000191547.64391.e3.

45. Sweeney, M., and Foldes, G. (2018). It Takes Two: Endothelial-Perivascular Cell Cross-Talk in Vascular Development and Disease. Front. Cardiovasc. Med. 5. 10.3389/fcvm.2018.00154.

46. Yazbeck, P., Cullere, X., Bennett, P., Yajnik, V., Wang, H., Kawada, K., Davis, V., Parikh, A., Kuo, A., Mysore, V., et al. (2022). DOCK4 regulation of Rho GTPases mediates pulmonary vascular barrier function. Arterioscler Thromb Vasc Biol 42, 886–902. 10.1161/ATVBAHA.122.317565.

47. Abraham, S., Scarcia, M., Bagshaw, R.D., McMahon, K., Grant, G., Harvey, T., Yeo, M., Esteves, F.O.G., Thygesen, H.H., Jones, P.F., et al. (2015). A Rac/Cdc42 exchange factor complex promotes formation of lateral filopodia and blood vessel lumen morphogenesis. Nat Commun 6, 7286. 10.1038/ncomms8286.

48. Michaelis, U.R. (2014). Mechanisms of endothelial cell migration. Cell Mol Life Sci 71, 4131–4148. 10.1007/s00018-014-1678-0.

49. Malin, D., Kim, I.-M., Boetticher, E., Kalin, T.V., Ramakrishna, S., Meliton, L., Ustiyan, V., Zhu, X., and Kalinichenko, V.V. (2007). Forkhead box F1 is essential for migration of mesenchymal cells and directly induces integrin-beta3 expression. Mol Cell Biol 27, 2486–2498. 10.1128/MCB.01736-06.

50. Towe, C.T., White, F.V., Grady, R.M., Sweet, S.C., Eghtesady, P., Wegner, D.J., Sen, P., Szafranski, P., Stankiewicz, P., Hamvas, A., et al. (2018). Infants with Atypical Presentations of Alveolar Capillary Dysplasia with Misalignment of the Pulmonary Veins Who Underwent Bilateral Lung Transplantation. The Journal of Pediatrics 194, 158–164.e1. 10.1016/j.jpeds.2017.10.026.

51. Nakajima, D., Oda, H., Mineura, K., Goto, T., Kato, I., Baba, S., Ikeda, T., Chen-Yoshikawa, T.F., and Date, H. (2020). Living-donor single-lobe lung transplantation for pulmonary hypertension due to alveolar capillary dysplasia with misalignment of pulmonary veins. Am J Transplant 20, 1739–1743. 10.1111/ajt.15762.

52. Dharmadhikari, A.V., Sun, J.J., Gogolewski, K., Carofino, B.L., Ustiyan, V., Hill, M., Majewski, T., Szafranski, P., Justice, M.J., Ray, R.S., et al. (2016). Lethal lung hypoplasia and vascular defects in mice with conditional Foxf1 overexpression. Biol Open 5, 1595–1606. 10.1242/bio.019208.

53. Deng, Z., Gao, W., Lan, Y.-W., Do, J., Kalin, T., and Kalinichenko, V. (2025). Nanoparticle Delivery of FOXF1-mRNA Protects Pulmonary Endothelial Cells From Injury in Neonatal Sepsis Model. Am J Respir Crit Care Med 211, A7705–A7705. 10.1164/ajrccm.2025.211.Abstracts.A7705.

54. Kohram, F., Deng, Z., Zhang, Y., Al Reza, A., Li, E., Kolesnichenko, O.A., Shukla, S., Ustiyan, V., Gomez-Arroyo, J., Acharya, A., et al. (2023). Demonstration of Safety in Wild Type Mice of npFOXF1, a Novel Nanoparticle-Based Gene Therapy for Alveolar Capillary Dysplasia with Misaligned Pulmonary Veins. Biologics 17, 43–55. 10.2147/BTT.S400006.

55. Zhang, Y., Tan, C., Liu, Z., Mao, X., Jiang, C., Mohammed, A.N., Li, X., Lu, R., Wang, A., Maihemuti, W., et al. (2026). Rebalancing NTRK2 isoforms promotes vascular regeneration in bronchopulmonary dysplasia. Cell Stem Cell 33, 125–141.e11. 10.1016/j.stem.2025.12.006.

56. Mattar, C.N.Z., Chew, W.L., and Lai, P.S. (2024). Embryo and fetal gene editing: Technical challenges and progress toward clinical applications. Molecular Therapy Methods & Clinical Development 32. 10.1016/j.omtm.2024.101229.

57. Jerabek, S., Kim, J., Sung, J., Jung, C., Kulmann, M.I.R., Isado, M., Jang, H.-S., Li, M., Bhatele, S., Kappy, M., et al. (2026). Efficient base editing and development in human embryos without chromosomal alterations. Preprint at bioRxiv, 10.64898/2026.05.30.728989 10.64898/2026.05.30.728989.

58. Riley, R.S., Kashyap, M.V., Billingsley, M.M., White, B., Alameh, M.-G., Bose, S.K., Zoltick, P.W., Li, H., Zhang, R., Cheng, A.Y., et al. (2021). Ionizable lipid nanoparticles for in utero mRNA delivery. Science Advances 7, eaba1028. 10.1126/sciadv.aba1028.

59. Gillich, A., Zhang, F., Farmer, C.G., Travaglini, K.J., Tan, S.Y., Gu, M., Zhou, B., Feinstein, J.A., Krasnow, M.A., and Metzger, R.J. (2020). Capillary cell-type specialization in the alveolus. Nature 586, 785–789. 10.1038/s41586-020-2822-7.

60. Paik, D.T., Tian, L., Williams, I.M., Rhee, S., Zhang, H., Liu, C., Mishra, R., Wu, S.M., Red-Horse, K., and Wu, J.C. (2020). Single-cell RNA-seq Unveils Unique Transcriptomic Signatures of Organ-Specific Endothelial Cells. Circulation 142, 1848–1862. 10.1161/CIRCULATIONAHA.119.041433.

61. Poling, H.M., Sundaram, N., Fisher, G.W., Singh, A., Shiley, J.R., Nattamai, K., Govindarajah, V., Cortez, A.R., Krutko, M.O., Ménoret, S., et al. (2024). Human pluripotent stem cell-derived organoids repair damaged bowel in vivo. Cell Stem Cell 31, 1513–1523.e7. 10.1016/j.stem.2024.08.009.

62. Wu, Y., Sidharta, M., Zhong, A., Persily, B., Li, M., and Zhou, T. (2023). Protocol for the design, conduct, and evaluation of prime editing in human pluripotent stem cells. STAR Protoc 4, 102583. 10.1016/j.xpro.2023.102583.

63. Wu, Y., Zhong, A., Sidharta, M., Kim, T.W., Ramirez, B., Persily, B., Studer, L., and Zhou, T. (2024). Robust and inducible genome editing via an all-in-one prime editor in human pluripotent stem cells. Nat Commun 15, 10824. 10.1038/s41467-024-55104-1.

64. Abramson, J., Adler, J., Dunger, J., Evans, R., Green, T., Pritzel, A., Ronneberger, O., Willmore, L., Ballard, A.J., Bambrick, J., et al. (2024). Accurate structure prediction of biomolecular interactions with AlphaFold 3. Nature 630, 493–500. 10.1038/s41586-024-07487-w.

65. Humphrey, W., Dalke, A., and Schulten, K. (1996). VMD: visual molecular dynamics. J Mol Graph 14, 33–38, 27–28. 10.1016/0263-7855(96)00018-5.

66. Clark, K.L., Halay, E.D., Lai, E., and Burley, S.K. (1993). Co-crystal structure of the HNF-3/fork head DNA-recognition motif resembles histone H5. Nature 364, 412–420. 10.1038/364412a0.

67. Van Der Spoel, D., Lindahl, E., Hess, B., Groenhof, G., Mark, A.E., and Berendsen, H.J.C. (2005). GROMACS: fast, flexible, and free. J Comput Chem 26, 1701–1718. 10.1002/jcc.20291.

68. Abraham, M.J., Murtola, T., Schulz, R., Páll, S., Smith, J.C., Hess, B., and Lindahl, E. (2015). GROMACS: High performance molecular simulations through multi-level parallelism from laptops to supercomputers. SoftwareX 1-2, 19–25. 10.1016/j.softx.2015.06.001.

69. Best, R.B., Zhu, X., Shim, J., Lopes, P.E.M., Mittal, J., Feig, M., and Mackerell, A.D. (2012). Optimization of the additive CHARMM all-atom protein force field targeting improved sampling of the backbone ö, ø and side-chain ÷(1) and ÷(2) dihedral angles. J Chem Theory Comput 8, 3257–3273. 10.1021/ct300400x.

70. Bessonett, S., Perry, G.A., Paul.gabriel, Luo, D., Grassmann, J., Flynn, W.F., Courtois, E., Robson, P., and Daigle, S.L. (2022). JAX - EZ Lysis Nuclei Isolation for 10x Genomics Assays.

71. Satpathy, A.T., Granja, J.M., Yost, K.E., Qi, Y., Meschi, F., McDermott, G.P., Olsen, B.N., Mumbach, M.R., Pierce, S.E., Corces, M.R., et al. (2019). Massively parallel single-cell chromatin landscapes of human immune cell development and intratumoral T cell exhaustion. Nat Biotechnol 37, 925–936. 10.1038/s41587-019-0206-z.

72. Hao, Y., Stuart, T., Kowalski, M.H., Choudhary, S., Hoffman, P., Hartman, A., Srivastava, A., Molla, G., Madad, S., Fernandez-Granda, C., et al. (2024). Dictionary learning for integrative, multimodal and scalable single-cell analysis. Nat Biotechnol 42, 293–304. 10.1038/s41587-023-01767-y.

73. Stuart, T., Srivastava, A., Madad, S., Lareau, C.A., and Satija, R. (2021). Single-cell chromatin state analysis with Signac. Nat Methods 18, 1333–1341. 10.1038/s41592-021-01282-5.

74. Young, M.D., and Behjati, S. (2020). SoupX removes ambient RNA contamination from droplet-based single-cell RNA sequencing data. Gigascience 9, giaa151. 10.1093/gigascience/giaa151.

75. McGinnis, C.S., Murrow, L.M., and Gartner, Z.J. (2019). DoubletFinder: Doublet Detection in Single-Cell RNA Sequencing Data Using Artificial Nearest Neighbors. Cell Syst 8, 329–337.e4. 10.1016/j.cels.2019.03.003.

76. Finak, G., McDavid, A., Yajima, M., Deng, J., Gersuk, V., Shalek, A.K., Slichter, C.K., Miller, H.W., McElrath, M.J., Prlic, M., et al. (2015). MAST: a flexible statistical framework for assessing transcriptional changes and characterizing heterogeneity in single-cell RNA sequencing data. Genome Biol 16, 278. 10.1186/s13059-015-0844-5.

77. Grant, C.E., Bailey, T.L., and Noble, W.S. (2011). FIMO: scanning for occurrences of a given motif. Bioinformatics 27, 1017–1018. 10.1093/bioinformatics/btr064.

78. Andreatta, M., and Carmona, S.J. (2026). UCell and pyUCell: single-cell gene signature scoring for R and Python. Bioinformatics 42, btag055. 10.1093/bioinformatics/btag055.

79. Gu, M., Shao, N.-Y., Sa, S., Li, D., Termglinchan, V., Ameen, M., Karakikes, I., Sosa, G., Grubert, F., Lee, J., et al. (2017). Patient-Specific iPSC-Derived Endothelial Cells Uncover Pathways that Protect against Pulmonary Hypertension in BMPR2 Mutation Carriers. Cell Stem Cell 20, 490–504.e5. 10.1016/j.stem.2016.08.019.

80. Liu, Z., Liu, Y., Yu, Z., Tan, C., Pek, N., O’Donnell, A., Wu, A., Glass, I., Winlaw, D.S., Guo, M., et al. (2024). APOE–NOTCH axis governs elastogenesis during human cardiac valve remodeling. Nat Cardiovasc Res 3, 933–950. 10.1038/s44161-024-00510-3.

81. Suarez-Arnedo, A., Torres Figueroa, F., Clavijo, C., Arbeláez, P., Cruz, J.C., and Muñoz-Camargo, C. (2020). An image J plugin for the high throughput image analysis of in vitro scratch wound healing assays. PLoS One 15, e0232565. 10.1371/journal.pone.0232565.

82. Xie, Z., Bailey, A., Kuleshov, M.V., Clarke, D.J.B., Evangelista, J.E., Jenkins, S.L., Lachmann, A., Wojciechowicz, M.L., Kropiwnicki, E., Jagodnik, K.M., et al. (2021). Gene Set Knowledge Discovery with Enrichr. Current Protocols 1, e90. 10.1002/cpz1.90.

83. Tang, D., Chen, M., Huang, X., Zhang, G., Zeng, L., Zhang, G., Wu, S., and Wang, Y. (2023). SRplot: A free online platform for data visualization and graphing. PLOS ONE 18, e0294236. 10.1371/journal.pone.0294236.

84. McGibbon, R.T., Beauchamp, K.A., Harrigan, M.P., Klein, C., Swails, J.M., Hernández, C.X., Schwantes, C.R., Wang, L.-P., Lane, T.J., and Pande, V.S. (2015). MDTraj: A Modern Open Library for the Analysis of Molecular Dynamics Trajectories. Biophys J 109, 1528–1532. 10.1016/j.bpj.2015.08.015.

85. Shrake, A., and Rupley, J.A. (1973). Environment and exposure to solvent of protein atoms. Lysozyme and insulin. J Mol Biol 79, 351–371. 10.1016/0022-2836(73)90011-9.

86. Gvritishvili, A.G., Gribenko, A.V., and Makhatadze, G.I. (2008). Cooperativity of complex salt bridges. Protein Sci 17, 1285–1290. 10.1110/ps.034975.108.

